# Energy Conservation Drives Adaptive Specialization Through Fine-tuned Organizations

**DOI:** 10.64898/2025.12.06.692764

**Authors:** Teresa Femenia, Naeimeh Atabaki-Pasdar, Suvi Linna-Kuosmanen, Kevin Grove, Manolis Kellis, Leandro Z. Agudelo

## Abstract

Energy conservation is at the basis of evolutionary optimization in living systems. Yet, how cells maintain function during metabolic stress while preserving adaptive capacity remains unclear. Here, we use a functional genomics approach to identify metabolic human accelerated regions (mHARs) genomic hubs (evolutionarily tuned regions) in adipose tissue that coordinate adaptive responses to nutrient deficit. During fasting, cooperative regulators PGC1A, YY1, and CTCF activate these hubs, promoting chromatin contacts that regulate lipid cycling genes such as Dgat1 and Atg4c. This activation maintains lipid and pH homeostasis while limiting oxidative stress, thereby preserving cellular plasticity. When disrupted, nutrient-stress adaptations are impaired leading to metabolic dysfunction and glucose intolerance. This is driven by NCOA7, an oxidative stress mediator that impairs lysosome function and transcriptional plasticity, triggering an inflammatory response. In cellular models, this mechanism blocks pH-mediated PGC1A droplet plasticity, further inhibiting mHAR-genomic hub activity and metabolic homeostasis. Human adipocytes show enhanced transcriptional activity of these genomic hubs compared to murine cells, with multi-enhancer mHAR regulation suppressing disease-associated variants linked to ANGPTL3 and PCSK9. Loss of this cooperative plasticity shifts cells to maladaptive responses, activating STAT1-driven disease gene expression. Our findings reveal that investigating the contextual activity of evolutionary adaptations can uncover functional hubs critical for stress resilience. This also shows that cells exploit nutrient fluctuations to drive tunable transcriptional organizations that lead to durable adaptations, identifying actionable targets for metabolic disease.

**One-Sentence Summary:** Evolutionary tuned nuclear compartments convert nutrient stress into adaptive transcriptional organizations that preserve plasticity and suppress metabolic disease.

## Introduction

Metabolic diseases such as obesity and type 2 diabetes affect millions globally. These conditions are characterized by a decline in metabolic plasticity, for instance, impaired nutrient sensing or inadequate response to food scarcity. This plasticity is not only crucial for maintaining cellular and organismal homeostasis but also for reducing the risk of associated comorbidities.

Metabolic plasticity is essential throughout evolution, optimizing energy use for survival, growth, and reproduction (*1*, *2*), and influencing the diversity of metabolic traits, such as energy expenditure, fat mass, body size, and reproductive abilities (*3–6*). For example, large primates show lineage-specific adaptations such as reduced energy expenditure and increased fat mass (*3*, *5*, *6*), which has been suggested to impact foraging, cognition, and metabolic endurance (*3–6*). The mobilization of nutrients is also essential for energy allocation and is influenced by the environment, diet, and genetics (*1*, *2*). Despite variations among species, all organisms need to adjust to limited nutrients, indicating the existence of pan-molecular signatures that are constrained yet subject to evolutionary fine-tuning (*7*).

One example of genomic fine-tuning includes the human accelerated regions (HARs), conserved non-coding elements with increased rates of human-specific mutations (*8*, *9*). Many HARs function as transcriptional enhancers with human-specific activity during embryonic development (*8*, *9*). Despite the critical roles of energy conservation in evolution and disease, it remains unclear how energetic states influence the activity of fine-tuned genomic regions linked to metabolic plasticity.

This question becomes more complex when we consider how genes are organized and dynamically regulated in 3D space. Dynamic genome organization plays a crucial role in transcriptional plasticity and response to environmental stimuli (*10*). Transcriptional compartments, nuclear organizations formed during transcription, facilitate the acquisition of cell-specific functions (*10*). These compartments occupy distinct chromosome territories, influencing interactions with nearby transcription factories and thus enabling gene coregulation (*11*, *12*). In many organisms, transcriptional features are conserved, exhibiting dynamic plasticity and stochasticity, which are all hallmarks of self-organized systems (*11*). Transcriptional organization is influenced by intrinsically disordered regions (IDRs) in transcriptional regulators (*13*, *14*), liquid-like phase separation (LLPS), and genome structural features (*13*), which facilitates the formation of multimodal nuclear compartments regulating distinct gene programs (*11*, *13*, *15–17*).

Environmental variation itself influences compartment formation and condensate properties (*18*, *19*). Cold and fasting conditions can reduce cytosolic volume, promoting molecular interactions, condensate formation (*20–23*), and modulating droplet properties for cellular fitness (*18*). Moreover, evolution fine-tunes molecular adaptations for efficient energy conservation by selecting residues favoring multimolecular assemblies (*18*, *24*), introducing mutations enhancing nutrient conservation (*25*), incorporating lineage- and trait-specific genome substitutions (*9*), and refining chromatin interactions (*26*). Despite these insights, it is not clear how energetic states are associated with fine-tuned genomic regions influencing transcriptional compartment properties.

We aim to identify common ground between these evolutionary and structural perspectives, by focusing on transcriptional adaptations that help maintain cellular homeostasis while responding to environmental changes (e.g., exercise, cold, and fasting (*27–29*)). For example, PGC1A transcriptional coactivators regulate gene programs for metabolic homeostasis by interacting with transcription factors (*30*) and forming nuclear condensates (*31*). In brown and beige adipocytes, cold-induced PGC1A coactivators regulate thermogenesis (*28*, *32*), which can help counteract elevated energy intake (*33*). In nutrient limitation, different adaptive processes are involved, such as lipid cycling, which involves the breakdown and re-esterification of fatty acids (*34–36*). This process, activated during fasting and cold exposure, helps prevent endoplasmic reticulum (ER) stress, mitochondrial dysfunction, and lipotoxicity (*36–40*). Yet, how lipid cycling influences transcriptional plasticity and long-term adaptations remains not fully understood.

In this study, we use a functional genomics approach to explore the molecular links between metabolic plasticity, disease, and evolution. We perform this in three integrated stages: First, comparative analyses between metabolic traits in mammals identify key relationships with adipose tissue mass and endurance to food scarcity. We then investigate lineage-specific genome adaptations to identify key functional transcriptional compartment hubs influencing cellular responses to energy deficit. Finally, we test whether these regions are active during nutrient stress and examine their species-specific features. We find that, in fasted adipocytes, distinct genomic regions enriched in human-specific mutations serve as central hubs for gene regulation through chromatin interactions across distant loci, and co-regulation of genes maintaining transcriptional plasticity. We show that these regions exhibit species-specific activity in cellular models and contribute to resilience against nutrient stress and metabolic disease in organismal models. Overall, our findings reveal a link between specific transcriptional organizations and lineage-specific variations during nutrient stress, identifying functional hubs with therapeutic potential for metabolic diseases.

## Results

### 1. Relationships between metabolic traits and energy allocation in mammals

To identify links between energy use and allocation in mammals, we re-evaluated metabolic trait parameters such as daily energy expenditure, body weight, and fat mass. Recent studies have shown that humans exhibit the highest body fat composition, which has been suggested to contribute to meeting energetic demands (*3–6*). It has also been shown that low energy expenditure and larger body size are associated with greater endurance to food scarcity (*4*, *41*). Thus, our goal was to find links between metabolic traits and fat-associated metabolic parameters in mammals, followed by identifying genomic regions linked to lineage-specific adaptations and metabolic function (**fig. S1A**).

Previous studies used the amount of energy stored in fat mass mainly by allometric scaling (10% of body weight) and basal metabolic rate to estimate how long mammals can survive with limited food intake (*4*). This theoretical parameter of fasting endurance score or survival time is obtained by assessing the ratio between energy content in fat mass and energy use in basal metabolic rate (*4*).

To examine this closely, we employed a similar approach using recently published data including experimentally assessed fat mass and energy use based on total energy expenditure (*3–6*) (TEE, **fig. S1A,** Table **S1A-D**). Our analysis showed that, compared to allometrically scaled fat mass (10% of body weight), experimentally assessed fat mass has a stronger association with this endurance parameter in mammals (**Fig. 1A**). These findings underscore the significance of energy storage and use in mammals, and in particular hominids, where cognitive demands and the energetic costs of long-distance hunting and foraging have shaped endurance capabilities (*5*, *42*). These relationships suggest that metabolic plasticity is subject to evolutionary optimization under specific constraints.

**Fig. 1.**
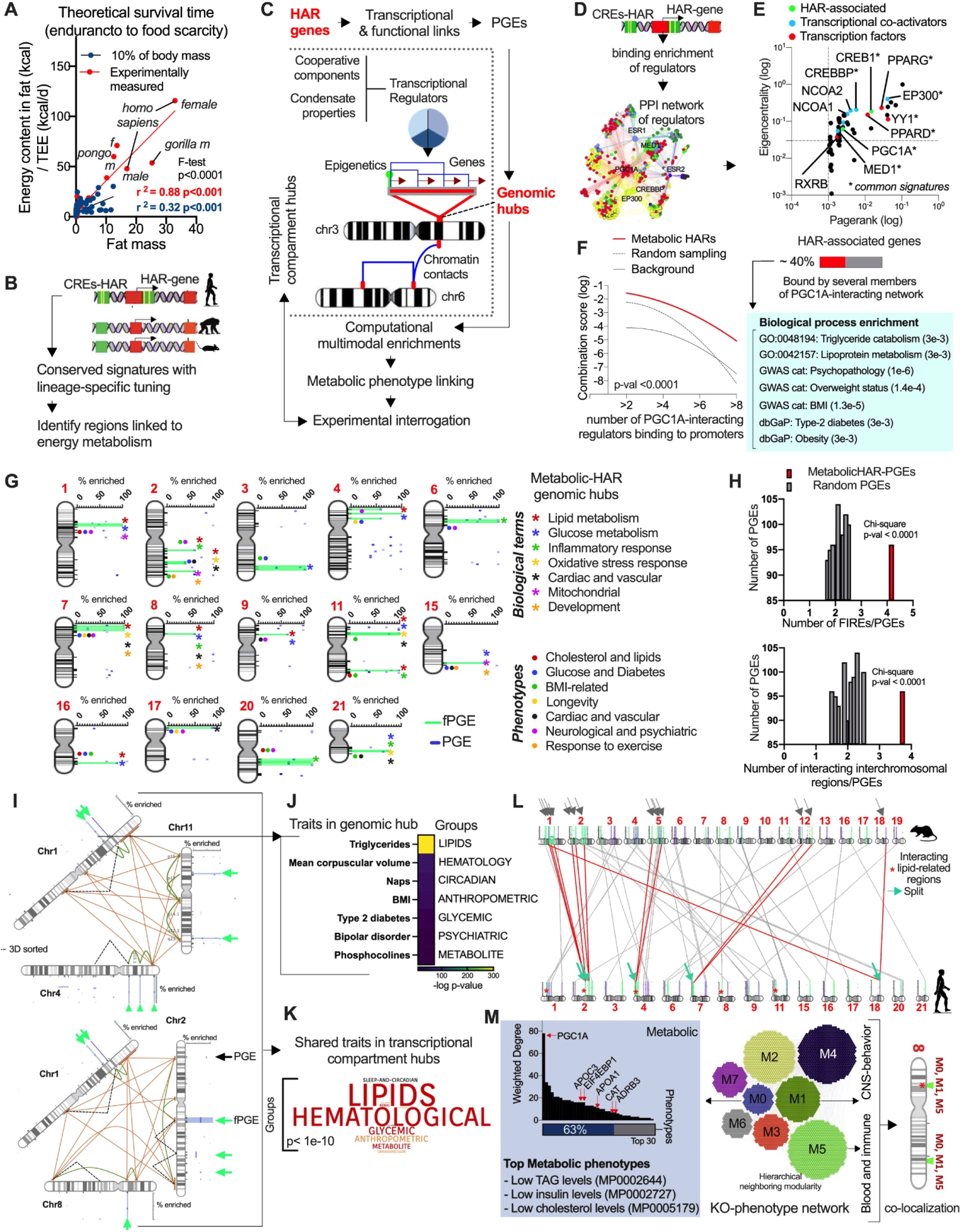
Functional genomics reveals metabolic acceleration hubs and associated nuclear compartments. **(A)** Theoretical survival time (fasting endurance score) for allometrically scaled fat mass (10% of body weight) and experimentally measured fat mass across mammals (previous reports (*5*, *6*)) and linear regression comparison (total energy expenditure, TEE). **(B)** Illustration of conserved regulatory regions with lineage-specific tuning such as human accelerated regions (HAR) and identification of HAR genomic regions associated with energy metabolism. **(C)** Illustration of HAR-associated transcriptional compartment hubs using positional gene enrichments (PGEs) of HAR-genes sharing functional enrichments, followed by interrogation of further enrichments across HAR-genomic hubs. **(D)** HAR elements and associated genes were used to identify transcriptional regulators (TRs) by binding enrichment, followed by network reconstruction of TRs using protein-protein interactions. **(E)** Network-based ranking of interacting transcriptional regulators binding HAR associated genes. In blue or red transcriptional regulators with similar function, and in green regulators directly associated with HAR elements (loci near to active HAR). **(F)** HAR genes bound (ChIP-seq binding enrichment) by >2 interacting PGC1A coregulators (PPIs filtered), termed metabolic HAR genes (red line). Combination score was assessed by dividing the number of bound transcriptional regulator (TR) combinations (TR_1_, TR_2_, …, TR_i_) by the number of co-regulated genes for each class (2TRs, 4TRs, 6TRs, 8TRs). Enrichment tests were performed, comparing the number of combinations for PGC1A-cooperative regulators binding HAR genes, and the same regulators binding similar numbers of random gene sets. The right panel shows biological enrichments for filtered HAR genes bound by >2 cooperative regulators. **(G)** PGEs for metabolic-HAR genes. Bio-enriched (fPGEs, hubs with functional enrichments) and non-enriched hubs (PGEs hubs). Biological and phenotype terms associated with each genomic hub. Arrow shows next steps to evaluate genome structural features. **(H)** Colocalized frequently-interacting regions (FIREs, defined from HiC compendia) with mHAR genomic hubs (mHAR PGEs) and random gene set PGE permutations. Lower panel, colocalization of interchromosomal regions from hESC-HiC data with mHAR genomic hubs. Chi-squared tests comparing mHAR with random hubs. **(I)** Reconstructed metabolic mHAR nuclear compartments displaying selected hubs (fPGEs in green). Orange and green lines represent inter-and intra-chromosomal contacts, respectively (from HiC data). Dotted line shows 3D imputed cis-contacts. **(J)** Regional genetic associations for mHAR genomic hub (Chr1:61-65 mb range) showing top enriched traits within trait-group class. **(K)** Regional genetic associations for mHAR genomic hubs within compartments, displayed as word-cloud based on the number of trait-group repetitions. **(L)** Murine chromosomal synteny of human mHAR genomic hubs. Green and gray arrows show split genomic hubs in mice. Red asterisk shows mHAR hubs with similar functional enrichments and with nuclear proximity in human HiC data. **(M)** Left panel, mHAR gene knockout and phenotype association network displayed as a hierarchical modular layout, displaying main phenotypic classes. Middle panel shows node-degree network analysis of the metabolic module. Arrows show mHAR hub genes sharing nuclear proximity in human HiC data. Right panel shows positional gene enrichments for genes within interacting modules. Hypergeometric test enrichments and Chi-square tests were used to determine the statistical significance between expected and observed frequencies, and network-based features for filtering associations.

Having identified potential evolutionary pressures on energy allocation, we next sought to understand the genomic basis for such adaptations. Conserved genomic regions linked to lineage-specific adaptations and similar phenotypic traits remain not fully understood. Therefore, investigating the functional impact of these DNA regions along with their response to metabolic stressors, remains an essential task (**Fig. 1B**, **fig. S1A**).

### 2. A functional genomics approach to investigate regions with human-specific mutations

To move from comparative physiological analysis to molecular mechanisms, we needed a systematic framework that both identifies and validates functional genomic regions. Understanding the relationship between gene activity and cell phenotypes requires integrating and measuring multimodal components related to transcriptional adaptation. This involves assessing gene programs within transcriptionally active genomic regions, genome organization, epigenetic modifications, DNA binding and interactions between transcriptional regulators, and transcriptional condensate plasticity (*11*, *13*).

Different approaches, from microscopy to profiling, have revealed that transcription occurs within active genomic regions known as transcriptional hubs, guiding 3D genome organization (*12*, *43*). These hubs are enriched in regulatory proteins, form chromatin modules of enhancer-promoter interactions, coordinating transcription of multiple genes for cell-specific functions (*12*, *43*).

Here, we define such regions as “genomic hubs,” representing spans of spatially clustered genes along chromosomes (**fig. S1B**). We use a previously established method, positional gene enrichments (PGEs) (*44*), to identify genomic hubs from gene sets of interest, for example, genes with common functional or transcriptional dependencies. This is followed by computational evaluation of multimodal relationships across the identified genomic regions (**fig. S1B**), and by selecting transcriptional compartment hubs (genomic hubs sharing functional relationships) for experimental validation through targeted and multimodal phenotyping methods (**fig. S1B**).

Building on this, we applied this approach to investigate regions with human-specific mutations (*8*, *9*) (**Fig. 1C, fig. S1B**). This consisted of identifying HAR-associated genes that share functional, transcriptional, and cell-type enrichments, followed by determining whether these genes positionally cluster into specific genomic hubs (**fig. S1C**). We further investigate these regions by assessing their functional annotation, genome structural features, genetic variants and phenotypes, along with murine conservation. Based on this, we categorize and select candidate regions for in-depth experimental validation in model organisms and cell systems (**fig. S1D-H**). Our goal was to understand how distinct transcriptional compartment hubs linked to evolutionary adaptations respond to nutrient stress.

#### 2.1. Metabolic HAR gene networks are linked to PGC1A and its regulatory co-factors

We began by identifying enriched molecular signatures, particularly in organs relevant to the metabolic traits we observed. Given our comparative trait analyses highlighting fat mass, we evaluated previously reported human-primate high-throughput data (*45*, *46*) as well as HAR-associated genes (*8*, *9*) (**fig. S2A**). We performed functional annotation of enriched signatures in tissue-specific networks as they capture relevant biological information from a range of experimental data (*47*). Transcriptomic and epigenomic data from brain and adipose tissue revealed that molecular signatures enriched in *Homo sapiens* are associated with transcriptional, mitochondrial, and lipid-related modules (**fig. S2B,** Table S**1E-H**). HAR-associated genes (within 1Mb of HARs) (*8*, *9*) showed similar biological enrichments (**fig. S2C,** Table **S1I-M**), suggesting that lineage-specific signatures share functional relationships, both from high-throughput data and from genes linked to HAR elements.

To refine our analysis and identify functional gene programs, we investigated whether a subset of HAR genes shares transcriptional regulation. To identify cooperative transcriptional regulators, we evaluated binding enrichment using data repositories (ChIP-seq and transcription factor binding sites (TFBS) enrichment), followed by filtering regulators with protein-protein interactions (PPIs), similar biological function, and HAR association (**fig. S2D,** Table **S2A-J**). This revealed that HAR genes are bound by transcriptional regulators with similar biological signatures and physical interactors (**fig. S2D, E,** Table **S2-S3A,B**). Network analysis of all possible interactors using PPIs showed co-activators and nuclear receptors with adipocyte function and lipid metabolism (**Fig. 1D, fig. S2E-G,** Table **S3C-G**).

Interestingly, from these interacting regulators, only CREB1 and PGC1A have genes associated with HAR elements (*8*, *9*) (**Fig. 1E**), highlighting the role of PGC1A as a hub regulator via interactions with metabolic transcription factors (*28*) and its association with rapidly evolving genomic elements. To pinpoint gene sets with higher likelihood of transcriptional co-dependencies, we filtered HAR genes with cooperative binding enrichments by these interacting regulators. This approach revealed a subset of around 40% of HAR-genes co-bound by at least 2 PGC1A co-regulators, associated with metabolic function, and enriched in adipocytes and adipose tissue (termed metabolic-HAR or mHAR genes; **Fig. 1F. fig. S2H, I,** Table S**4A-E**). These findings identify subsets of lineage-specific signatures that share metabolic and transcriptional enrichments, highlighting the need to further understand how distinctive genomic adaptations impact metabolic regulation.

#### 2.2. Metabolic HAR genes cluster in specific genomic hubs

Having identified a functionally enriched set of mHAR genes, we next asked whether chromosomal organization reveals key structural principles. We examined whether metabolic HAR genes are located in specific genomic regions, display specific genome structural features, and exhibit conservation in organismal models (**fig. S1D-G**). Using positional gene enrichment analysis (*44*) (PGEs), we first determined mHAR genomic hubs (**Fig. 1G, fig. S3A-B**). We found that mHAR genes are positionally clustered in distinct genomic hubs (1-15mb), more so than in random gene sets permutations (**fig. S3A-B**). Annotation of the positionally clustered genes for each identified genomic hub revealed specific functional and phenotypic enrichments (fPGEs, or functional mHAR genomic hubs, **Fig. 1G**, Table **S4**). For example, genomic hubs with robust positional clustering exhibit functions and phenotype enrichments related to lipid, glucose, and mitochondrial as well as disease enrichments, including lipid disease and diabetes (**Fig. 1G**, Table **S4C-E**). Overall, identifying metabolic HAR genomic hubs with specific functional enrichments suggest that these regions might share higher-order genome structural features that facilitate coordinated gene regulation.

#### 2.3 Metabolic HAR genomic hubs share function and nuclear proximity

To investigate genome-wide structural relationships for mHAR genomic hubs, we implemented a computational graph-based pipeline that evaluates their relationships using a compendium of chromatin contact maps from previous studies (*48*, *49*) (**fig. S3C**). We used studies that identified conserved frequently interacting regions (FIREs) in HiC maps from 21 primary human tissues and cell types (*48*), and another study that identified interacting genomic regions enriched in active epigenetic marks from human embryonic stem cells (ESCs) (*49*). Compared to genomic hubs from random gene set permutations, metabolic HAR genomic hubs are often located in interacting regions (**Fig. 1H**, Table **S5A-C**), suggesting that these hubs share genome structural features that might impact gene coregulation.

Given that chromatin contacts influence gene co-expression (*12*, *49*), we evaluated the spatial relationships between mHAR genomic hubs. We used an interaction network from hESCs HiC maps (*49*) (nodes are regions and edges are contacts), followed by graph-based modeling and analysis of regions overlapping mHAR genomic hubs (**fig. S3D,** Table **S5**). This analysis revealed that mHAR genomic hubs are positioned in highly interconnected chromatin regions (**fig. S3E,** Table **S5D, E**). We also focused on bridging regions as they are critical links between subnetworks. For example, hubs with high connectivity harbor mHAR-genes associated with lipid and glucose function, such as the top bridging region around Chr1:47-52mb (**fig. S3F, G**). Further verification and in-depth examination across various HiC datasets, including 3D reconstruction, confirmed the interactions and close spatial positioning of chromosomes containing these connected mHAR hubs (**fig. S4A-D, fig. S5A-F**).

This approach revealed subnetworks of interconnected mHAR hubs linked to the chr1:48-63mb region (and other hubs in chr2, chr4, chr8, and chr11), sharing lipid and glucose functional enrichments (**Fig. 1I, fig. S5G**). Using the vista repository to identify active HAR elements, we found that nearly 84% of mHAR genomic hubs contain experimentally validated non-coding elements in human embryos (**fig. S5H**). Moreover, each mHAR hub within the interacting nuclear compartment exhibits active elements, suggesting that active human-specific mutations are clustered in interacting regions linked to metabolic function (**fig. S5I**, **fig. S6A, B**).

Next, we evaluated enhancer-promoter associations in these mHAR genomic hubs using data from the GeneHancer repository (*50*) (coordinates of enhancer-promoter links; **fig. S6C**). Graph-based modeling and analysis revealed modules of mHAR-associated regulatory regions (**fig. S6D, E,** Table S**6**). By focusing on functional mHAR hubs with structural dependencies (**fig. S6F**), we determined local transcriptional regulators by binding enrichment analysis of promoters and enhancers within modules, followed by identifying those with PPIs and similar biological function (**fig. S6G-I,** Table S**6, S7**). This approach revealed that enhancers and promoters within mHAR-associated modules exhibit transcriptional and biological enrichments related to adipocyte function and glucose metabolism (**fig. S6I,** Table **S7**). Overall, these results suggest that certain genomic hubs with human-specific mutations exhibit functional associations with energy metabolism, with distinctive higher order and local genome structural features.

#### 2.4 Metabolic HAR genomic hubs with nuclear proximity show metabolic trait association

The structural features that were observed suggested that these regions might harbor shared generic variants or variants linked to metabolic traits. We assessed genetic variant enrichments using the common metabolic disease portal, which estimates variant effects for genomic regions through meta-analysis of associations (*51*). We determined the most significant associations by phenotype group and top traits within each mHAR genomic hub (**Fig. 1J**, Table S**8**). For example, the genomic hub in chr1:63mb displayed enrichments for lipid and glucose traits such as triglycerides and type 2 diabetes (**Fig. 1J**, Table S**8**), consistent with the annotation of mHAR genes in this hub. Other mHAR genomic hubs with functional and spatial relationships (within chr1, 2, 4, 8, and 11) revealed consistent phenotypic groups and top-trait associations for each and combined genomic hub, including lipid, hematological, and glycemic groups, and the top-trait waist-hip ratio (**Fig. 1K, fig. S6J,** Table S**8**).

These results suggest that mHAR genomic hubs, sharing functional, transcriptional, structural, and genetic features might have played a role in fine-tuning the activity of these regions via genetic substitutions during evolution (*52–54*). This also indicates that these regions might be syntenic with similar trait associations in other mammals, including organismal models such as mice (*52–54*).

#### 2.5 Metabolic HAR genomic hubs exhibit murine synteny and phenotype conservation

Before testing the activity of these regions experimentally, we aimed at establishing whether they are conserved in model organisms. To evaluate the degree of murine conservation of mHAR hubs, we examine their chromosome synteny and phenotype association in mouse models. About 80% of these hubs maintained their clustering and synteny (**Fig. 1L**, **fig. S7A,** Table S**9**). Interestingly, some genomic hubs, including those with similar function and nuclear spatial proximity, were split in murine chromosomes (e.g. hChr 2, 4, 7, and 18; **Fig. 1L**, Table S**9**), suggesting specific rearrangements in humans.

To evaluate phenotype associations in these hubs, we built a network of relationships between mHAR genes and their knockout traits from mouse repository data. Graph-based analyses revealed key modules such as metabolic and immune, and ranked phenotypes and genes by connectivity (global or modular network connectivity; **Fig. 1M, fig. S7B, C**). Metabolic phenotypes, for example, showed the highest connectivity scores across the network (**fig. S7C**), including significant shared enrichments (**fig. S7D**). Genes within mHAR genomic hubs that share structural and functional associations were mostly located in metabolic, immune, and CNS modules, and exhibited a high degree of connectivity such as Pgc1a in the metabolic module (**Fig. 1M, fig. S7E**). This analysis further revealed that these modules are highly associated with each other (**fig. S7F**), and that their genes are positionally colocalized in specific genomic hubs (**Fig, 1M, fig. S7G, H**), supporting conserved function and phenotypic associations in evolutionary tuned genomic regions.

In summary, our computational integration and analysis suggest links between energy metabolism, use, and allocation across mammals and particularly in hominids. We identified genomic regions enriched in human-specific mutations, associated with metabolic function and traits in humans, and conserved in structure and phenotype in mice. mHAR hub genes are enriched in adipose tissue, are associated with lipid and glucose metabolism, and are bound by metabolic factors such as PGC1A and coregulators. These conserved features in mice allowed us to test mechanistic principles experimentally, while the human-specific mutations in mHAR elements suggested additional evolutionary tuning that we investigate in cellular models. In the following sections, we evaluate the activity of these transcriptional hubs in murine models to understand their conservation and their impact on metabolic adaptations and disease (**fig. S7I**).

### 3. mHAR hubs and coregulators show cell-specific activity during nutrient stress in murine models

Having identified candidate genomic hubs computationally, we next tested whether these regions are transcriptionally active during nutrient stress. Given our adipose tissue metabolic trait association with the fasting endurance score in mammals, we next explored mHAR genomic hub activity during nutritional stress by focusing on syntenic murine regions for the human Chr1:48-63mb genomic span, as it contains highly connected hubs linked to lipid-glucose metabolism and to similar phenotypes in mice (**Fig. 2A**). Our goal was to first evaluate mHAR gene activity and transcriptional compartment hub formation after nutrient limitation.

**Fig. 2.**
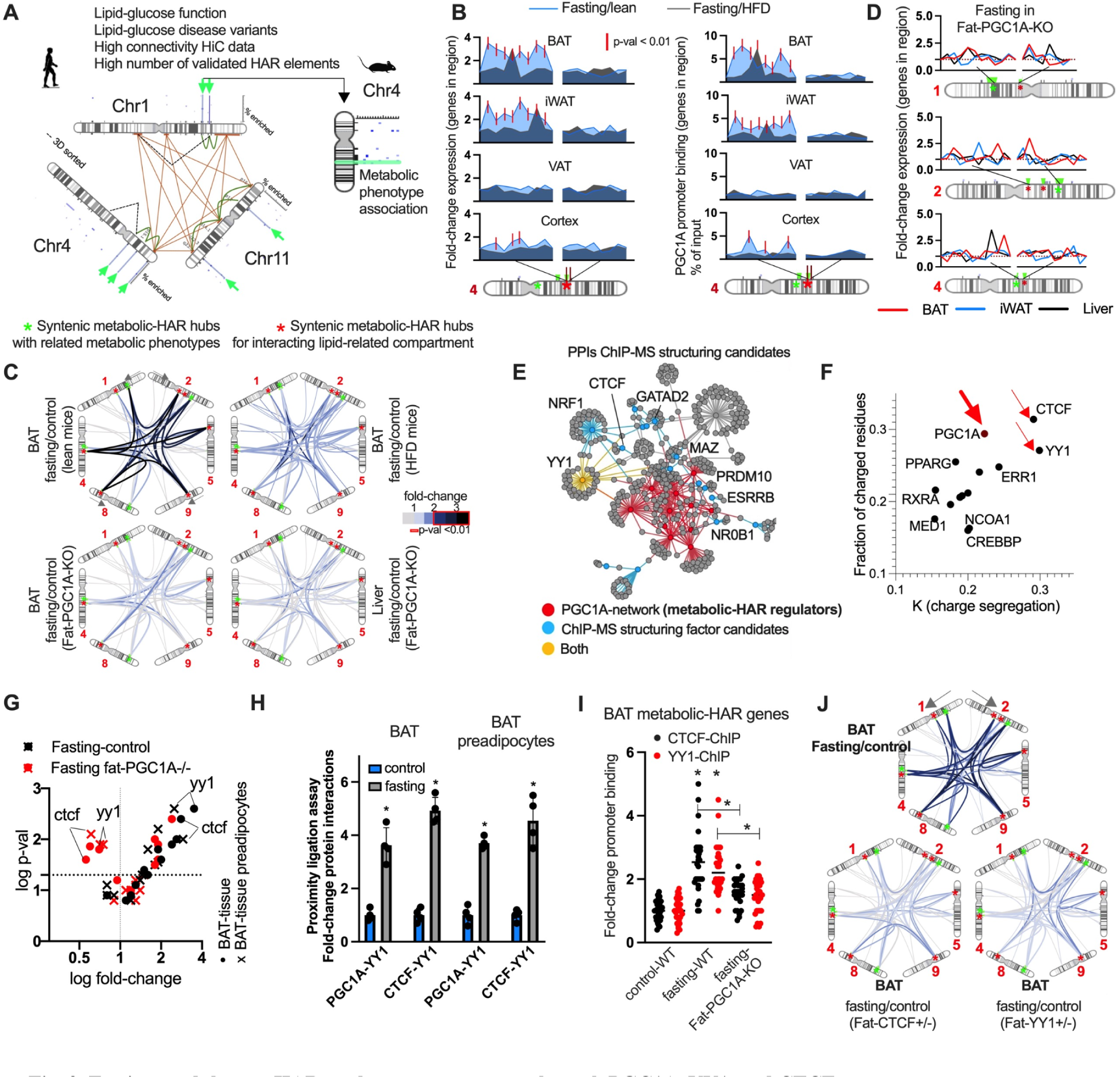
Fasting modulates mHAR nuclear compartments through PGC1A, YY1, and CTCF. **(A)** Illustration showing mHAR associated nuclear compartment and functional genomic hubs (in green). It displays features for mHAR genomic hub in chromosome 1, as well as syntenic hub with phenotype conservation. **(B)** left panel, gene expression of hub-forming genes in mHAR region (chr4:97-100mb), and neighboring region. Each peak represents gene expression (arranged by genomic positions), after fasting (24h) in tissues from lean and HFD (1 week) mice. Red line shows significant changes between groups p-value<0.01. Right panel, ChIP-qPCR fold-change binding by PGC1A to gene promoters within the hub. **(C)** Targeted *in situ* chromosome conformation assays (3C) for mHAR associated nuclear compartment in tissues from lean and HFD (1 week) wildtype mice, and fat-Pgc1a-/- mice after fasting (24h). Targeted contact variations are represented as fold-change over control groups (not fasted littermates). Contacts with FC > 2 have p-value <0.01. **(D)** Fold change gene expression of hub-forming genes in mHAR associated nuclear compartment hubs as described in **B**. **(E)** Protein-protein interactions for PGC1A cooperative transcriptional regulators (in red) and structuring transcriptional candidates (in blue) selected by previously reported ChIP-MS data (*55*). In yellow shared regulators. **(F)** CIDER analysis of conformational protein features (from amino acid sequences). Graph shows fraction of charge residues and charge segregation across regulators. **(G)** Gene expression for PGC1A-cooperative transcriptional regulators in tissues and sorted cells from control and fat-Pgc1a-/- mice after fasting (24h). **(H)** Proximity ligation assays for transcriptional interactions from tissue extracts in mice described in **G**. **(I)** ChIP-qPCR binding of YY1 and CTCF to mHAR genes within the nuclear compartment. Each dot represents the fold-change variation of their binding to different promoter regions. **(J)** Targeted *in situ* chromosome conformation assays (3C) for mHAR associated nuclear compartment compartment in brown fat from control and fat-specific Yy1+/− and Ctcf/- heterozygous mice after fasting (24h). Contact variations shown as described in C. Animal experiments were done with n=5-8 mice per group, both female and male mice were included. Data show mean values and SEM. Cell experiments were done with 3 independent replicates. Chi-square test was used to determine the statistical significance between expected and observed frequencies. Unpaired, two-tailed student’s t-test was used when two groups were compared, and ANOVA followed by fisher’s least significant difference (LSD) test for post hoc comparisons for multiple groups. * p-value <0.05.

In lean but not in HFD mice (1 week), acute fasting induced the expression of mHAR genomic hub genes and PGC1A binding to their promoters in brown fat (BAT), inguinal fat (iWAT), liver, sorted pre-adipocytes, and hepatocytes, with no significant changes in brain cortex, visceral fat, and sorted monocytes (**Fig. 2B, fig. S8A-C**). In lean but not in HFD mice, fasting increased relative chromatin contacts between mHAR genomic hubs (interrogation of mHAR gene promoter regions interactions using a targeted 3C approach), as well as gene expression within these hubs (**Fig. 2C, fig. S8D-G**). This response was absent in adipose-specific (AT) Pgc1a-/- mice (**Fig. 2C, D, fig. S8H-K**), suggesting that, during fasting, functional mHAR genomic hubs are transcriptionally active, form chromatin contacts indicative of co-transcriptional hubs, and are under PGC1A regulation, as adipocyte perturbation blunts their regulation locally and systemically (**fig. S8L-M**).

These results established PGC1A as central to mHAR hub activation, but PGC1A typically works with other cofactors. To select co-regulators of mHAR hubs, we examined different features in PGC1A interactors. These include chromatin structuring, protein interaction and domain conformations, as well as PGC1A regulation. We used previous studies that identified mediators of enhancer-promoter contacts via ChIP-mass spectrometry (ChIP-MS) (*55*, *56*), followed by assessing their link to PGC1A using PPIs. This revealed known structuring factors such as CTCF, factors with metabolic function such as NRF1, and factors like YY1 shown to interact with PGC1A during metabolic adaptations (*57*) (**Fig. 2E**, Table S**10**).

To complement this, we evaluated protein conformational features based on amino acid sequences using the classification of intrinsically disordered ensemble regions tool (CIDER) (*58*), which revealed that PGC1A and the structuring factors YY1 and CTCF display protein regions with similar features such as charged residues and charge segregation (**Fig. 2F, fig. S9A**). These features are linked to protein droplet properties in context-dependent environments such as solvation or protonation, which might impact transcriptional plasticity (*58*).

We next evaluated coregulators gene activity, revealing that Yy1 and Ctcf show increased gene expression in BAT, iWAT, liver, and sorted cells from fasted control mice, but absent in fasted AT-Pgc1a-/- mice (**Fig. 2G, fig. S9B**). Proximity ligation assays showed increased relative protein interactions in fasted control mice, but reduced in AT-Pgc1a-/- mice (**Fig. 2H, fig. S9C**). In control mice, fasting increased CTCF and YY1 binding to the same mHAR genomic hub promoters, which was absent in AT-Pgc1a -/- mice (**Fig. 2I, fig. S9D**). In addition, mHAR hubs chromatin contacts seen in fasted control mice were absent in both liver and fat from AT-Yy1+/− and AT-Ctcf+/− heterozygous mice (**Fig. 2J, fig. S9E-G**).

In summary, mHAR genomic hub activity is modulated by cooperative regulators, such as PGC1A, YY1, and CTCF. These regulators share protein conformational features, interact with genome structuring candidates, and are under PGC1A regulation during fasting in adipose tissue. These results suggest that their cooperative activity might contribute to genome structural plasticity during nutrient stress.

### 4. mHAR hub genes regulate acute fasting endurance through lipid and lysosome homeostasis

Having established that mHAR hubs are active and coregulated during fasting, we next sought to identify the specific genes within these hubs that mediate functional adaptations. Before assessing their role in genome plasticity regulation, we first aimed at identifying critical genes within the mHAR genomic hubs that are under control by these coregulators, potentially mediating functional adaptations to nutrient stress. In mice with fat-specific perturbation of mHAR coregulators, 24 hour fasting reduced the expression of mHAR hub genes associated with sterol, oxysterol, and peroxisome pathways in brown and white adipose tissue, including Dgat1 and Atg4c genes reduced across all mutants (**Fig. 3A, fig. S10A**).

**Fig. 3.**
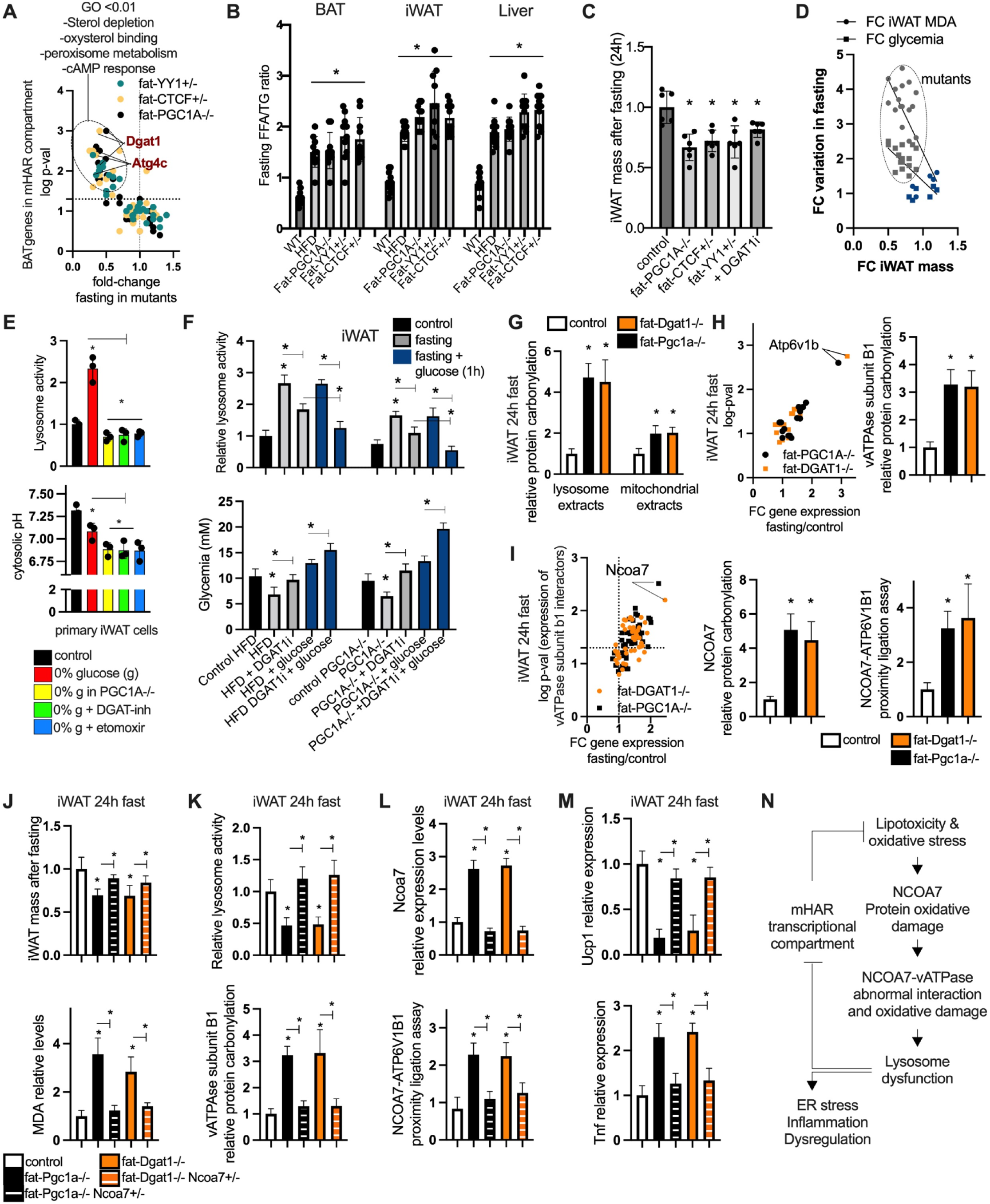
mHAR genomic hubs regulate acute fasting endurance through lipid and lysosome homeostasis. **(A)** Expression of mHAR genes within associated nuclear compartment in inguinal fat from fat-specific mutant mice after fasting followed by functional enrichment of significant genes. **(B)** Relative free fatty acids/triglycerides tissue ratio after fasting (24h). **(C)** Relative inguinal fat mass variation in fasted (24h) control, fat-mutant mice, and mice treated with DGAT1-inhibition (2 mg/kg, 24 h i.p.). **(D)** Fold-to-fold change plot showing fat mass, glycemia, MDA lipid peroxidation variation in fasted control and mutant mice. **(E)** Primary cell cultures (brown and white adipocytes) from wildtype and PGC1A-/- mice treated with 0% glucose (12 hours), and, as indicated, glucose deprivation in combination with DGAT1-inhibitor (1μM) and the Cpt1a channel inhibitor etomoxir (50μM). Cells assessed for relative lysosome activity (top panel) and cytosolic pH levels (lower panel). **(F)** Inguinal fat relative lysosome activity (top panel) after 24 hours fasting and 24-hour fast with glucose injections (1hour, by i.p prior evaluation) in HFD (1 week), HFD + DGAT1-inhibition mice (2 mg/kg, 24h i.p), fat-mutant mice, and fat-mutant mice treated with DGAT1-inhibition. Blood glucose levels (lower panel) in the same conditions and quantified 150 min after glucose injections. **(G)** Protein carbonylation assays in inguinal fat protein extracts (lysosomal and mitochondrial) from fasted control and mutant mice. **(H)** Left panel, proton pumps gene expression screen in inguinal fat tissue from fasted control and mutant mice. Right panel, inguinal fat relative protein carbonylation in vATPase subunit B1 from fasted control and mutant mice. **(I)** Left panel, inguinal fat gene expression screen for vATPase subunit B1 protein interactors (from biogrid) as in H. Middle panel, inguinal fat relative protein carbonylation in NCOA7 and, in right panel, protein relative interactions by proximity ligation assays in mice as in H. In fasted (24h) fat-specific mice with single or double genetic perturbations, inguinal fat mass and MDA lipid peroxidation levels **(J)**, inguinal fat lysosome activity and vATPase-protein carbonylation **(K)**, inguinal fat Ncoa7 expression and NCOA7-vATPase interaction by PLA assays **(L)**, and inguinal fat mass Ucp1 and Tnf expression **(M)**. **(N)** Summary illustration for NCOA7-mediated perturbation of fasting adaptations and its relationship with mHAR-associated transcriptional compartments. Animal experiments were done with n=5-6 mice per group, both female and male mice were included. Data show mean values and SEM. Cell experiments were done with 3 independent replicates. Unpaired, two-tailed student’s t-test was used when two groups were compared, and ANOVA followed by fisher’s least significant difference (LSD) test for post hoc comparisons for multiple groups. * p-value <0.05.

ATG4C, linked to autophagy, participates in generating fatty acids through lipid droplet autophagy (*59*), while DGAT1 esterifies fatty acids into triglycerides (TGs). During fasting, increased DGAT1 activity has been associated with reductions in ER stress, mitochondrial dysfunction, and lipotoxicity (*37*, *38*). In humans, the Atg4c gene is located in the Chr1:63mb region, within a mHAR genomic hub associated with high chromatin connectivity (**fig. S10B**). The Dgat1 locus is in a highly interactive Chr8:144mb region, which is structurally associated with the functional hubs sharing nuclear proximity (**fig. S10B**).

Given its different location in mice, we evaluated chromatin contacts between mHAR Atg4c and Dgat1 hubs in our murine models. This showed that fasting increased chromatin contacts between these hubs in BAT, iWAT, and liver from control mice, but not in mice with deletion of coregulators (**fig. S10C**). This suggests that context-dependent activity and interactions are conserved despite Dgat1 region arrangement. Furthermore, the cooperative transcriptional regulation of these hubs highlights the importance of Dgat1 and Atg4c for lipid mobilization and fasting adaptations.

Next, we evaluated the free fatty acid (FFA)-to-TGs ratio in tissues with mHAR genomic hub activity. After a 24-hour fast, the FFA-to-TGs ratio increased in BAT, iWAT, and liver from both HFD (1 week) and mutant mice compared to lean controls (**Fig. 3B**), supporting the link between FFA and mHAR genomic hub activity. We next assessed the impact of 6-hour fasting on TG accumulation after intraperitoneal (i.p.) injections of DGAT1- and autophagy inhibitors (bafilomycin) in BAT of control and fat-specific mutant mice. TG formation was observed in both groups after a 6-hour fast but was hindered by inhibiting either DGAT1 or autophagy (**fig. S10D**), suggesting a time-dependent saturation of fatty acid esterification.

Subsequently, we examined the relationship between FFA accumulation and lipid peroxidation and its deregulation in our models. Increased tissue levels of malondialdehyde (MDA), a lipid peroxide, were found in BAT, iWAT and liver from mutant mice and mice with inhibited DGAT1 or autophagy (**fig. S10E**). We found a reduction in iWAT mass in our models, which inversely correlated with MDA levels and glycemia (**Fig. 3C, D, fig. S10F**).

These results suggest that inhibiting mHAR genomic hub activity through dietary factors, fat-specific deletion of coregulators, or pharmacological inhibition, reduces FFA esterification and increases MDA levels after fasting, influencing lipid mobilization endurance, genome adaptations, and fat mass (**Fig. 3C, fig. S10G**). Given these findings, we focus on understanding the molecular mechanisms within adipose tissue.

The accumulation of lipid peroxides suggested potential damage to cellular organelles, particularly those with lipid-rich membranes. Oxidative damage to membrane lipids in mitochondria and lysosomes can disrupt metabolic homeostasis and pH balance, especially during nutrient deprivation (*60–62*). In this condition, mitochondrial ATP generation and lysosomal proton transport regulate cellular pH, necessary for transcriptional adaptations (*63–66*). In line with this, we next investigated how variations in nutrients such as glucose impact lipid-pH homeostasis and mHAR genome plasticity.

In primary brown and white adipocytes, we found that glucose deprivation alone increased mitochondrial and lysosomal activity while reducing cytosolic pH (**Fig. 3E, fig. S10H-J**). The same response was impaired in cells lacking Pgc1a or treated with DGAT1-inhibitor or etomoxir (a mitochondrial CPT1 inhibitor that reduces fatty acid import and oxidation), which showed reduced respiration and lysosomal activity, while increasing cytosolic acidity and lipid peroxidation (**fig. S10H-K**). We next tested in our *in vivo* models if fat tissue pH is modified during fasting and is linked to glucose levels. Fasted mutant mice exhibited lower tissue pH in BAT and iWAT compared to control fasted mice (**fig. S11A**). This was associated with the effect of glucose on fasted adipose tissue, as a single dose of glucose injections (i.p. 1h) after 24 hour fasting normalized tissue pH in lean control and HFD mice, but not in HFD mice treated with DGAT1 inhibitors or fasted mutant mice (**fig. S11B**).

We then evaluated if glucose was being metabolized differently in both models given our observations on mitochondrial and lysosome function. DGAT1 inhibition in fasted HFD and AT-Pgc1a-/- mice reduced mitochondrial and lysosome activity, increased lactate levels after glucose injections (1h i.p.) in fat depots, which might contribute to glucose intolerance (**Fig. 3F, fig. S11C, D**). We tested if these metabolic disruptions influence transcriptional plasticity. In both fat depots, DGAT1-inhibition during fasting reduced mHAR genomic hubs chromatin contacts (**fig. S11E, F**). This suggests a link between mHAR hub transcriptional activity and adaptation to nutrient deprivation, as they regulate genes influencing lipid, lysosome, and pH homeostasis (**fig. S11G**).

### 5. NCOA7 modulation influences acute fasting endurance

Oxidative damage caused by lipid peroxidation can impair protein function and interactions (*67*, *68*), suggesting a potential molecular mediator of the phenotypes we observed. Given the iWAT mass reduction, we focused on this tissue to identify mediators of oxidative damage. First, we observed increased relative carbonylation (oxidative damage) in protein extracts from lysosomes in fasted AT-Pgc1a-/- and AT-Dgat1-/- mice, compared to fasted control mice (**Fig. 3G**). Since lysosomes contribute to pH homeostasis, we evaluated expression levels of proton pumps. In fat depots from fasted mutant mice, we found a significant upregulation of vATPase subunits, particularly the Atp6v1b1, which also exhibited increased oxidative damage in targeted protein extracts (**Fig. 3H**). This suggests that lipid-mediated oxidative damage of vATPase might contribute to fasting endurance through disrupting lysosomal and pH homeostasis.

vATPase complexes interact with proteins that modulate their activity and assembly (*69*). By using the biogrid database we identified vATPase protein interactors and measured their expression in our models. This revealed a substantial upregulation in Ncoa7 expression in iWAT and BAT (**Fig. 3I, fig. S12A**). NCOA7 is a transcriptional coactivator associated (through direct interaction) with oxidative stress response proteins such as OXR1 (*70*, *71*). We next investigated if oxidative damage in NCOA7 could subsequently impair lysosome function via vATPase interaction. NCOA7 protein carbonylation increased substantially in iWAT from fasted mutant mice compared to fasted lean control (**Fig. 3I**). Proximity ligation assays revealed increased relative interactions between NCOA7 and the Atp6v1b1 subunit in the same models (**Fig. 3I**), suggesting a potential link between oxidative stress, NCOA7, and lysosomal function during fasting.

To evaluate the effect of NCOA7 perturbation, we generated mice carrying double perturbations including adipocyte-specific Pgc1a-/- and whole body heterozygous Ncoa7+/−, as well as fat-Dgat1-/- and Ncoa7+/−. Previous studies have shown that, in unchallenged conditions, AT-Dgat1-/- mice appeared normal with moderately reduced body fat, but upon fasting (16h), display lipotoxicity, ER stress, and inflammation (*38*, *72*). Ncoa7+/− mice showed normal lysosome function, no protein carbonylation excess in lysosome extracts, maintaining low Ncoa7 transcript levels after fasting (**fig. S12B-D**). In contrast, fat Dgat1-/- or Pgc1a-/- mice reduced iWAT mass and increased MDA levels in both fat depots, which was restored when Ncoa7 was also perturbed (**Fig. 3J, fig. S12E**).

In both fat depots, reducing Ncoa7 expression improved lysosome function, reduced vATPase oxidation, and reduced Ncoa7-vATPase interaction (**Fig. 3K, L, fig. S12F, G**), highlighting Ncoa7 as a mediator of oxidative stress impairing lysosome function. To understand Ncoa7 regulation, we evaluated functional enrichments in co-expressed genes (archived at ARCHS4 RNAseq repository). This revealed links to proinflammatory pathways including IFNG, NFKB, and STAT1 signaling (**fig. S12H**). Accordingly, mice with impaired fasting adaptations increased STAT1 binding to Ncoa7 promoter, while mice with double perturbation reduced this response (**fig. S12I**). In both fat depots, this led to increased thermogenic Ucp1 gene expression, reduced expression of Tnf and ER-stress marker gene Xbp1s (**Fig. 3M, fig. S12J, K**). This was associated with increased mHAR genomic hub chromatin contacts and gene expression in mice with double perturbation (**fig. S12L-N**). These findings suggest that Ncoa7 impairs acute fasting adaptations in adipose tissue, which impact mHAR activity and genome plasticity (**Fig. 3N**).

### 6. NCOA7 influences genome plasticity and metabolic homeostasis in intermittent fasting

The results on acute fasting suggested that NCOA7 is a key mediator of maladaptation yet how this is translated to chronic interventions remains unknown. In addition, IF is known to improve metabolism and overall health, although the underlying molecular mechanisms remain unclear. In previous studies, AT-Dgat1-/- deletion mice on a HFD showed decreased body weight and fat mass, glucose intolerance, and ER-stress (*72*). In this study, AT-Dgat1-/- mice and mice with decreased Dgat1 expression (mHAR coregulator knockouts) have a compromised adaptation to acute fasting, with for example reduced fat mass, glucose intolerance, lipotoxicity, impaired lysosome function, and reduced genome plasticity. To investigate how these mice adapt over time we examined intermittent fasting (IF) post-HFD, in which mice are fasted for 24 hours twice a week for six weeks.

In normal HFD mice IF reduced body weight gain, fat mass (inguinal), and enhanced glucose tolerance (**Fig. 4A-C**). In the AT-Dgat1-/- HFD mice, IF resulted in a greater reduction of body weight and fat mass, and glucose intolerance (**Fig. 4A-C**). In HFD mice with double deletion of AT-Dgat1-/- and whole body Ncoa7+/− deletion, IF showed less pronounced body weight and fat mass reduction, improved glucose tolerance, despite not changing food intake (**Fig. 4A-C, fig. S13A**).

**Fig. 4.**
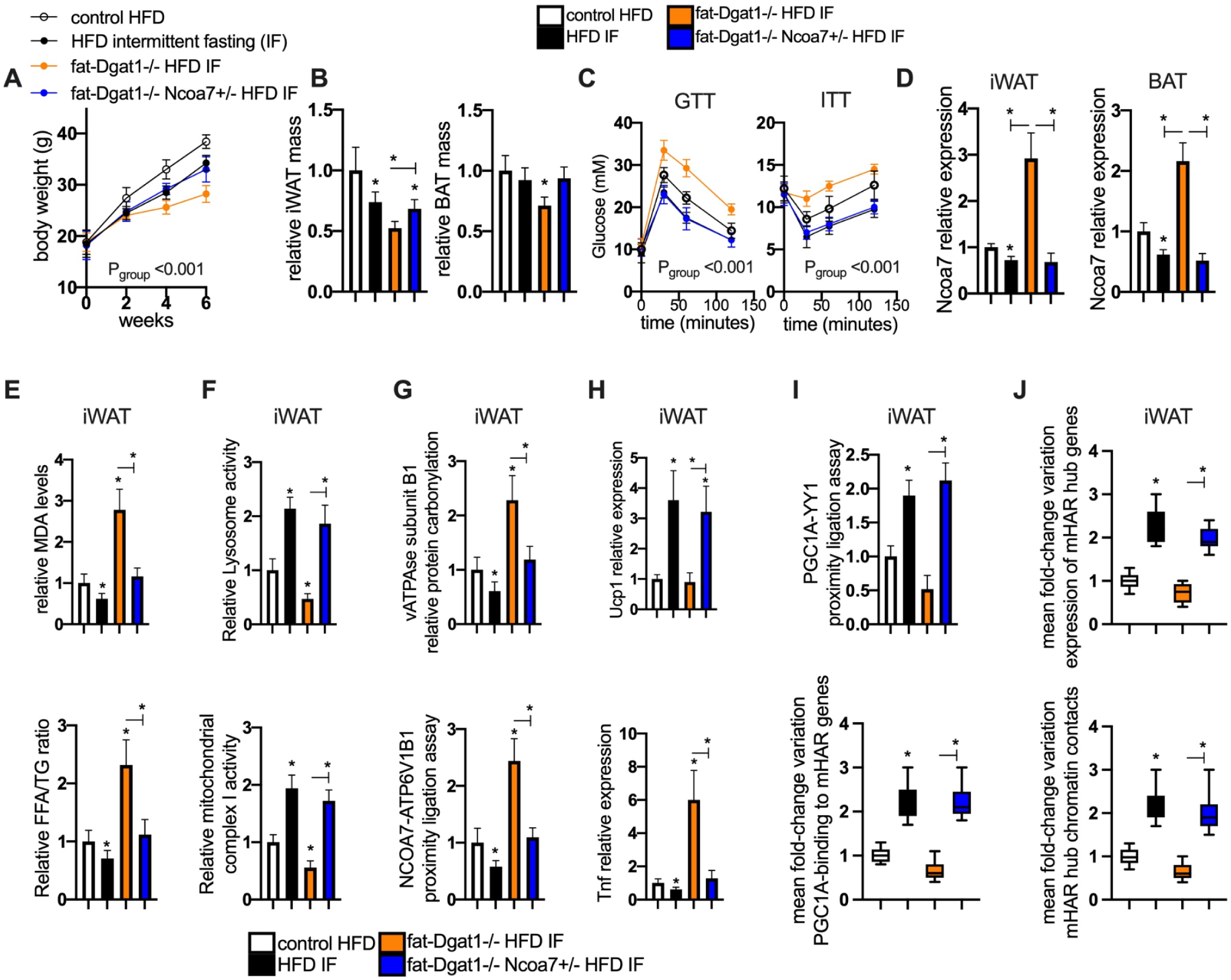
Ncoa7 modulation influences metabolic homeostasis in intermittent fasting. **(A)** Body weight evolution in control HFD mice and mice (control and mutant mice) on a HFD and treated with 2-day bouts of intermittent fasting (IF) every week. **(B)** In mice treated as in A, relative inguinal (left panel) and brown (right panel) fat mass variation. **(C)** Glucose (left panel) and insulin (right panel) tolerance test in mice treated as in A. **(D)** Ncoa7 gene expression in inguinal (left panel) and brown (right panel) fat from mice treated as in A. **(E)** Inguinal fat MDA lipid peroxidation levels (top panel) and free-fatty-acids/TG ratio (lower panel) from mice as in A. **(F)** Inguinal fat lysosome (top panel) and mitochondrial activity levels (lower panel) from mice as in A. **(G)** Inguinal fat vATPase subunit protein carbonylation (top panel) and NCOA7-vATPase subunit relative interaction levels (lower panel) from mice as in A. **(H)** Ucp1 (top) and Tnf (down) gene expression. **(I)** Proximity ligation assays for protein interactions (top panel) and PGC1A binding by ChIP-qPCR to mHAR genomic hub genes within the associated nuclear compartment (lower panel). Data in the lower panel is shown as mean fold-change variations in binding to at least 10 mHAR genomic hub promoters. **(J)** Top panel, mean fold-change variation in expression of mHAR genomic hub genes (within associated compartment, n=12-16 genes) in inguinal fat from mice as in A. Lower panel shows chromatin contacts (targeted chromosome conformation assays) between mHAR genomic hubs shown as mean fold change variation in at least 10 contacts per group. Animal experiments were done with n=5-6 mice per group, both female and male mice were included. Data show mean values and SEM. Unpaired, two-tailed student’s t-test was used when two groups were compared, and ANOVA followed by fisher’s least significant difference (LSD) test for post hoc comparisons for multiple groups. Two-way ANOVA was used to estimate significance between groups constrained by time measurements, and Tukey test for multiple comparisons. * p-value <0.05.

Allelic deletion of Ncoa7, like IF, reduced MDA levels and FFA/TG ratio, restored lysosome and mitochondrial function, and decreased vATPase carbonylation and Ncoa7 interaction (**Fig. 4D-G, fig. S13B-D**). This resulted in enhanced thermogenic Ucp1 gene expression, reduced Tnf and ER-stress marker gene expression (**Fig. 4H, fig. S13E, F**). This improved the systemic inflammation seen in AT-DGAT1-/- mice, with reduced MCP1 and TNFa in serum along with reduced expression in liver and brain cortex (**fig. S13G-I**). Similar to IF models, mice with double deletion showed enhanced mHAR genome plasticity, increased mHAR genomic hub contacts, gene expression, and coregulator protein interactions (**Fig. 4I, J, fig. S13J-L**).

Overall, these results show that Dgat1 in adipocytes helps at preserving metabolic homeostasis during chronic fasting interventions and that NCOA7 is a key node that connects oxidative stress and transcriptional dysfunction. Ncoa7 reduction rescues impaired fasting adaptations by restoring lysosome function, decreasing oxidative stress, and preserving genome plasticity. These improvements contribute to phenotypes exhibiting less inflammation and enhanced glucose tolerance, supporting NCOA7 as a key mediator of impaired adaptations, influencing transcriptional plasticity and helping us understand how nutrient deprivation influences gene regulation.

### 7. PGC1A transcriptional plasticity and NCOA7 influences cell resilience to nutrient stress

Having identified NCOA7 as a key mediator in organismal models, we next sought to understand the molecular mechanisms by which lipid and lysosome homeostasis impact transcriptional plasticity in cellular systems. We focused on PGC1A droplet properties, IDRs, and gene regulation under nutrient stress. First, we evaluated *in silico* IDR amino acid sequences, identifying disordered regions in PGC1A and interacting coregulators (**fig. S14A-D**, Table S**11**). In line with previous work showing that environmental changes influence evolutionary tuning of droplet properties (*18*, *24*), we examine the level of conservation in PGC1A residues and IDRs, including features responsive to environmental variation. We used the ConSurf package, which uses neural networks to determine conservation scales of functional and structural residues across species (*73*) (**Fig. 5A, fig. S14E,** Table **S11**).

**Fig. 5.**
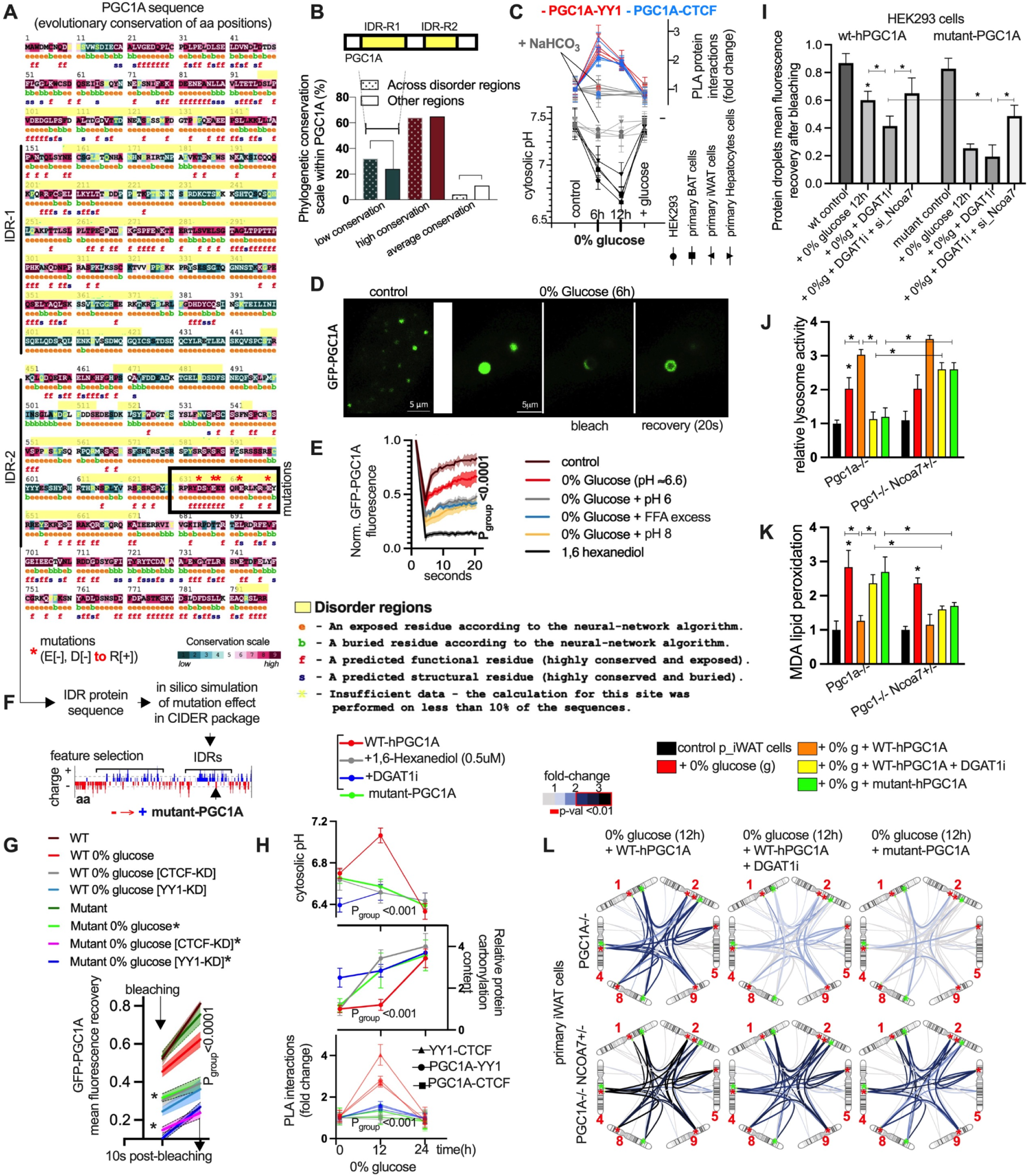
NCOA7 and PGC1A plasticity influences mHAR hub activity in nutrient stress. **(A)** Conservation scale in PGC1A amino acid (aa) residues from phylogenetic analysis using CONSURF. IDR1, 2 stretch on the side and highlighted in yellow over the aa sequence. Top down and down right shows conservation scale for each position and description of positional predictions. Square box with red asterisks shows location of site-directed mutations on highly conserved negatively charged residues (negative to positive amino acid substitution). **(B)** Proportion of conservation ranges (low, average, and high) for PGC1A aa residues within IDRs or other regions as described in A. In yellow, consensus intrinsically disordered regions. **(C)** Cytosolic pH (lower panel) and relative protein interactions by proximity ligation assays (top panel) in HEK293 cells and murine primary cell cultures in response to glucose starvation (12h) with or without NaHCO_3_ (1mM) and glucose pH correction. **(D)** Confocal microscopy images for GFP-PGC1A in HEK293 living cells in control conditions (left panel) and 0% glucose (6h; right panel) with fluorescence recovery and photobleaching assays (FRAP). **(E)** FRAP analysis, time of fluorescence recovery of PGC1A droplet in living cells in control and glucose starvation (6h), with or without pH variations (6 and >7.4), FFAs (10uM), and 1,6-hexanediol (0.5uM). **(F)** Pipeline description for in silico simulations of residue mutations affecting conformational features in CIDER. Lower panel, illustration of PGC1A-amino acid sequence, net charge variation, and IDR regions, and location of protein mutations. **(G)** Mean bleaching recovery in HEK293 cells (control or with 0% glucose conditions) transfected with wildtype or mutant PGC1A vectors (asterisk), and with siRNA for transcriptional interactors (KD=knockdown). **(H)** Glucose starved HEK293 cells (0, 12, and 24 hours) transfected with different PGC1A expression vectors (wildtype and mutant), with or without 1,6-hexanediol (0.5uM) and DGAT1-inhibitor (1μM). Top panel, cytosolic pH. Middle panel, protein carbonylation assay. Lower panel, proximity ligation assays. **(I)** Mean bleaching recovery in HEK293 cells (control or with 0% glucose conditions) transfected with wildtype or mutant PGC1A vectors, and treated together with DGAT1-inhibition (1μM) alone or with siRNA for Ncoa7. **(J)** Relative lysosome activity in inguinal primary adipocytes control (from fat-Pgc1a-/- mice or mice with double perturbations fat-Pgc1a-/- and Ncoa7+/−) and under different conditions including glucose deprivation (12h) either alone or with nucleoporation (using the nucleofector instrument and protocols for primary cells) of PGC1A vectors alone or together with DGAT1-inhibition (12h, 1μM). **(K)** Malondialdehyde (MDA) lipid peroxidation levels in cells as described in **J**. **(L)** Targeted in situ chromosome conformational assays (3C) for mHAR-associated nuclear compartment in primary cells from mutant mice in 0% glucose (12 hours), or in combination with DGAT1-inhibition (1μM). Cells transfected with either wildtype or mutant PGC1A expression vectors. Cell experiments were done with 3 independent replicates. Data show mean values and SEM. Unpaired, two-tailed student’s t-test was used when two groups were compared, and ANOVA followed by fisher’s least significant difference (LSD) test for post hoc comparisons for multiple groups. Two-way ANOVA was used to estimate significance between groups constrained by time or concentration-based measurements, and Tukey test for multiple comparisons. * p-value <0.05 when indicated and when compared to the control group.

Interestingly, PGC1A IDRs contained a higher proportion of residues with low-conservation compared to other regions (**Fig. 5B**). As our earlier CIDER analysis indicated charge-related conformational features in PGC1A (**fig. S14F**), we evaluated charged residues within these regions. Notably, positively charged residues (R, K) have a higher percentage of low conservation compared to negatively charged residues (D, E) (**fig. S14G,** Table S**11**). These observations suggest that PGC1A droplet features are influenced by charge variations in the environment, with these features ongoing evolutionary tuning.

To test whether these charge-related features influence PGC1A behavior under pH variation, we used both in vitro reconstitution and live cell imaging. To test charge variation, we first used turbidity assays, showing that recombinant PGC1A proteins increased solution turbidity upon protonation, both individually and in complexes with interactors (**fig. S14H, I**). This was affected by alkalinization, arginine, and excess acidity (**fig. S14J**). Next, we investigated how pH variation during nutrient stress affects PGC1A plasticity in living cells. In HEK293 and murine primary cells, glucose deprivation led to acidification and increased interaction with the contextual coregulators, which were reduced by correction with NaHCO_3_ and addition of glucose (**Fig. 5C**). Notably, glucose removal led to the formation of large, dynamic, and spherical PGC1A droplets, influenced by ionic strength and salt concentration (**Fig. 5D, fig. S14K**). Fluorescence recovery after photobleaching (FRAP) revealed that PGC1A droplets shift to a gel-like state with slower but still dynamic recovery during glucose shortage (**Fig. 5E**). Conversely, droplet recovery was arrested by extreme pH variation, fatty acid excess, and 1,6 hexanediol (**Fig. 5E**). These observations suggest that electrostatic regulation of PGC1A properties might influence contextual interactions during nutrient stress.

To determine whether specific residues within IDRs are responsible for these pH-dependent properties, we turned to targeted mutagenesis. Previous work has revealed that the c-terminal region in PGC1A mediates condensate formation (*31*), so we focused on its IDR (IDR2) to evaluate *in silico* (using CIDER) whether residue substitutions influence charge-related features (**Fig. 5F**). By focusing on stretches of less-conserved positively charged and highly conserved negatively charged residues, we found that mutating conserved residues (E[-], D[-] to R[+]) substantially changed charge-related features (**fig. S15A, B**). As these might potentially impact the response to proton variation, we generated a mutant PGC1A construct to assess its droplet properties, function, and plasticity (**Fig. 5F**).

In glucose-deprived HEK293 cells, PGC1A droplet recovery was reduced when YY1 and CTCF were perturbed (RNAi), with a more pronounced effect in cells transfected with mutant PGC1A (**Fig. 5G**). Arrested droplet recovery was associated with increased protein oxidation and reduced lysosome activity, particularly in DGAT1-inhibition and mutant PGC1A conditions (**fig. S15C, D**). After glucose deprivation, cells with mutant PGC1A and DGAT1-inhibition displayed increased cytosolic acidity and protein oxidative damage, as well as reduced coregulator interactions compared to cells with wildtype PGC1A (**Fig. 5H**). Using iWAT PGC1A-/- cells in unchallenged conditions revealed that mutant PGC1A (via nucleofection) was still able to regulate OXPHOS gene expression similarly to wildtype constructs (**fig. S15E, F**). This response was disrupted in glucose-deprived conditions and worsened in DGAT1 inhibition (**fig. S15E, F**), supporting a context-dependent effect on PGC1A plasticity and gene regulation.

Finally, we tested whether NCOA7 (our identified mediator of lysosomal dysfunction) also controls PGC1A droplet dynamics. In glucose-starved HEK293 cells, Ncoa7 deletion rescued the arrested droplet recovery induced by DGAT1-inhibition and mutant-PGC1A (**Fig. 5I**). In primary adipocytes, glucose-starved PGC1A-/- cells transfected with wildtype PGC1A restored pH balance and lysosomal activity, reducing protein oxidation, lipid peroxidation, ER stress and inflammatory gene expression, compared to cells with mutant-PGC1A or in DGAT1-inhibited conditions (**Fig. 5J, K, fig. S15G-J**). Glucose deprivation increased mHAR hub chromatin contacts and transcriptional interaction only in cells with wildtype PGC1A (**Fig. 5L, fig. S15K**). Notably, adaptations that were impaired in mutant-PGC1A and DGAT1 inhibition conditions were restored by allelic deletion of Ncoa7 (**Fig. 5J-L, fig. S15G-K**).

These findings indicate that, during glucose deprivation, DGAT1 inhibition disrupts lysosomal function and cytosolic pH control via lipid-derived damage, impairing transcriptional plasticity. Evolutionary changes in IDR residues linked to charge influence adaptations to pH in nutrient deficit, while reducing NCOA7 preserves lysosome function and transcriptional plasticity (**fig. S15L**).

### 8. Species-specific mHAR activity in adipocytes during nutrient stress

After establishing the molecular mechanisms in murine systems, we next asked whether human cells exhibit similar or enhanced responses, given that mHARs contain human-specific mutations. In human primary adipocytes and similar to murine models, glucose deprivation led to cytosolic pH variation, PGC1A binding to mHAR hub genes, mHAR gene expression, and mHAR hub chromatin contacts (**Fig. 6A-D, fig. S16A, B**). In glucose deficit, Pgc1a knockdown blunted fatty acid esterification and led to increased oxidative stress (**Fig. 6E**). Pgc1a and Yy1 silencing reduced long-and short-range mHAR chromatin contacts (**fig. S16C, D**). Silencing top-regulated mHAR hub genes increased fatty acid levels and oxidative stress, while reducing TGs, with these effects exacerbated by etomoxir (mitochondrial CPT1 inhibitor; **fig. S16E, F**).

**Fig. 6.**
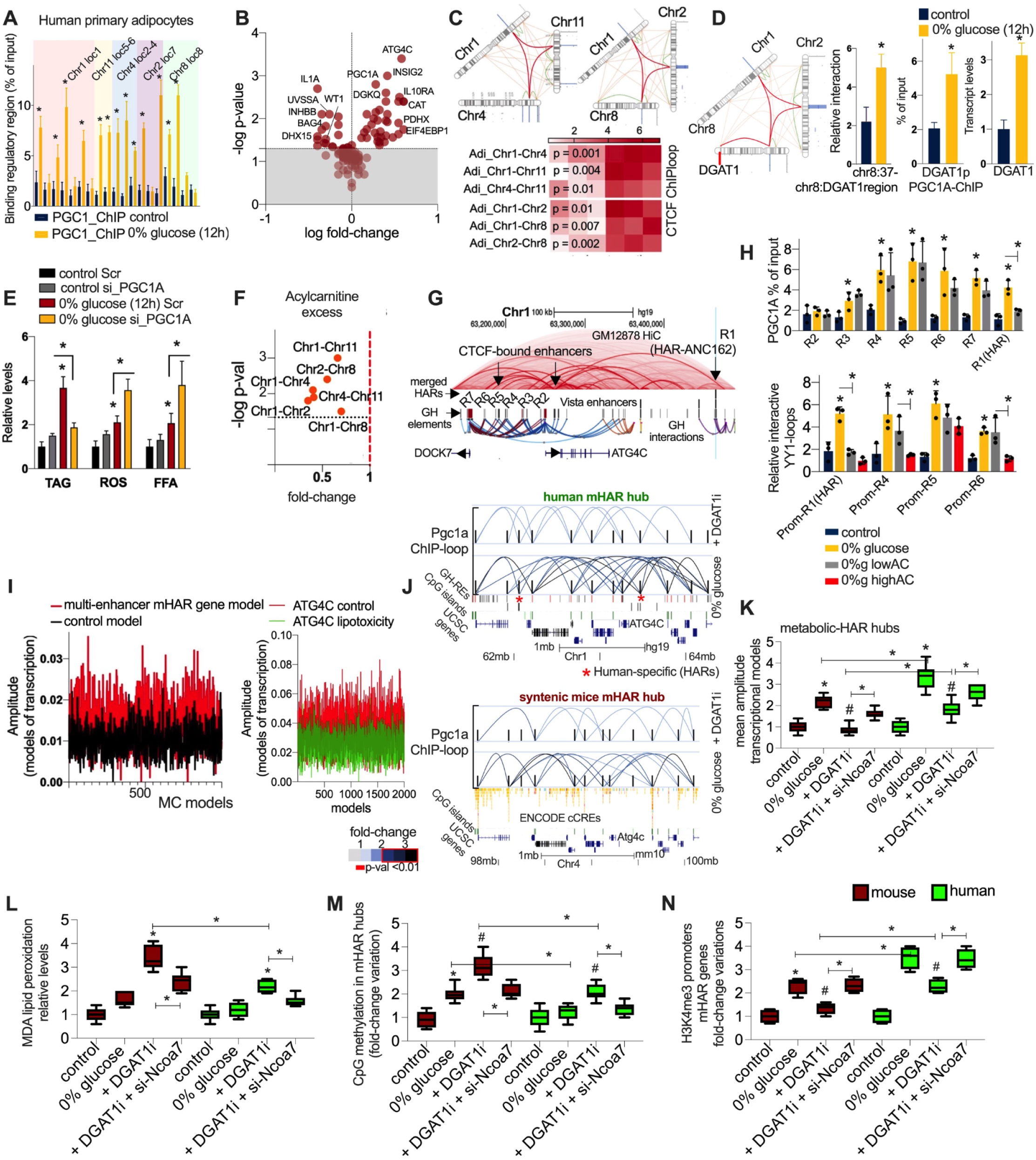
Species-specific mHAR activity in nutrient stress. **(A)** ChIP-qPCR PGC1A binding to HAR elements within mHAR-associated nuclear compartment in human primary adipocytes after 12 h 0% glucose. **(B)** Expression of mHAR genes within nuclear compartments as in A**. (C)** The top panel shows an illustration of the mHAR-associated nuclear compartments. Lower panel shows CTCF-mediated ChIP-loop of mHAR hub interactions in cells as described in A. **(D)** DGAT1-associated nuclear compartment (left panel). From left to right, CTCF-mediated ChIP-loop homotypic interaction in chr8 (mHAR region to DGAT1 location), PGC1A binding to DGAT1 promoter, and DGAT1 expression levels. All assays were done as in A. **(E)** Triglycerides (TAG), reactive oxygen species (ROS), and free fatty acid (FFA) assays in human adipocytes after 0% glucose (12h) and with siRNA for PGC1A. **(F)** CTCF-mediated long-range ChIP-loop mHAR genomic hub interactions in human adipocytes after glucose starvation (12h) with acylcarnitine (AC) excess (20μM). **(G)** Human genome track showing the HAR-element, regulatory elements (R1 to R7) and interactions (from GeneHancer), GM12878-Hi-C data, promoter locations, and CTCF-bound enhancers (found at the human active enhancer to interpret regulatory variants database; HACER). **(H)** Top panel, ChIP-qPCR PGC1A binding to regulatory elements (R1 to R7 as described in G) in human primary adipocytes in glucose deprivation ± low concentration of acylcarnitines (AC; 1μM). Lower panel, YY1 ChIP-loop enhancer-promoter interactions in the same conditions and with high AC concentration (10μM). **(I)** Amplitude of Monte Carlo (MC) models from dynamic gene expression for Atg4c (promoter with multi-enhancer interactions) and control gene (promoter without enhancer interactions) variation in human primary adipocytes after 6 hours of glucose deprivation. Right panel shows the amplitude of transcriptional models for ATG4C dynamic variation in glucose deprivation (6h) and together with high concentration of ACs (20μM). **(J)** PGC1A ChIP-loop in mHAR genomic hub from human (top panel) and mice cells (lower panel; each on top of human and murine genome tracks showing genehancer regions, CpG islands, and gene loci). Contacts (from genehancer regions) from primary white adipocytes after glucose starvation (12h) with or without DGAT1-inhibition (1uM). Contacts are represented as fold-change over control comparison, FC>2 with p-value <0.01. **(K)** Human and murine primary adipocytes as in J, and with DGAT1-inhibition combined with siRNA for Ncoa7 (scramble control in other conditions). Mean amplitude of MC transcriptional models based on dynamic gene expression of mHAR-genes. **(L)** MDA lipid peroxidation levels as in K. **(M)** CpG island methylation assays within the mHAR genomic hub in cells as in K. **(N)** H3K4me3 epigenetic fold-change binding by Chip-qPCR to mHAR gene promoters in cells as in K. Cell experiments were done with 3 independent replicates. Data show mean values and SEM. Unpaired, two-tailed student’s t-test was used when two groups were compared, and ANOVA followed by fisher’s least significant difference (LSD) test for post hoc comparisons for multiple groups. * p-value <0.05 when indicated and when compared to the control group. # p-value <0.05 when compared to 0% glucose control in panels K to N.

Given that FFA accumulation is associated with increased acylcarnitines (ACs) levels, which are linked to metabolic disease (69), we investigated AC excess effects on mHAR genome plasticity. For example, AC excess reduced mHAR hub contacts and PGC1A binding to HAR enhancers (**Fig. 6F, fig. S16G**). During glucose deficit and compared to other hub enhancers linked to the Atg4c promoter (R2-R7), the HAR-element ANC162 (R1) was the only region affected by low concentration of ACs, showing reduced binding of PGC1A and YY1-mediated loops (**Fig. 6G, H**). This suggests that this HAR element might be more exposed during transcriptional organization in low-glucose states and therefore more sensitive to AC accumulation.

This differential sensitivity of HAR elements suggested that multi-enhancer architecture might provide additional regulatory flexibility. Given that multi-enhancer activity influences transcriptional responses (*74*, *75*), we evaluated enhancer interactions and dynamic gene expression of mHAR genes during nutrient stress. Compared to nearby genes, mHAR promoter genes displayed a higher number of YY1-mediated enhancer interactions (coordinates using GeneHancer links; **fig. S16H**). We next compared the dynamic expression of genes with less enhancer contacts (linked to Angptl3) to mHAR genes with multi-enhancer contacts during glucose deprivation (in serial time-frames of 10-15 min for at least one hour; **fig. S16I**). We found increased fold-range variations in mHAR gene expression (**fig. S16I**) compared to active genes with less enhancer linking. To quantify whether multi-enhancer genes exhibit distinctive dynamic properties, we analyzed these fluctuations using nonlinear sine-wave and Monte Carlo modeling (**fig. S16I, J**).

Interestingly, mHAR genes with multi-enhancer contacts displayed transcriptional models with higher amplitude, which were affected by acylcarnitine excess (**Fig. 6I**), suggesting that fatty acid excess impairs transcriptional output in mHAR genes, influencing metabolic responses.

By assessing species-specific adaptations, we found that, despite PGC1A activity in mHAR genome hubs in glucose deprivation, binding to HAR elements (linked to Atg4c, Dgat1, and Pgc1a promoters) was reduced in murine orthologue elements (**fig. S16K**). This was associated with increased mHAR hub local interactions and increased amplitude of transcriptional models, more in human than murine adipocytes. This was associated with reduced transcriptional parameters by DGAT1-inhibition, which were less affected in human cells (**Fig. 6J, K, fig. S16L, M**). This translated into human-specific features, including reduced lipid peroxidation and pH acidification, increased H3K4me3 marks in active mHAR promoters, and decreased CpG DNA methylation in mHAR hubs. In both cellular models, deletion of Ncoa7 restored the deleterious effects of Dgat1 inhibition (**Fig. 6K-N, fig. S16L, N**).

The observed species-specific activity suggests tuning of genomic regions harboring genes that maintain transcriptional plasticity and metabolic homeostasis. The enhanced amplitude of transcriptional responses further suggests evolutionary tuning of the key machinery, with human cells extracting greater organizational gains from equivalent nutrient stress, consistent with the amplified fasting endurance observed at the organismal level.

### 9. Local disease variants are repressed by multi-enhancer regulation of mHAR genes

The enhanced activity of mHAR genes in human cells suggested they might provide protection against metabolic disease, potentially by suppressing pathogenic gene expression nearby. We focused on two representative hubs: the chr1 Atg4c-hub containing HAR element ANC162 near disease-associated gene Angptl3, and the chr8 Dgat1-hub containing HAR element HACNS71 near disease variants linked to lipid metabolism. Our goal was to determine whether disrupting mHAR elements would de-repress these disease genes, and whether this could be reversed by restoring PGC1A function.

To investigate how multi-enhancer mHAR activity impacts local disease-associated variants and genes, we used human cell models and genome editing of HAR elements and nearby disease-associated elements in active and interacting hubs (chr 1 and chr 8). In the chr1 Atg4c-hub, for example, we mutated the active HAR element ANC162 in a YY1-TFBS showing a single nucleotide polymorphism (rs171124210, **Fig. 7A, fig. S17A**). This HAR element is associated with a multi-enhancer cluster (regions R1 to R7; **fig. S17A**) linked to Atg4c gene regulation and to a CTCF-bound element R5 (identified using the HACER database (*76*)) strongly associated with lipid disease (rs12130333, linked gene locus Angptl3, **Fig. 7A, fig. S17B**). CTCF-binding sites in the R5-enhancer near the disease variant were identified as previously shown (*77*), and mutated using CRISPR/Cas9 (**fig. S17B**). In the chr 8 Dgat1-hub, we mapped and mutated CTCF-binding sites in the active HAR-HACNS71 element (**fig. S17C**) and in a nearby region exhibiting a disease variant linked to lipid metabolism as well (rs55831924, **Fig. 7A, fig. S17D**, see supplementary text).

**Fig. 7.**
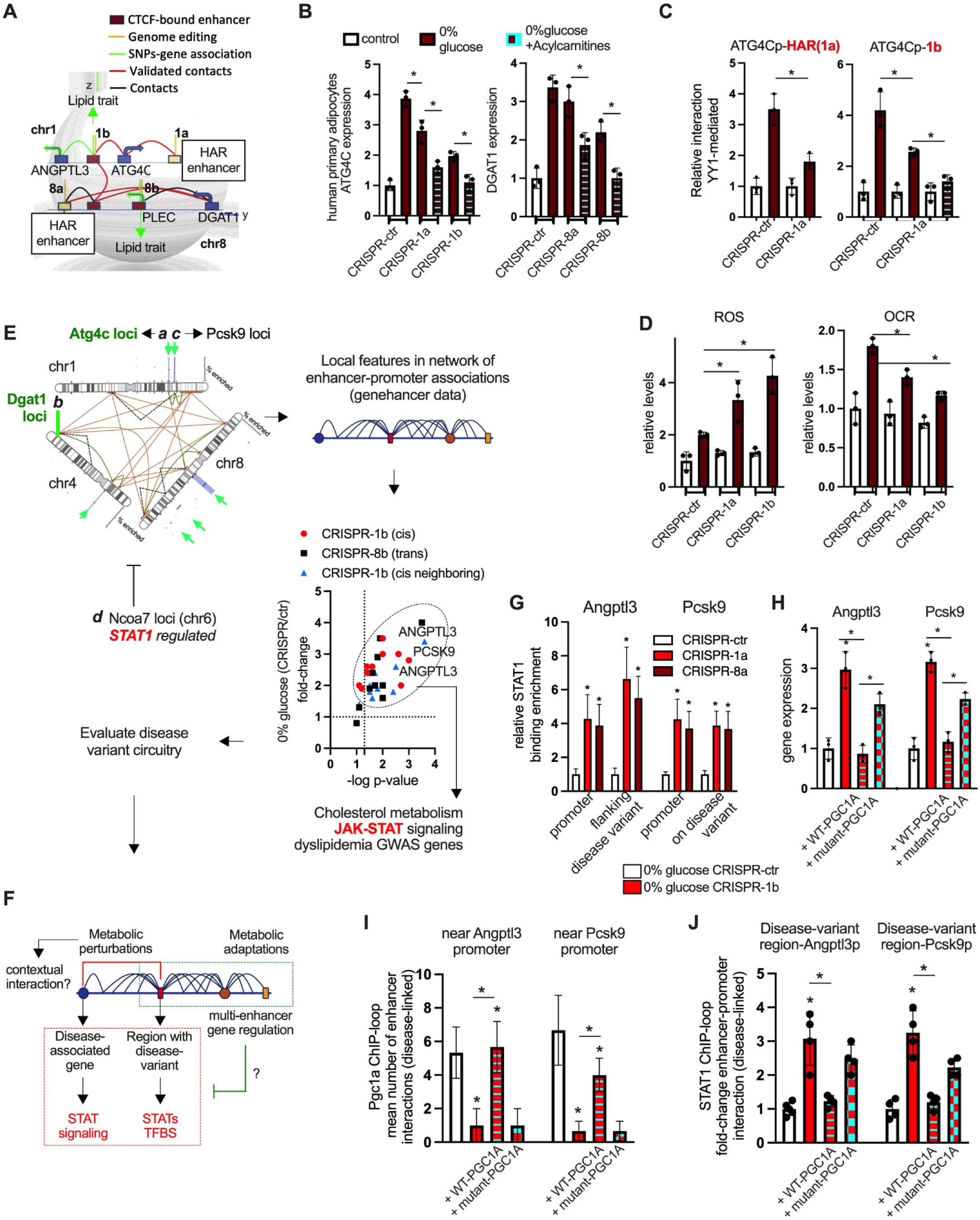
Dissection of mHAR disease circuitry and cooperative hijacking of disease variants and genes. **(A)** Illustration showing local dissection of structural dependencies between mHAR ATG4C and DGAT1 genomic hubs, location of related GWAS variants with associated genes, regulatory elements, and coordinates (1a, 1b, 8a and 8b) for genome editing with CRISPR/Cas9 system. **(B)** ATG4C (left panel) and DGAT1 (right panel) gene expression in human adipocytes displaying targeted mutations as described in A after glucose starvation or together with acylcarnitine (10uM). **(C)** Left panel shows relative YY1-mediated loops (ChIP-loop) between HAR-element and ATG4C_promoter in control mutation and edited YY1 binding site in HAR element human primary adipocytes after glucose starvation (12h). Right panel shows enhancer 1b-ATG4C_promoter loops in the same conditions (control mutation and HAR-mutation) and with acylcarnitine (10uM). **(D)** Left panel shows oxidative stress and the right panel shows oxygen consumption rate functional assays in the same conditions. **(E)** Schematic illustration for the evaluation of disease-variant circuitry. mHAR associated nuclear compartment shows functional regions (a to d) linked to molecular adaptations to nutrient stress and associated with disease variants. Dissection of local features and screening of perturbations. **(F)** Summary illustration of disease variant circuitry. **(G)** STAT1 binding enrichment by ChIP-qPCRs on selected disease-associated regions, either flanking or on top of disease variants in genome edited cells after glucose deprivation (12h). **(H)** ANGPTL3 and PCSK9 expression in genome edited human adipocytes after glucose starvation (12h), and transfected with wildtype PGC1A or IDR-mutant PGC1A expression vectors (empty plasmids used as control). **(I)** PGC1A ChIP-loop, mean number of local interactions for the enhancer with disease-associated variants in genome edited cells after glucose deprivation (12h) and as described in I. Positive regulatory region interactions have a fold-change over control of (FC) >2 and a p-value <0.01. **(J)** STAT1 ChIP-loop, fold-change enhancer-promoter interaction (promoter-enhancer with disease-associated variant) in the same conditions as in J. Cell experiments were done with 3 independent replicates. Data show mean values and SEM. Unpaired, two-tailed student’s t-test was used when two groups were compared, and ANOVA followed by fisher’s least significant difference (LSD) test for post hoc comparisons for multiple groups; * p-value <0.05..

Genome-edited cells exposed to glucose deficit showed that mutating mHAR enhancers and associated elements impaired Atg4c and Dgat1 expression, which was further reduced when ACs were added (**Fig. 7A, B, fig. S18A**). This also resulted in reduced enhancer-promoter contacts linked to these genes (**Fig. 7C**, **fig. S18B, C**). These mutations impaired the functional response to glucose starvation, resulting in increased FFA, excess cytosolic acidity and ROS, and reduced mitochondrial respiration (**Fig. 7D, fig. S18D, E**). In line with our previous observations, these results suggest that mHAR-linked elements maintain key chromatin interactions via contextual structural regulators during nutrient stress for cell resilience.

To evaluate if specific genes in mHAR modules display distinctive enhancer-promoter linking features, we used our network-based approach to analyze GeneHancer data, followed by assessing their activity in genome-edited cells. We evaluated disease-associated genes locally, and in the neighboring hub where Pcsk9 is located, as it is strongly associated with cholesterol dysregulation (**Fig. 7E, fig. S19A**). From this approach, we found that genes exhibiting a reduced number of enhancer linkings (named here type B non-cooperative elements) increased their expression in CRISPR-perturbed cells during glucose deprivation, including the dyslipidemia-linked genes Angptl3 and Pcsk9 (*78*, *79*) (**Fig. 7E, fig. S19B-D**). Interestingly, these genes also shared ontological features such as JAK-STAT inflammatory signaling, similar to Ncoa7 (**Fig. 7E, F, fig. S19C-D**). This suggests that, during nutrient deficit, impairing mHAR and linked elements by genetic or environmental perturbations increase the expression of disease genes with specific linking features and inflammatory annotation.

In other words, when mHAR cooperative plasticity is compromised, cells lose the ability to translate nutrient-stress-driven organization into adaptive specialization. Notably, beyond a tolerable threshold, this makes the cell respond through the activation of disease-associated genes by STAT. This can be considered a maladaptive response where the effect and amplitude of environmental variation is not specializing but instead disease associated.

Supporting this, silencing Angptl3 and Pcsk9 in genome edited cells during nutrient stress reduced lipid peroxidation and the expression of Il6 and Tnf, which we found were correlated (**fig. S19E, F**). Detailed evaluation of the mHAR-axis disease variant circuitry (**fig. S21A**) revealed that STATs TFBS were located close to each variant: near the Plec-linked variant (Chr8), flanking the Angptl3- (Chr1) and Ncoa7-linked (Chr6) variants, and directly on the Pcsk9-linked variant (Chr1, **Fig. 7F, fig. S20B-E**). In glucose perturbed and genome edited cells, we observed a significant increase in STAT1 binding in both the promoters and locations near or directly on the disease-associated variants (**Fig. 7G, fig. S20F**). This suggests the existence of a structural and functional regulatory switch where local cooperative plasticity is affected in perturbed fasting adaptations, favoring STAT1 regulation of disease-associated genes.

We next tested if this is reversed by modulating genome plasticity. Transfection with wildtype, but not IDR-mutant PGC1A constructs, reduced Angptl3 and Pcsk9 expression, increased PGC1A-multi-enhancer contacts, and reduced STAT1-mediated loops between disease-linked elements (**Fig. 7H-J**). This reduced inflammatory gene expression and lipid peroxidation in the same conditions (**fig. S21A**). In mice cohorts, Angptl3 and Pcsk9 expression correlated with worse lipid parameters and pH acidification in white fat and liver (**fig. S21B**).

In fasted fat-specific mutant mice, adenoviral delivery of wildtype, but not IDR-mutant PGC1A, reduced lipid peroxidation, improved tissue pH, and reduced Angptl3 and Pcsk9 expression in white fat and liver (despite liver specific targeting; **fig. S21C-G**). Similar to our *in vitro* observations, wildtype but not IDR-mutant PGC1A vector reduced STAT1-mediated loops and was associated with an increased number of PGC1A-mediated enhancer contacts in disease-linked regions, supporting mHAR hub regulation while reducing disease-associated gene expression (**fig. S21H-K**). This suggests that loss of cooperative genome plasticity due to reduced PGC1A activity and mHAR-axis mutations increases STAT1-binding elements near or on disease-associated variants, promoting the expression of genes linked to metabolic dysfunction (**Fig. 8**).

**Fig. 8.**
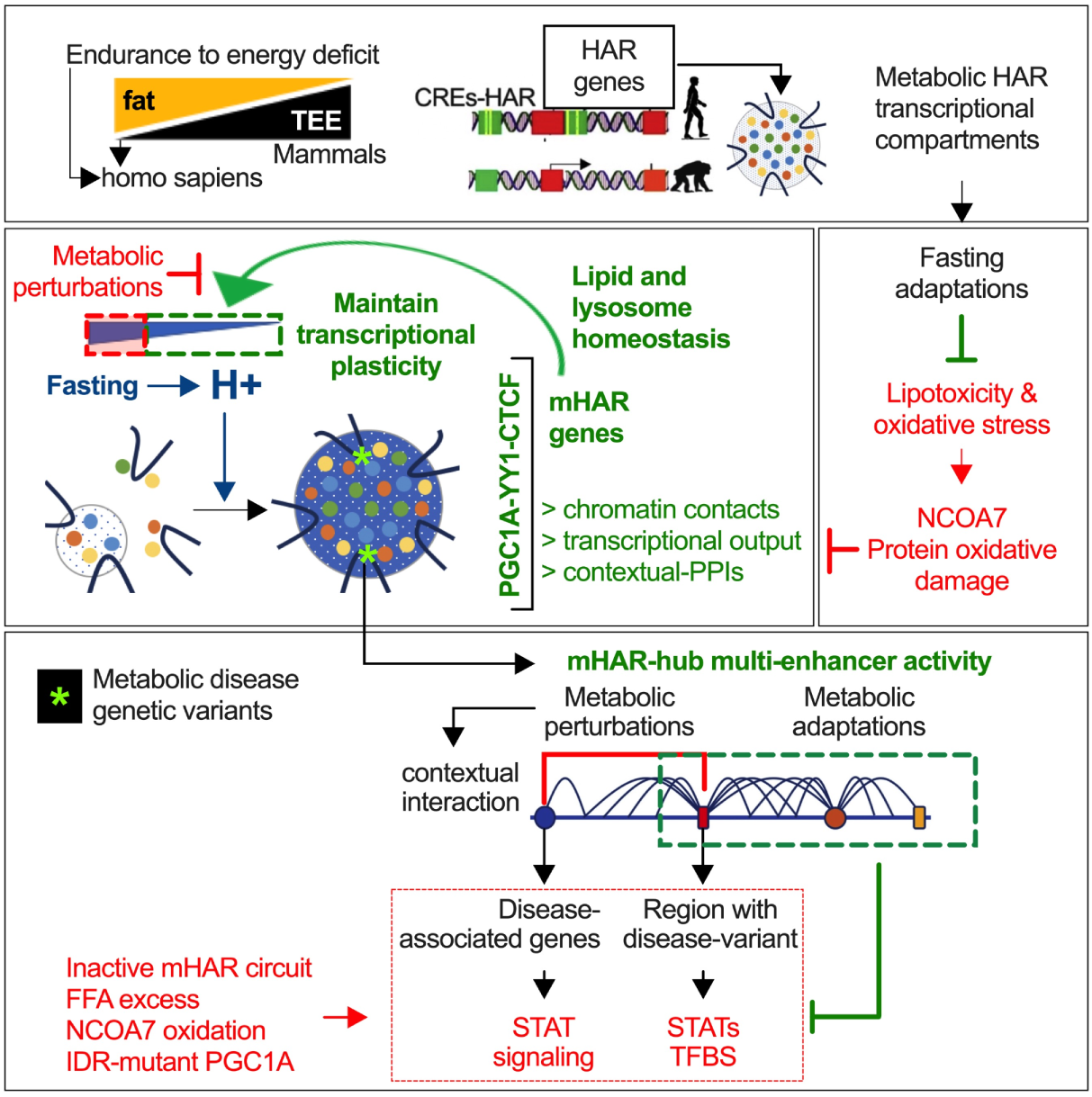
mHAR-associated transcriptional compartments modulate nutrient stress resilience.

## Discussion

Comparative analysis showed a strong association between fat mass and fasting endurance score in mammals. This score is amplified in hominids, leading us to explore molecular links between endurance to energy conservation and evolutionary adaptations, focusing on species-specific mutations such as HARs. Our goal was to identify functional genomic hubs enriched in human-specific mutations, conserved in model organisms, and responsive to nutrient stress.

Our work uncovers molecular links between evolutionary adaptation and dynamic transcriptional organization, identifying a mechanistic chain of adaptation to nutrient stress resilience; evolutionary adaptive regions or key genomic hubs, regulated by cooperative transcriptional regulators, maintain lipid-pH homeostasis to preserve transcriptional plasticity. When this chain is intact, cells convert nutrient fluctuations into durable adaptive states. If this is broken, the same stressor drives inflammation, disease gene activation, and metabolic disease phenotypes. This work shows that genome plasticity is a druggable axis for metabolic resilience leading to the identification of key molecular targets.

### Operational principles of nutrient-driven transcriptional organization

Our functional genomics framework identifies metabolic-HAR genomic hubs, exhibiting cell-type enrichment, transcriptional and genomic relationships, genetic association, and murine phenotype conservation. In model organisms, fasting regulates their activity by cooperative transcriptional regulators, which mediate their chromatin interaction and lead lipid cycling genes regulation. Key genes include, Dgat1 and Atg4c, essential for fasting endurance through the maintenance of lipid-pH homeostasis. When this axis is perturbed genetically or pharmacologically, NCOA7 is found to be a key dysfunction node.

Loss of lipid homeostasis leads to oxidative damage of lysosomal vATPase, mediated by NCOA7 interaction, impairing pH regulation and transcriptional and cellular plasticity. Reducing NCOA7 levels breaks this feedforward cycle, restoring lysosome function, mHAR hub activity, and transcriptional organizations, which enhances both acute and chronic fasting adaptations. This positions NCOA7 as a molecular switch determining whether nutrient stress produces adaptation or dysfunction.

In primary adipocytes, glucose deficit recapitulates mHAR-linked adaptations and promotes cytosolic acidity, influencing the cooperative and structural regulation of mHAR genes involved in maintaining lipid and pH homeostasis. This highlights energy conservation as a feedback integrator that tunes transcriptional efficiency. To explore how cytosolic pH affects transcriptional compartment properties in nutrient stress, we focused on PGC1A (key regulator of key mHAR hubs). IDRs in PGC1A exhibit evolutionary changes in charged residues, and mutating these residues disrupts its ability to respond to nutrient-mediated acidity, affecting its coregulatory plasticity. Cells with IDR-mutant PGC1A or DGAT1 inhibition show impaired adaptation to glucose deficit with reduced lysosomal activity, increased cytosolic acidity and oxidative damage. NCOA7 perturbation rescues these effects, highlighting its role in linking lysosome dysfunction to loss of transcriptional plasticity.

### Evolutionary tuning amplifies human-specific transcriptional responses

In human primary adipocytes, we confirm local structural properties of mHAR hub genes, revealing multi-enhancer contacts and amplified transcriptional responses compared to murine cells. PGC1A binding to HAR elements in human adipocytes is higher than murine orthologue elements, despite local mHAR hub activity. This is associated with increased mHAR gene activity (e.g., Atg4c and Dgat1), reduced lipid peroxidation and cytosolic acidity, along with enhanced epigenetic plasticity. These findings suggest that evolutionary tuning of key transcriptional responses maintains cell plasticity for enhanced species-specific adaptations.

Genome editing of key mHAR hub elements results in impaired adaptations to glucose deficit. These elements maintain local structural contacts that regulate genes modulating nutrient stress resilience. Loss of multi-enhancer mHAR contacts by genome editing is associated with increased STAT1 binding and regulation of neighboring disease-associated variants and genes, such as Angptl3 and Pcsk9. In both cell and mouse models, perturbations in mHAR regulation are rescued by PGC1A but not by IDR-mutant PGC1A. PGC1A reduces the activity of disease circuitry by maintaining multi-enhancer contacts that hijack enhancers with STAT1-driven disease-variants. This reveals a key adaptive logic in which cooperative genome plasticity during energy conservation suppresses pathogenic expression, while its loss enables disease promoting gene activation. The fasting endurance score in hominids may then reflect more than energy storage capacity, for example, enhanced transcriptional buffering against metabolic perturbations.

### Implications for adaptive system design

Our findings show that organizational principles extend beyond metabolism and biological adaptation. Cellular identity is influenced by the genome organization where functional units are transcriptional hubs (interacting chromatin spans) instead of individual regulatory elements or isolated genes. If we consider cells as computational units, this architectural strategy addresses critical computational problems: compression of data by clustering functionally related genes in interacting compartments, rapid retrieval by chromatin contacts and context-dependent outputs where the same elements provide different programs depending on 3D organizational state.

This impacts both biology and computation. In biology, it helps describe why many disease variants are mapped to noncoding regions or seemingly unrelated genes. Instead, disruption of dynamic hub organization might lead to pathogenic adaptations. In computation, it informs design principles in which functional units adapt hierarchically through self-organization, spatial co-localization enhances efficiency and oscillation-driven reorganization transforms environmental variation into computational signal instead of noise.

The dynamic compartments we observe are an example of this strategy. Environmental variations induce reversible assembly in which cooperative factors coordinate remote contacts, creating a form of organizational memory independent of sequence-level modifications. This represents proof that systems may achieve massive parallelism through 3D nested organizational memory exploiting new computational instances by integrating external inputs and evolving architectures through modularity and plasticity. Existing artificial systems are able to change parameters in fixed architectures; biological systems are able to change computational structure itself. By studying how living systems fine-tune and dynamically reorganize to process information, we may uncover design rules applicable to adaptive computation and systems engineering. The nucleus exhibits specific mechanisms for information storage, compression, retrieval, and evolutionary refinement, providing a blueprint for systems that balance stability with plasticity under energetic constraints.

### Limitations

There are a number of limitations that should be considered. Given that our studies focus on adipocytes, additional characterization is needed to determine whether other mHAR hubs lead to similar adaptations under stress. In addition, even though we show that pH tune specific compartment and droplet features, further empirical demonstration should address key biophysical principles for optimal droplet viscosity and transcriptional output. Addressing in vivo and real time pH measurements should provide a better picture of metabolic adaptations. It would also be interesting to extrapolate our disease variant findings to other pathogenic genes and potentially include dynamic integration of this regulatory logic in systematic evaluation of disease variants. Further, the therapeutic potential of NCOA7 modulation needs to be validated in other disease models along with potential off-target effects of this modulation. Finally, our framework needs to be expanded to answer, for example, whether nutrient-stress cycles impact epigenetic memory across mHAR hubs or other higher-order 3D genomic hubs domains coordinating cell identities. Understanding how transient metabolic stressors encode lasting cellular memory can connect acute adaptation to lasting programming effects.

### Concluding perspective

Our study provides a framework linking transcriptional organization, evolutionary adaptation, and metabolic function. Examining the contextual activity of evolutionary adaptations reveals molecular mechanisms connecting specific transcriptional compartments with metabolic endurance. We show that evolutionary tuned genomic hubs convert nutrient fluctuations into adaptive transcriptional states through dynamic nuclear organization. This mechanism operates through NCOA7 as a critical point between adaptation and dysfunction. Human-specific enhancement via mHARs and potentially IDRs in key regulators may underlie to some extent the amplified fasting endurance in hominids, indicating that genome plasticity itself is a crucial target of metabolic selection.

Thus, genome plasticity can be viewed as a druggable axis for metabolic disease, where NCOA7 is a candidate for discerning cellular trajectories towards either pathogenesis or adaptation. Future studies could explore how cooperative transcriptional adaptations impact epigenetic modifications for phenotype diversity and cell memory, leading to lasting cellular changes to environmental variation. More broadly, the notion that cells exploit and not only endure metabolic perturbations to tune functional organization and durable outcomes suggest a general design rule beyond metabolism. These operational rules may inform the engineering of adaptive systems that need robust function with energetic constraints from synthetic biology to machine learning. These findings establish genome plasticity as a key axis for therapeutic discovery and conceptual frameworks for understanding and engineering adaptive resilience.

## Acknowledgements

We thank all the members from the various labs who have contributed to this project. We also thank Amy Grayson for her help reviewing and editing the manuscript.

## Author contributions

Project design and Conceptualization: LZA. Computational Methodology: LZA with feedback from MK, NAP, SKL, KG. Experimental design and validation: LZA, TF, SKL, AG, KG. Supervision of the work: LZA, MK, TF. Funding acquisition: LZA, MK, TF. Writing of original draft: LZA. Writing & editing: LZA, TF. All authors reviewed the manuscript.

## Data and materials availability

Data is available in the supplementary materials.

## Supplementary Materials

Figures S1-21

Tables S1-S12

Materials and methods

## References and Notes

1. C. R. White, L. A. Alton, C. L. Bywater, E. J. Lombardi, D. J. Marshall, Metabolic scaling is the product of life-history optimization. Science. 377, 834–839 (2022).

2. C. R. White, D. J. Marshall, L. A. Alton, P. A. Arnold, J. E. Beaman, C. L. Bywater, C. Condon, T. S. Crispin, A. Janetzki, E. Pirtle, H. S. Winwood-Smith, M. J. Angilletta Jr, S. F. Chenoweth, C. E. Franklin, L. G. Halsey, M. R. Kearney, S. J. Portugal, D. Ortiz-Barrientos, The origin and maintenance of metabolic allometry in animals. Nat Ecol Evol. 3, 598–603 (2019).

3. H. Pontzer, D. A. Raichlen, A. D. Gordon, K. K. Schroepfer-Walker, B. Hare, M. C. O’Neill, K. M. Muldoon, H. M. Dunsworth, B. M. Wood, K. Isler, J. Burkart, M. Irwin, R. W. Shumaker, E. V. Lonsdorf, S. R. Ross, Primate energy expenditure and life history. Proc. Natl. Acad. Sci. U. S. A. 111, 1433–1437 (2014).

4. S. L. Lindstedt, M. S. Boyce, Seasonality, Fasting Endurance, and Body Size in Mammals. The American Naturalist. 125 (1985), pp. 873–878.

5. H. Pontzer, M. H. Brown, D. A. Raichlen, H. Dunsworth, B. Hare, K. Walker, A. Luke, L. R. Dugas, R. Durazo-Arvizu, D. Schoeller, J. Plange-Rhule, P. Bovet, T. E. Forrester, E. V. Lambert, M. E. Thompson, R. W. Shumaker, S. R. Ross, Metabolic acceleration and the evolution of human brain size and life history. Nature. 533, 390–392 (2016).

6. A. Navarrete, C. P. van Schaik, K. Isler, Energetics and the evolution of human brain size. Nature. 480, 91–93 (2011).

7. L. López-Maury, S. Marguerat, J. Bähler, Tuning gene expression to changing environments: from rapid responses to evolutionary adaptation. Nat. Rev. Genet. 9, 583–593 (2008).

8. K. S. Pollard, S. R. Salama, N. Lambert, M.-A. Lambot, S. Coppens, J. S. Pedersen, S. Katzman, B. King, C. Onodera, A. Siepel, A. D. Kern, C. Dehay, H. Igel, M. Ares Jr, P. Vanderhaeghen, D. Haussler, An RNA gene expressed during cortical development evolved rapidly in humans. Nature. 443, 167–172 (2006).

9. J. A. Capra, G. D. Erwin, G. McKinsey, J. L. R. Rubenstein, K. S. Pollard, Many human accelerated regions are developmental enhancers. Philos. Trans. R. Soc. Lond. B Biol. Sci. 368, 20130025 (2013).

10. E. H. Finn, T. Misteli, Molecular basis and biological function of variability in spatial genome organization. Science. 365 (2019), doi:10.1126/science.aaw9498.

11. T. Misteli, The Self-Organizing Genome: Principles of Genome Architecture and Function. Cell. 183, 28–45 (2020).

12. C. H. Eskiw, N. F. Cope, I. Clay, S. Schoenfelder, T. Nagano, P. Fraser, Transcription factories and nuclear organization of the genome. Cold Spring Harb. Symp. Quant. Biol. 75, 501–506 (2010).

13. D. Hnisz, K. Shrinivas, R. A. Young, A. K. Chakraborty, P. A. Sharp, A Phase Separation Model for Transcriptional Control. Cell. 169, 13–23 (2017).

14. B. R. Sabari, A. Dall’Agnese, A. Boija, I. A. Klein, E. L. Coffey, K. Shrinivas, B. J. Abraham, N. M. Hannett, A. V. Zamudio, J. C. Manteiga, C. H. Li, Y. E. Guo, D. S. Day, J. Schuijers, E. Vasile, S. Malik, D. Hnisz, T. I. Lee, I. I. Cisse, R. G. Roeder, P. A. Sharp, A. K. Chakraborty, R. A. Young, Coactivator condensation at super-enhancers links phase separation and gene control. Science. 361 (2018), doi:10.1126/science.aar3958.

15. A. R. Strom, A. V. Emelyanov, M. Mir, D. V. Fyodorov, X. Darzacq, G. H. Karpen, Phase separation drives heterochromatin domain formation. Nature. 547, 241–245 (2017).

16. J. A. Riback, L. Zhu, M. C. Ferrolino, M. Tolbert, D. M. Mitrea, D. W. Sanders, M.-T. Wei, R. W. Kriwacki, C. P. Brangwynne, Composition-dependent thermodynamics of intracellular phase separation. Nature. 581, 209–214 (2020).

17. A. Klosin, F. Oltsch, T. Harmon, A. Honigmann, F. Jülicher, A. A. Hyman, C. Zechner, Phase separation provides a mechanism to reduce noise in cells. Science. 367, 464–468 (2020).

18. J. A. Riback, C. D. Katanski, J. L. Kear-Scott, E. V. Pilipenko, A. E. Rojek, T. R. Sosnick, D. A. Drummond, Stress-Triggered Phase Separation Is an Adaptive, Evolutionarily Tuned Response. Cell. 168, 1028–1040.e19 (2017).

19. A. W. Fritsch, A. F. Diaz-Delgadillo, O. Adame-Arana, C. Hoege, M. Mittasch, M. Kreysing, M. Leaver, A. A. Hyman, F. Jülicher, C. A. Weber, Local thermodynamics governs the formation and dissolution of protein condensates in living cells,, doi:10.1101/2021.02.11.430794.

20. A. Tantos, P. Friedrich, P. Tompa, Cold stability of intrinsically disordered proteins. FEBS Lett. 583, 465–469 (2009).

21. R. P. Joyner, J. H. Tang, J. Helenius, E. Dultz, C. Brune, L. J. Holt, S. Huet, D. J. Müller, K. Weis, A glucose-starvation response regulates the diffusion of macromolecules. Elife. 5 (2016), doi:10.7554/eLife.09376.

22. M. A. Mourão, J. B. Hakim, S. Schnell, Connecting the dots: the effects of macromolecular crowding on cell physiology. Biophys. J. 107, 2761–2766 (2014).

23. L. B. Persson, V. S. Ambati, O. Brandman, Cellular Control of Viscosity Counters Changes in Temperature and Energy Availability. Cell. 183, 1572–1585.e16 (2020).

24. G. Hultqvist, E. Åberg, C. Camilloni, G. N. Sundell, E. Andersson, J. Dogan, C. N. Chi, M. Vendruscolo, P. Jemth, Emergence and evolution of an interaction between intrinsically disordered proteins. Elife. 6 (2017), doi:10.7554/eLife.16059.

25. L. Shenhav, D. Zeevi, Resource conservation manifests in the genetic code. Science. 370, 683–687 (2020).

26. I. E. Eres, K. Luo, C. J. Hsiao, L. E. Blake, Y. Gilad, Reorganization of 3D genome structure may contribute to gene regulatory evolution in primates. PLoS Genet. 15, e1008278 (2019).

27. K. Baar, A. R. Wende, T. E. Jones, M. Marison, L. A. Nolte, M. Chen, D. P. Kelly, J. O. Holloszy, Adaptations of skeletal muscle to exercise: rapid increase in the transcriptional coactivator PGC-1. FASEB J. 16, 1879–1886 (2002).

28. P. Puigserver, Z. Wu, C. W. Park, R. Graves, M. Wright, B. M. Spiegelman, A cold-inducible coactivator of nuclear receptors linked to adaptive thermogenesis. Cell. 92, 829–839 (1998).

29. J. C. Yoon, P. Puigserver, G. Chen, J. Donovan, Z. Wu, J. Rhee, G. Adelmant, J. Stafford, C. R. Kahn, D. K. Granner, C. B. Newgard, B. M. Spiegelman, Control of hepatic gluconeogenesis through the transcriptional coactivator PGC-1. Nature. 413, 131–138 (2001).

30. S. F. Schmidt, S. Mandrup, Gene program-specific regulation of PGC-1{alpha} activity. Genes Dev. 25 (2011), pp. 1453–1458.

31. J. Pérez-Schindler, B. Kohl, K. Schneider-Heieck, V. Adak, J. Delezie, G. Maier, T. Sakoparnig, E. Vargas-Fernández, B. Karrer-Cardel, D. Ritz, A. Schmidt, M. Hondele, S. Hiller, C. Handschin, RNA-bound PGC-1α controls gene expression in liquid-like nuclear condensates,, doi:10.1101/2020.09.23.310623.

32. M. Harms, P. Seale, Brown and beige fat: development, function and therapeutic potential. Nat. Med. 19, 1252–1263 (2013).

33. J. Nedergaard, T. Bengtsson, B. Cannon, New powers of brown fat: fighting the metabolic syndrome. Cell Metab. 13 (2011), pp. 238–240.

34. E. T. Chouchani, L. Kazak, B. M. Spiegelman, New Advances in Adaptive Thermogenesis: UCP1 and Beyond. Cell Metab. 29, 27–37 (2019).

35. S. F. Beer, P. M. Bircham, S. R. Bloom, P. M. Clark, C. N. Hales, C. M. Hughes, C. T. Jones, D. R. Marsh, P. R. Raggatt, A. L. Findlay, The effect of a 72-h fast on plasma levels of pituitary, adrenal, thyroid, pancreatic and gastrointestinal hormones in healthy men and women. J. Endocrinol. 120, 337–350 (1989).

36. S. Kersten, The impact of fasting on adipose tissue metabolism. Biochimica et Biophysica Acta (BBA) - Molecular and Cell Biology of Lipids. 1868 (2023), p. 159262.

37. T. B. Nguyen, S. M. Louie, J. R. Daniele, Q. Tran, A. Dillin, R. Zoncu, D. K. Nomura, J. A. Olzmann, DGAT1-Dependent Lipid Droplet Biogenesis Protects Mitochondrial Function during Starvation-Induced Autophagy. Dev. Cell. 42, 9–21.e5 (2017).

38. C. Chitraju, N. Mejhert, J. T. Haas, L. G. Diaz-Ramirez, C. A. Grueter, J. E. Imbriglio, S. Pinto, S. K. Koliwad, T. C. Walther, R. V. Farese Jr, Triglyceride Synthesis by DGAT1 Protects Adipocytes from Lipid-Induced ER Stress during Lipolysis. Cell Metab. 26, 407–418.e3 (2017).

39. X. X. Yu, D. A. Lewin, W. Forrest, S. H. Adams, Cold elicits the simultaneous induction of fatty acid synthesis and β-oxidation in murine brown adipose tissue: prediction from differential gene expression and confirmation in vivo. The FASEB Journal. 16 (2002), pp. 155–168.

40. E. P. Mottillo, P. Balasubramanian, Y.-H. Lee, C. Weng, E. E. Kershaw, J. G. Granneman, Coupling of lipolysis and de novo lipogenesis in brown, beige, and white adipose tissues during chronic β3-adrenergic receptor activation. J. Lipid Res. 55, 2276–2286 (2014).

41. C. C. Lindsey, BODY SIZES OF POIKILOTHERM VERTEBRATES AT DIFFERENT LATITUDES. Evolution. 20, 456–465 (1966).

42. D. E. Lieberman, D. M. Bramble, The evolution of marathon running : capabilities in humans. Sports Med. 37, 288–290 (2007).

43. G. van Mierlo, O. Pushkarev, J. F. Kribelbauer, B. Deplancke, Chromatin modules and their implication in genomic organization and gene regulation. Trends Genet. 39, 140–153 (2023).

44. K. De Preter, R. Barriot, F. Speleman, J. Vandesompele, Y. Moreau, Positional gene enrichment analysis of gene sets for high-resolution identification of overrepresented chromosomal regions. Nucleic Acids Res. 36, e43 (2008).

45. D. Swain-Lenz, A. Berrio, A. Safi, G. E. Crawford, G. A. Wray, Comparative Analyses of Chromatin Landscape in White Adipose Tissue Suggest Humans May Have Less Beigeing Potential than Other Primates. Genome Biol. Evol. 11, 1997–2008 (2019).

46. S. Berto, I. Mendizabal, N. Usui, K. Toriumi, P. Chatterjee, C. Douglas, C. A. Tamminga, T. M. Preuss, S. V. Yi, G. Konopka, Accelerated evolution of oligodendrocytes in the human brain. Proc. Natl. Acad. Sci. U. S. A. 116, 24334–24342 (2019).

47. C. S. Greene, A. Krishnan, A. K. Wong, E. Ricciotti, R. A. Zelaya, D. S. Himmelstein, R. Zhang, B. M. Hartmann, E. Zaslavsky, S. C. Sealfon, D. I. Chasman, G. A. FitzGerald, K. Dolinski, T. Grosser, O. G. Troyanskaya, Understanding multicellular function and disease with human tissue-specific networks. Nat. Genet. 47, 569–576 (2015).

48. A. D. Schmitt, M. Hu, I. Jung, Z. Xu, Y. Qiu, C. L. Tan, Y. Li, S. Lin, Y. Lin, C. L. Barr, B. Ren, A Compendium of Chromatin Contact Maps Reveals Spatially Active Regions in the Human Genome. Cell Rep. 17, 2042–2059 (2016).

49. S. Kaufmann, C. Fuchs, M. Gonik, E. E. Khrameeva, A. A. Mironov, D. Frishman, Inter-chromosomal contact networks provide insights into Mammalian chromatin organization. PLoS One. 10, e0126125 (2015).

50. S. Fishilevich, R. Nudel, N. Rappaport, R. Hadar, I. Plaschkes, T. Iny Stein, N. Rosen, A. Kohn, M. Twik, M. Safran, D. Lancet, D. Cohen, GeneHancer: genome-wide integration of enhancers and target genes in GeneCards. Database. 2017 (2017), doi:10.1093/database/bax028.

51. C. J. Willer, Y. Li, G. R. Abecasis, METAL: fast and efficient meta-analysis of genomewide association scans. Bioinformatics. 26, 2190–2191 (2010).

52. A. Hafner, A. Boettiger, The spatial organization of transcriptional control. Nat. Rev. Genet. 24, 53–68 (2023).

53. N. A. Leypold, M. R. Speicher, Evolutionary conservation in noncoding genomic regions. Trends in Genetics. 37 (2021), pp. 903–918.

54. M. Kellis, B. Wold, M. P. Snyder, B. E. Bernstein, A. Kundaje, G. K. Marinov, L. D. Ward, E. Birney, G. E. Crawford, J. Dekker, I. Dunham, L. L. Elnitski, P. J. Farnham, E. A. Feingold, M. Gerstein, M. C. Giddings, D. M. Gilbert, T. R. Gingeras, E. D. Green, R. Guigo, T. Hubbard, J. Kent, J. D. Lieb, R. M. Myers, M. J. Pazin, B. Ren, J. A. Stamatoyannopoulos, Z. Weng, K. P. White, R. C. Hardison, Defining functional DNA elements in the human genome. Proc. Natl. Acad. Sci. U. S. A. 111, 6131–6138 (2014).

55. X. Ji, D. B. Dadon, B. J. Abraham, T. I. Lee, R. Jaenisch, J. E. Bradner, R. A. Young, Chromatin proteomic profiling reveals novel proteins associated with histone-marked genomic regions. Proc. Natl. Acad. Sci. U. S. A. 112, 3841–3846 (2015).

56. A. S. Weintraub, C. H. Li, A. V. Zamudio, A. A. Sigova, N. M. Hannett, D. S. Day, B. J. Abraham, M. A. Cohen, B. Nabet, D. L. Buckley, Y. E. Guo, D. Hnisz, R. Jaenisch, J. E. Bradner, N. S. Gray, R. A. Young, YY1 Is a Structural Regulator of Enhancer-Promoter Loops. Cell. 171, 1573–1588.e28 (2017).

57. J. T. Cunningham, J. T. Rodgers, D. H. Arlow, F. Vazquez, V. K. Mootha, P. Puigserver, mTOR controls mitochondrial oxidative function through a YY1-PGC-1alpha transcriptional complex. Nature. 450, 736–740 (2007).

58. A. S. Holehouse, J. Ahad, R. K. Das, R. V. Pappu, CIDER: Classification of Intrinsically Disordered Ensemble Regions. Biophysical Journal. 108 (2015), p. 228a.

59. B. Levine, G. Kroemer, Biological Functions of Autophagy Genes: A Disease Perspective. Cell. 176, 11–42 (2019).

60. R. Singh, S. Kaushik, Y. Wang, Y. Xiang, I. Novak, M. Komatsu, K. Tanaka, A. M. Cuervo, M. J. Czaja, Autophagy regulates lipid metabolism. Nature. 458, 1131–1135 (2009).

61. A. S. Rambold, S. Cohen, J. Lippincott-Schwartz, Fatty acid trafficking in starved cells: regulation by lipid droplet lipolysis, autophagy, and mitochondrial fusion dynamics. Dev. Cell. 32, 678–692 (2015).

62. P. Mitchell, Coupling of phosphorylation to electron and hydrogen transfer by a chemi-osmotic type of mechanism. Nature. 191, 144–148 (1961).

63. J. R. Casey, S. Grinstein, J. Orlowski, Sensors and regulators of intracellular pH. Nat. Rev. Mol. Cell Biol. 11, 50–61 (2010).

64. J. A. Mindell, Lysosomal acidification mechanisms. Annu. Rev. Physiol. 74, 69–86 (2012).

65. J. I. Gutierrez, G. P. Brittingham, Y. Karadeniz, K. D. Tran, A. Dutta, A. S. Holehouse, C. L. Peterson, L. J. Holt, SWI/SNF senses carbon starvation with a pH-sensitive low-complexity sequence. Elife. 11 (2022), doi:10.7554/eLife.70344.

66. B. J. Czowski, R. Romero-Moreno, K. J. Trull, K. A. White, Cancer and pH Dynamics: Transcriptional Regulation, Proteostasis, and the Need for New Molecular Tools. Cancers. 12 (2020), doi:10.3390/cancers12102760.

67. A. Negre-Salvayre, C. Coatrieux, C. Ingueneau, R. Salvayre, Advanced lipid peroxidation end products in oxidative damage to proteins. Potential role in diseases and therapeutic prospects for the inhibitors. Br. J. Pharmacol. 153, 6–20 (2008).

68. M. J. Davies, The oxidative environment and protein damage. Biochimica et Biophysica Acta (BBA) - Proteins and Proteomics. 1703 (2005), pp. 93–109.

69. M. Forgac, Vacuolar ATPases: rotary proton pumps in physiology and pathophysiology. Nat. Rev. Mol. Cell Biol. 8, 917–929 (2007).

70. P. L. Oliver, M. J. Finelli, B. Edwards, E. Bitoun, D. L. Butts, E. B. E. Becker, M. T. Cheeseman, B. Davies, K. E. Davies, Oxr1 is essential for protection against oxidative stress-induced neurodegeneration. PLoS Genet. 7, e1002338 (2011).

71. Y. Zhang, X. Chen, Y. Zhao, M. Ponnusamy, Y. Liu, The role of ubiquitin proteasomal system and autophagy-lysosome pathway in Alzheimer’s disease. Rev. Neurosci. 28, 861–868 (2017).

72. C. Chitraju, T. C. Walther, R. V. Farese Jr, The triglyceride synthesis enzymes DGAT1 and DGAT2 have distinct and overlapping functions in adipocytes. J. Lipid Res. 60, 1112–1120 (2019).

73. H. Ashkenazy, S. Abadi, E. Martz, O. Chay, I. Mayrose, T. Pupko, N. Ben-Tal, ConSurf 2016: an improved methodology to estimate and visualize evolutionary conservation in macromolecules. Nucleic Acids Res. 44, W344–50 (2016).

74. A. J. M. Larsson, P. Johnsson, M. Hagemann-Jensen, L. Hartmanis, O. R. Faridani, B. Reinius, Å. Segerstolpe, C. M. Rivera, B. Ren, R. Sandberg, Genomic encoding of transcriptional burst kinetics. Nature. 565, 251–254 (2019).

75. T. Fukaya, B. Lim, M. Levine, Enhancer Control of Transcriptional Bursting. Cell. 166, 358–368 (2016).

76. J. Wang, X. Dai, L. D. Berry, J. D. Cogan, Q. Liu, Y. Shyr, HACER: an atlas of human active enhancers to interpret regulatory variants. Nucleic Acids Res. 47, D106–D112 (2019).

77. J. D. Ziebarth, A. Bhattacharya, Y. Cui, CTCFBSDB 2.0: a database for CTCF-binding sites and genome organization. Nucleic Acids Res. 41, D188–94 (2013).

78. M. Abifadel, M. Varret, J.-P. Rabès, D. Allard, K. Ouguerram, M. Devillers, C. Cruaud, S. Benjannet, L. Wickham, D. Erlich, A. Derré, L. Villéger, M. Farnier, I. Beucler, E. Bruckert, J. Chambaz, B. Chanu, J.-M. Lecerf, G. Luc, P. Moulin, J. Weissenbach, A. Prat, M. Krempf, C. Junien, N. G. Seidah, C. Boileau, Mutations in PCSK9 cause autosomal dominant hypercholesterolemia. Nat. Genet. 34, 154–156 (2003).

79. J. Wang, M. R. Ban, G. Y. Zou, H. Cao, T. Lin, B. A. Kennedy, S. Anand, S. Yusuf, M. W. Huff, R. L. Pollex, R. A. Hegele, Polygenic determinants of severe hypertriglyceridemia. Hum. Mol. Genet. 17, 2894–2899 (2008).

