## Supplementary materials for "Energy Conservation Drives Adaptive Specialization Through Fine-tuned Organizations"

### 1. Supplementary figures

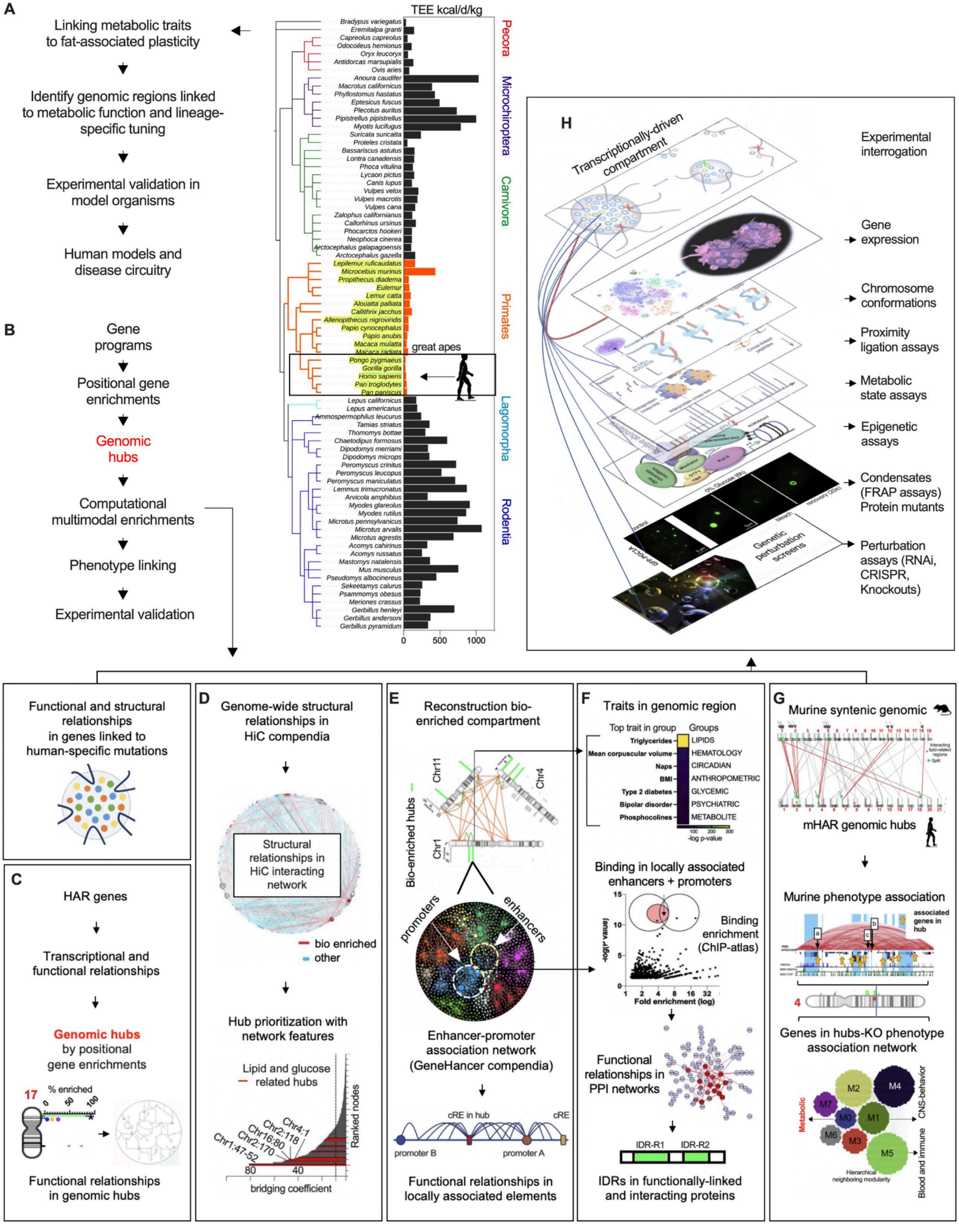

**1.1. Fig. S1. A functional genomics approach to investigate genomic hubs associated with HAR genes** *(related to Fig. 1)*.

**(A)** Illustration of the approach to identify conserved regulatory regions with lineage-specific tuning (e.g., human accelerated regions (HAR)) and linked genes associated with energy metabolism. Right panel shows total energy expenditure (TEE) across mammals (from Pontzer et al. 2014 and 2016)(*1*, *2*).
**(B)** Schematic illustration to investigate genomic hubs from gene programs. Genomic hubs are evaluated computationally across different data modalities to identify molecular dependencies and associated phenotypic adaptations, followed by in-depth experimental validation.
**(C)** Identification of mHAR genomic hubs by positional gene enrichment of HAR-associated genes, followed by hub prioritization by functional enrichments.
**(D)** To evaluate genome structural features, we use colocalization of mHAR genomic hubs (from previous step) with chromatin structural data (e.g. HiC compendia). To identify higher-order structural dependencies, we mapped mHAR genomic hubs on or neighboring interacting chromatin regions (from previously reported HiC data (*3*)[,(4)]), followed by their evaluation through network modeling (of interactions) and prioritization of network features (such as connectivity, compartment forming regions, etc) for regions harboring functionally enriched and metabolic HAR genomic hubs.
**(E)** Illustration of reconstructed bio-enriched compartment following evaluation of chromatin structural data ( homotypic and heterotypic contacts) and 3D modeling of intrachromosomal proximity. Middle and lower panels show genome-wide network reconstruction of enhancer-promoter interactions (using GeneHancer compendia(*5*)), followed by network modeling and features prioritization of genes and regulatory regions within bio-enriched nuclear compartments and mHAR genomic hubs.
**(F)** Regional genetic associations for mHAR genomic hub (Chr1:61-65 mb range) showing top enriched traits within trait-group class using the Common metabolic diseases knowledge portal or HugeAmp portal for human genetic and functional genomic information. Middle panel shows, in genomic hubs of interest and using sub-networks of enhancer-promoter associations, binding of transcriptional regulators by ChIPseq binding enrichment using the ChIPseq atlas portal. Transcriptional regulators are selected by co-binding information (enhancers and promoters), by functional relationships, and by physical interactions using protein-protein interaction data. For selected transcriptional regulators, amino acid sequences are studied computationally to identify disordered regions, conformational features, and conservation.
**(G)** Top panel shows mammalian conservation of mHAR genomic hubs by chromosomal synteny. Lower panel shows identification of phenotype enrichment in genomic hubs using network modeling and analysis of gene-knockout phenotype associations followed by positional gene enrichments of linked genes.
**(H)** Schematic representation showing experimental interrogation of transcriptional nuclear compartments using gene expression, chromosome conformation assays, proximity ligation assays for protein interactions, epigenetic assays, metabolic state assays, condensate assays, and perturbation of key components by RNAi, CRISPR/cas9 system, and murine knockout models.

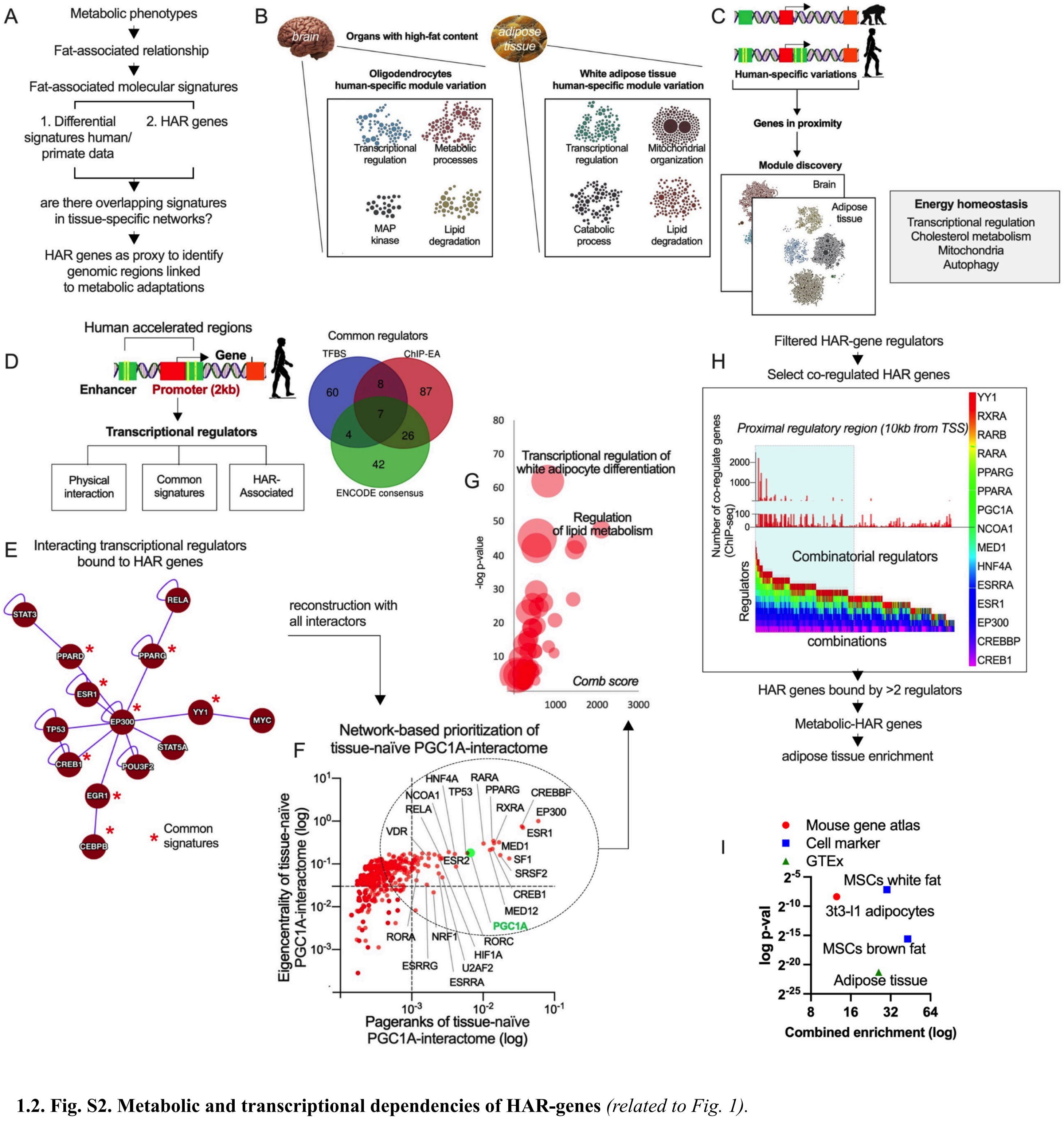

**1.2. Fig. S2. Metabolic and transcriptional dependencies of HAR-genes** *(related to Fig. 1).*

**(A)** Pipeline illustration showing the interrogation of metabolic parameters in mammals, in particular fat-associated functional relationships (the theoretical survival time or fasting endurance) followed by the identification of common molecular signatures across different data sets in tissues of interest using species comparison by high-throughput data, and genes associated with human-accelerated regions (HARs). Common signatures are evaluated in tissue-specific networks(*6*).
**(B)** Network-based discovery of biological processes(*6*) in human enriched gene sets from tissues with high-fat content. Data derived from differential signatures by RNAseq and ATACseq comparing human and primate adipose and brain tissues(*7*, *8*). In boxes, biological ontology terms for enriched modules.
**(C)** HAR-associated genes illustration followed by network functional enrichments as in B. Right panel shows common biological terms.

**(D)** Left panel, illustration showing HAR elements and genes, followed by a schematic approach to identify binding transcriptional regulators (to promoters) and the filtering of regulators by physical interactions (using protein-protein interactions), similar biological function, and HAR-association (proximity to HAR elements). Right panel, venn diagram of found regulators by TFBS-conservation, ENCODE, and ChIP-seq binding enrichment.
**(E)** Network reconstruction of common transcriptional regulators of HAR genes displaying physical interactions and similar biological function.
**(F)** Full reconstruction of interactors (all interactors per regulator) for common regulators and network-based ranking based on connectivity.
**(G)** Functional biological enrichment (using Enrichr analysis tool) for interacting regulators with high connectivity from **F**.
**(H)** Pipeline illustration to identify HAR genes with transcriptional dependencies. Graph panel shows combined binding of PGC1A and interacting coregulators, showing number of co-regulated genes (from ChIP-seq atlas repository) and pairwise combination of bound regulators (selected from network-based ranking as in F). HAR genes bound by more than 2 regulators are selected for functional, tissue- and cell-specific enrichment analysis.
**(I)** Tissue- and cell-type enrichments of mHAR genes (using Enrichr analysis tool).

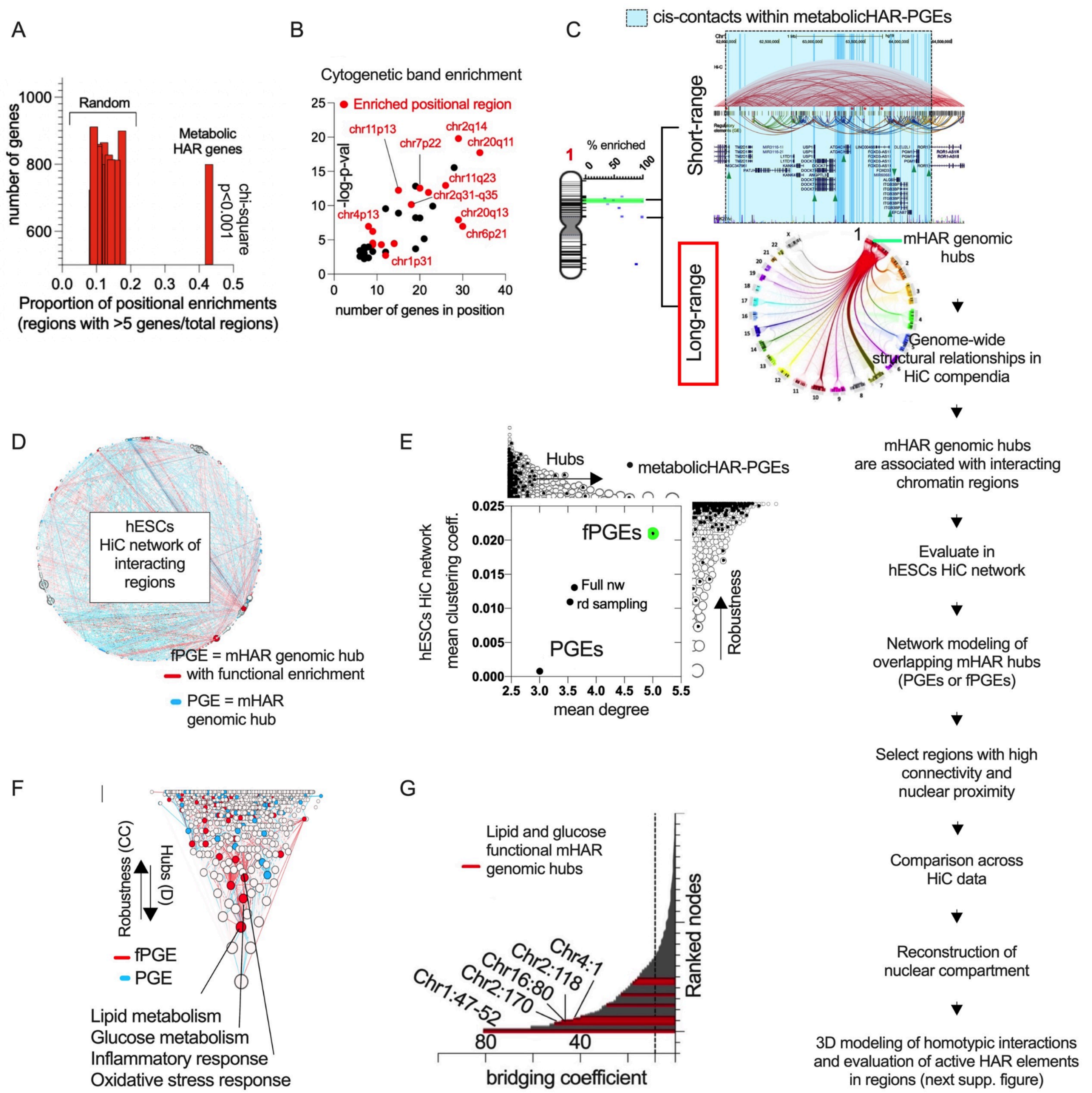

**1.3. Fig. S3. mHAR genomic hubs long-range structural relationships** *(related to Fig. 1)*.

**(A)** Proportion of mHAR genomic hubs by positional gene enrichments (PGEs) of metabolic HAR genes with > 5 genes per genomic hub compared to PGEs from random gene set permutations.
**(B)** Cytogenetic band enrichments for mHAR genes.
**(C)** Pipeline illustration to evaluate genome-wide structural relationships for mHAR genomic hubs. mHAR bio-enriched genomic hub (fPGEs; in green) and other hubs (blue dots) in chromosome 1. Top right panel shows a genome browser picture with short-range (chr1:62-65mb) relationships including range (bleu box), mHAR gene loci (green arrow), HiC local contacts, and enhancer-promoter associations (GeneHancer regions). Lower right panel shows long-range chromatin contacts for chromosome 1, followed by step by step description of integrative analysis.

**(D)** Human embryonic stem cells (hESCs) HiC data and network (from Kaufmann et al. 2015) was used to evaluate mHAR genomic hubs structural dependencies. Network of interacting chromatin regions (circular layout), displaying colocalized mHAR hubs. In red, mHAR hubs with biological enrichment and blue, non-enriched (node size by pagerank).
**(E)** Mean clustering coefficient and node-degree across interacting regions from hESC-HiC network (from D). Graph network features for bio-enriched (in green) and non-enriched hubs (PGEs).
**(F)** Clustering coefficient (robustness) and degree (hubs) hierarchies from hESC-HiC network. mHAR genomic hubs in top-ranked nodes (interacting regions) display metabolic function.
**(G)** Bridging coefficient in hESC-HiC network. In red, top bridging chromatin regions harbouring mHAR genes with lipid and glucose function.
Hypergeometric test enrichments and Chi-square tests were used to determine the statistical significance between expected and observed frequencies.

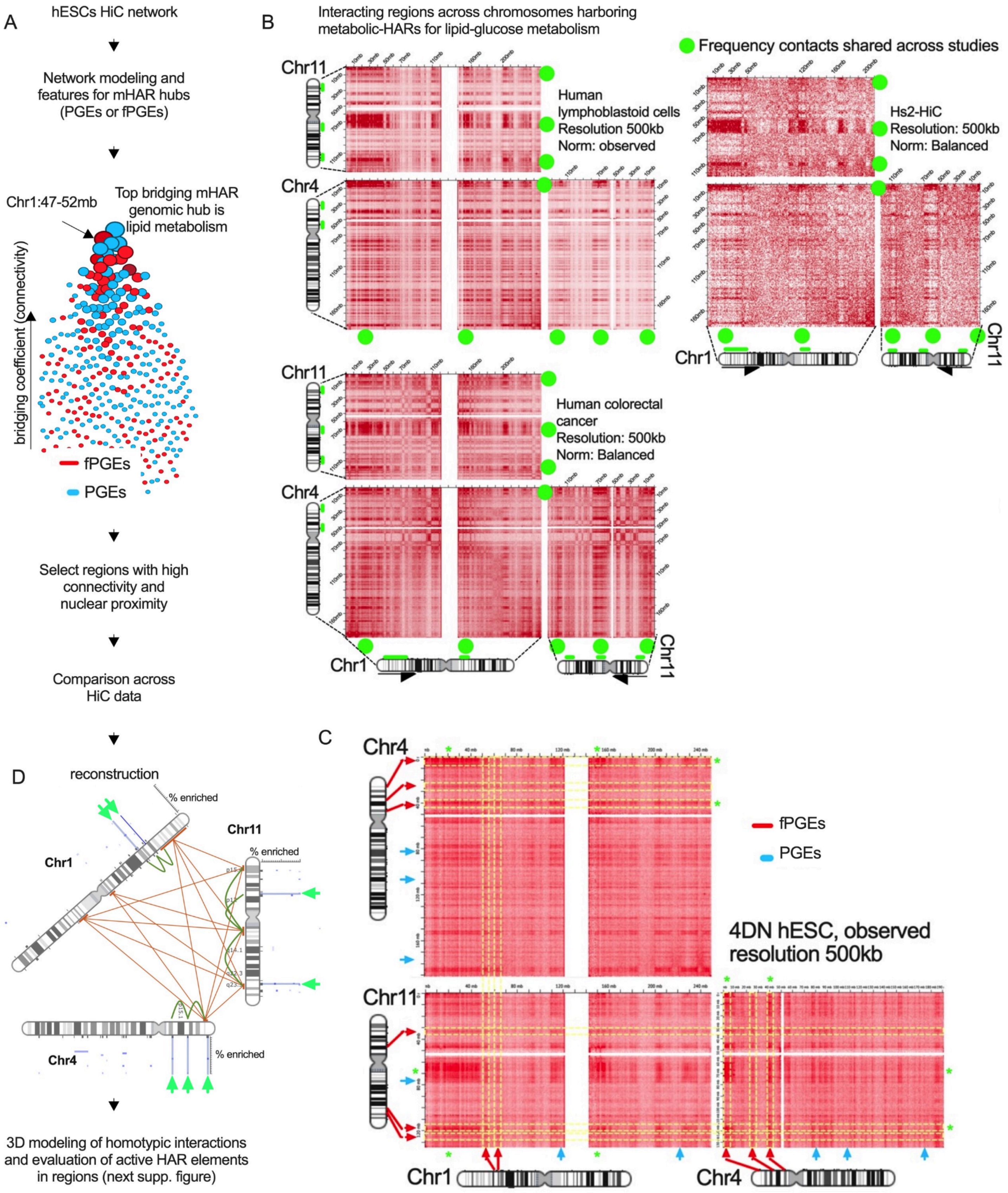
A
hESCs HiC network
Network modeling and features for mHAR hubs (PGEs or fPGEs)
Chr1:47-52mb
Top bridging mHAR genomic hub is lipid metabolism
bridging coefficient (connectivity)
fPGEs
PGEs
Select regions with high connectivity and nuclear proximity
Comparison across HiC data
D
reconstruction
% enriched
Chr1
Chr11
Chr4
3D modeling of homotypic interactions and evaluation of active HAR elements in regions (next supp. figure)
B
Interacting regions across chromosomes harboring metabolic-HARs for lipid-glucose metabolism
Frequency contacts shared across studies
Chr11
Chr4
Chr1
Human lymphoblastoid cells
Resolution 500kb
Norm: observed
Hs2-HiC
Resolution: 500kb
Norm: Balanced
Human colorectal cancer
Resolution: 500kb
Norm: Balanced
C
fPGEs
PGEs
4DN hESC, observed resolution 500kb

##### 1.4. Fig. S4. mHAR genomic hubs long-range structural relationships *(related to Fig. 1)*.

**(A)** Pipeline illustration to evaluate genome-wide structural relationships for mHAR genomic hubs in human embryonic stem cells (hESCs) HiC data. Middle graph panel shows a hierarchical network layout (scaled upwards for bridging score) for interacting mHAR genomic hubs. In red functional mHAR genomic hubs or functional positional gene enrichments (fPGEs), in blue PGEs without biological enrichments.
**(B)** Contact frequency maps across other studies showing interacting chromatin regions (green dots) in chromosomes harboring selected bio-enriched mHAR hubs colocalized in nodes with high connectivity.
**(C)** Contact frequency maps for selected chromosomes harboring mHAR hubs (red, functionally enriched; blue, non-enriched), in relation to heterotypic chromatin contacts.
**(D)** Illustration for reconstructed bio-enriched nuclear compartment harboring mHAR genomic hubs with lipid-glucose related functions. Location of fPGEs in green and PGEs in blue dots. Homotypic and heterotypic interactions.

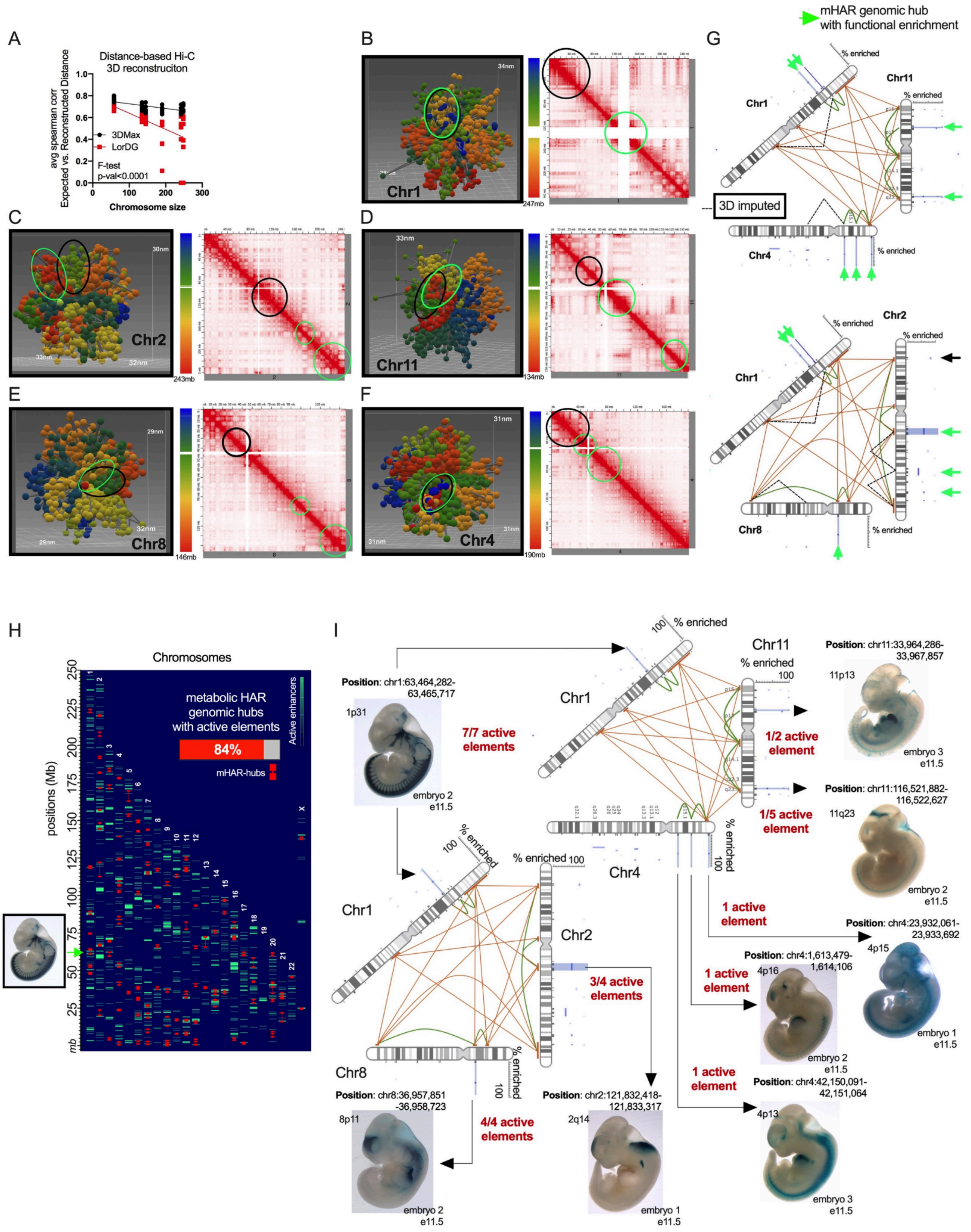
A
Distance-based Hi-C 3D reconstruciton
avg spearman corr Expected vs. Reconstructed Distance
3DMax
LorDG
F-test p-val<0.0001
Chromosome size
B
Chr1
C
Chr2
D
Chr11
E
Chr8
F
Chr4
G
mHAR genomic hub with functional enrichment
% enriched
3D imputed
H
Chromosomes
positions (Mb)
metabolic HAR genomic hubs with active elements
84%
mHAR-hubs
Active enhancers
I
Position: chr1:63,464,282-63,465,717
1p31
7/7 active elements
embryo 2 e11.5
Position: chr11:33,964,286-33,967,857
11p13
1/2 active element
embryo 3 e11.5
Position: chr11:116,521,882-116,522,627
11q23
1/5 active element
embryo 2 e11.5
1 active element
Position: chr4:23,932,061-23,933,692
4p15
embryo 1 e11.5
Position: chr4:1,613,479-1,614,106
4p16
1 active element
embryo 2 e11.5
1 active element
Position: chr4:42,150,091-42,151,064
4p13
embryo 3 e11.5
3/4 active elements
Position: chr8:36,957,851-36,958,723
8p11
4/4 active elements
embryo 2 e11.5
Position: chr2:121,832,418-121,833,317
2q14
embryo 1 e11.5

##### 1.5. Fig. S5. 3D reconstruction and activity within mHAR genomic hubs *(related to Fig. 1)*.

**(A)** Comparison of distance based 3D-HiC reconstruction for chromosome size biased between 3DMax and LorDG softwares. Graph shows expected vs. reconstructed distances related to chromosome size.
For Chr1 **(B)**, Chr2 **(C)**, Chr11 **(D)**, Chr8 **(E)**, and Chr4 **(F),** 3D reconstruction using 3DMax for chromosomes harboring selected mHAR genomic hubs. Black circle in the picture shows interrelated 3D structures (also associated with mHAR genomic hubs) and top homotypic interacting regions (green circle). Blue and green circle in contact maps shows region ranges for associated 3D structures.
**(G)** Illustration for reconstructed bio-enriched nuclear compartment harboring mHAR genomic hubs with lipid-glucose related functions. Location of fPGEs in green and PGEs in blue dots. Homotypic (orange) and heterotypic (green) interactions from contact maps as well as 3D-imputed homotypic interacting regions with a dotted line.
**(H)** Metabolic-HAR genomic hub relationships with active enhancers experimentally validated during embryonic development (archived at the VISTA enhancer repository, Visel et al., 2007). Chromosome landscape map showing positional regions with high density of active enhancers. Percentage of mHAR genomic hubs displaying active elements.
**(I)** Representation of reconstructed compartments displaying number of active elements within metabolic HAR genomic hubs and representative picture of enhancer activity found in the VISTA database.
Linear regression and extra sum-of-squares F test to compare models between conditions.

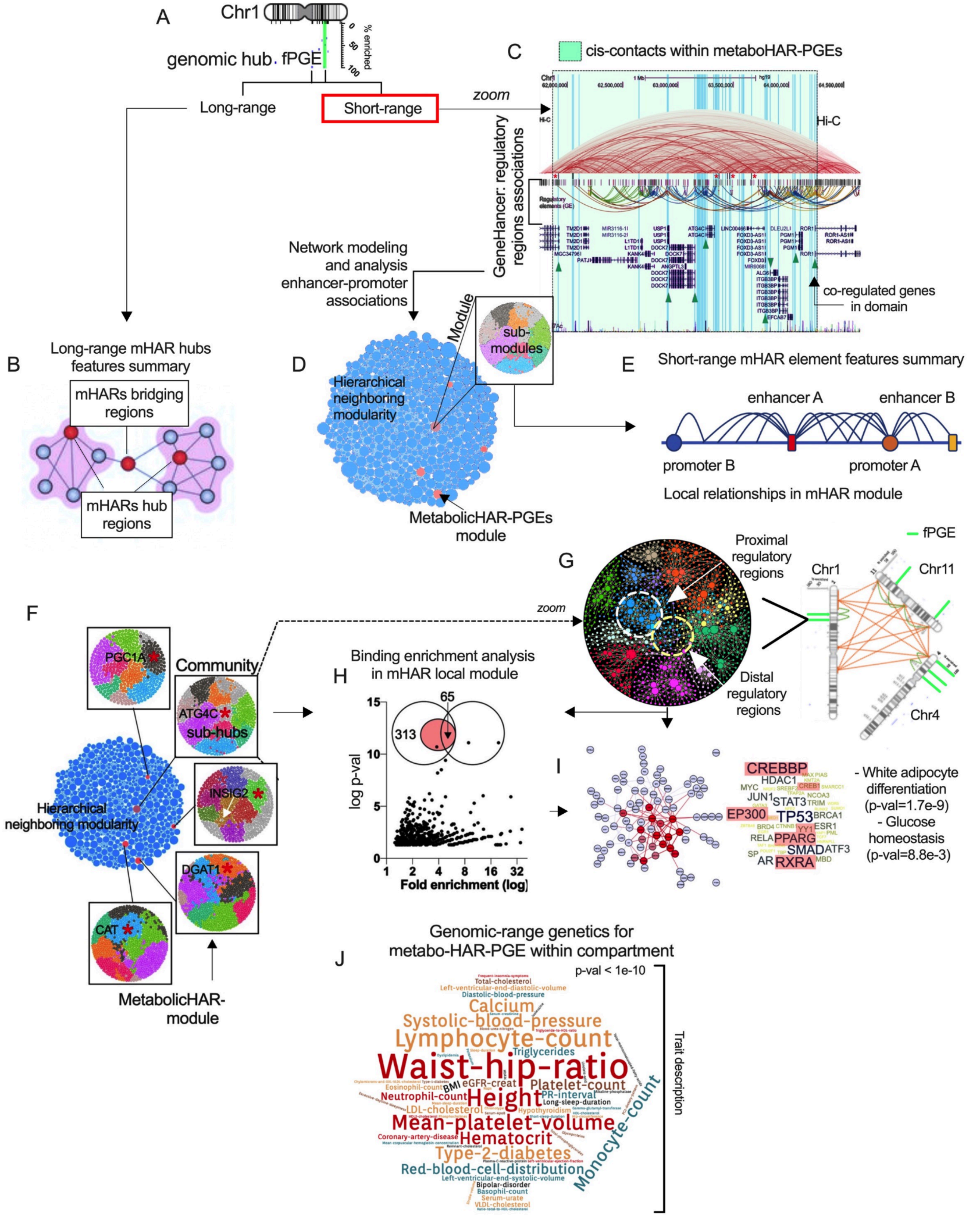
A
Chr1
genomic hub
fPGE
% enriched
Long-range
Short-range
zoom
C
cis-contacts within metaboHAR-PGEs
Hi-C
GeneHancer: regulatory regions associations
co-regulated genes in domain
Network modeling and analysis enhancer-promoter associations
B
Long-range mHAR hubs features summary
mHARs bridging regions
mHARs hub regions
D
Module
sub-modules
Hierarchical neighboring modularity
MetabolicHAR-PGEs module
E
Short-range mHAR element features summary
enhancer A
enhancer B
promoter B
promoter A
Local relationships in mHAR module
F
Community
PGC1A
ATG4C
sub-hubs
INSIG2
DGAT1
CAT
Hierarchical neighboring modularity
MetabolicHAR-module
G
zoom
Proximal regulatory regions
Distal regulatory regions
fPGE
Chr1
Chr11
Chr4
H
Binding enrichment analysis in mHAR local module
log p-val
Fold enrichment (log)
313
65
I
CREBBP
HDAC1
MYC
JUN1
STAT3
EP300
TP53
BRCA1
RELA
PPARG
ESR1
SMAD
ATF3
AR
RXRA
- White adipocyte differentiation (p-val=1.7e-9)
- Glucose homeostasis (p-val=8.8e-3)
J
Genomic-range genetics for metabo-HAR-PGE within compartment
p-val < 1e-10
Trait description
Waist-hip-ratio
Lymphocyte-count
Systolic-blood-pressure
Calcium
Height
Mean-platelet-volume
Hematocrit
Type-2-diabetes
Red-blood-cell-distribution
Monocyte-count
Platelet-count
Triglycerides
Neutrophil-count
LDL-cholesterol
Eosinophil-count
BMI
eGFR-creat
PR-interval
Long-sleep-duration
Total-cholesterol
Diastolic-blood-pressure
Left-ventricular-end-diastolic-volume
Left-ventricular-end-systolic-volume
Hypothyroidism
Coronary-artery-disease
Bipolar-disorder
Basophil-count
Serum-urate
VLDL-cholesterol

##### 1.6. Fig. S6. mHAR genomic hubs short-range structural and disease relationships *(related to Fig. 1).*

**(A)** mHAR bio-enriched genomic hub (fPGEs; in green) in chromosome 1.
**(B)** Long-range chromatin relationships: illustration shows network-based features for mHAR genomic hubs modeled in HiC data
**(C)** Genome browser picture with short-range (chr1:62-65mb) relationships including range (green box), mHAR gene loci regulated by PGC1A and cooperative transcription factors (green arrows), HiC local contacts (source GM12878), and enhancer-promoter associations (GeneHancer repository).
**(D)** Network-mediated interrogation of *cis*-genome-wide relationships among regulatory regions (Fishilevich, S. et al., 2017). Network hierarchical clustering (by modularity) of linking between regulatory elements from Genehancer database, displaying all modules (local neighboring regions), metabolic-HAR modules (in red), and submodule regions.
**(E)** Summary illustration shows network-based features (e.g., multi-enhancer hubs) for regulatory elements from GeneHancer data. Identification of local relationships associated with metabolic HAR elements.
**(F)** Regulatory elements interaction network. In red, metabolic-HAR modules and submodules from mHAR hubs within reconstructed nuclear compartments. Associated HAR genes marked with red asterisk in each submodule.
**(G)** Right panel, mHAR associated nuclear compartments displaying selected hubs (functionally-related in green). Left panel, cis-regulatory network within selected mHAR hub module. Circular layout and pagerank for node size. Dotted circles show local mHAR neighborhoods within the module. Distal regulatory regions refer to enhancers while proximal regions refer to promoters.
**(H)** ChIP-seq enrichment analysis for regulatory regions within the local metabolic-HAR module. Venn diagram showing intersection between regulators bound to both enhancer and promoters.
**(I)** Network of protein-protein interactions for identified bound regulators to both enhancer and promoters within the module. Nodes in red share biological functions. Right panel shows a word-cloud with word size corresponding to the number of interactions. Names in red correspond to PGC1A interacting regulators with top ontological enrichments for interactors.
**(J)** Regional genetic associations for mHAR genomic hubs within selected compartments, displayed as word-cloud based on the repetition of traits-descriptions.

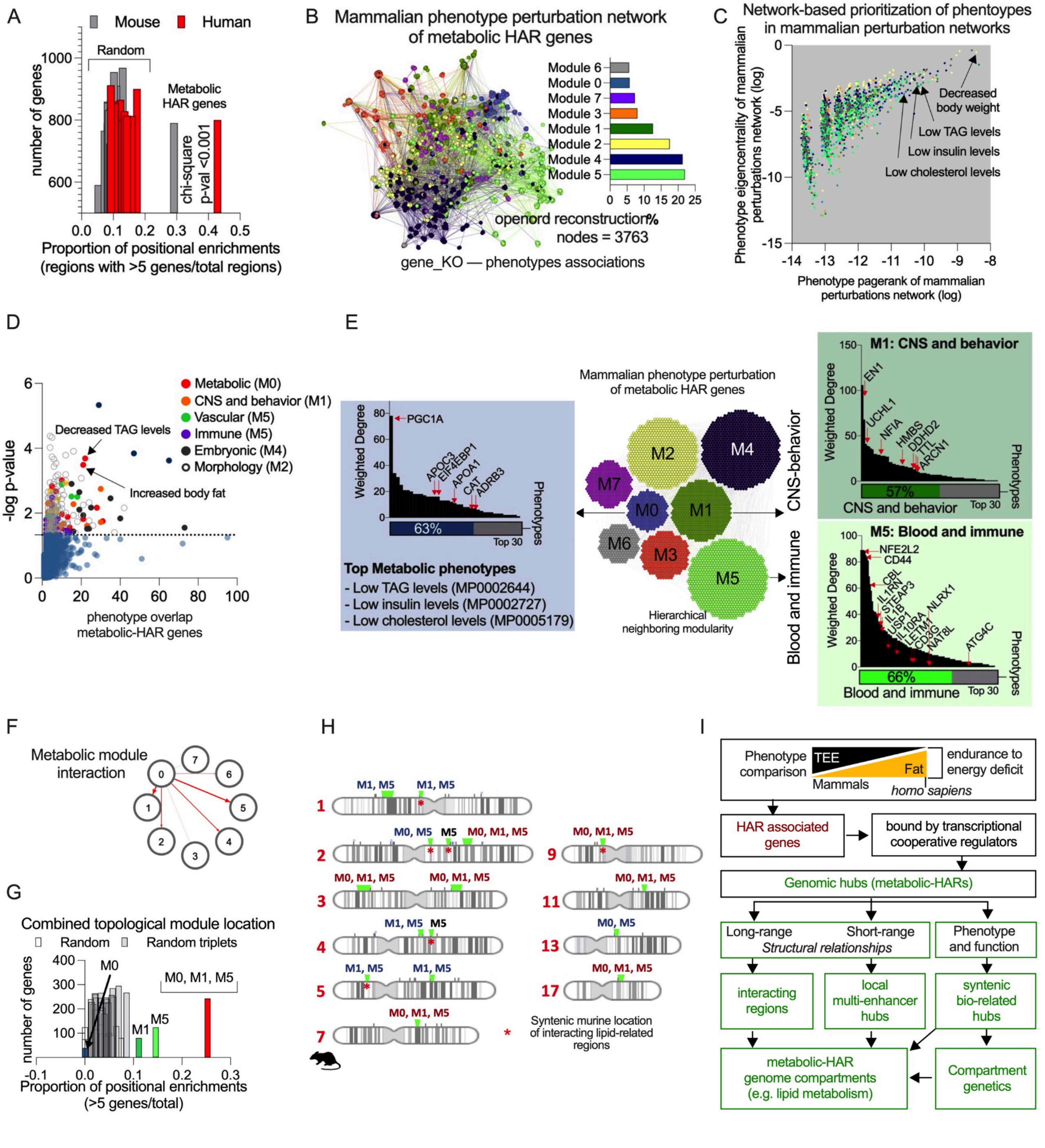

**1.7. Fig. S7. Synteny and phenotypes in mHAR gene knockout models** *(related to Fig. 1)*.

**(A)** Proportion of mHAR genomic hubs by positional gene enrichments (PGEs) of metabolic HAR genes with > 5 genes per genomic hub in both mice and human, and compared to PGEs from random gene set permutations
**(B)** Network integration of murine genetic perturbations (geneKO-phenotype associations) for mHAR genes in openord layout and modularity (right panel).
**(C)** Network analysis and ranking of all murine perturbation relationships for mHAR genes, showing top related phenotypes.
**(D)** Phenotype enrichment for mHAR genes (using Enrichr analysis tool), showing phenotypes associated with metabolic modules in network integration (panel B) as top enrichments.

**(E)** mHAR gene knockout and phenotype association network displayed as a hierarchical modular layout, displaying main phenotypic classes. Sub Panels from arrows show node-degree network analysis of modules harboring mHAR hub genes. Arrows show mHAR genes sharing nuclear proximity in human HiC data.
**(F)** Metabolic inter-module associations, phenotype and genes shared between modules.
**(G)** proportion of PGEs with >5 genes in hubs. In green, hubs formed by genes from each phenotype module. In red, combined positional enrichments. (chi-square test pval<0.001).
**(H)** Share genome hub positions (from G) for related modules (metabolic-related) in murine regions (green). Red asterisks show syntenic regions with nuclear proximity in humans.
**(I)** Summary illustration showing functional relationships for mammalian metabolic phenotypes (top panel) followed by the pipeline evaluation of HAR associated nuclear compartments related to metabolic phenotypes.
Hypergeometric test enrichments and Chi-square tests were used to determine the statistical significance between expected and observed frequencies.

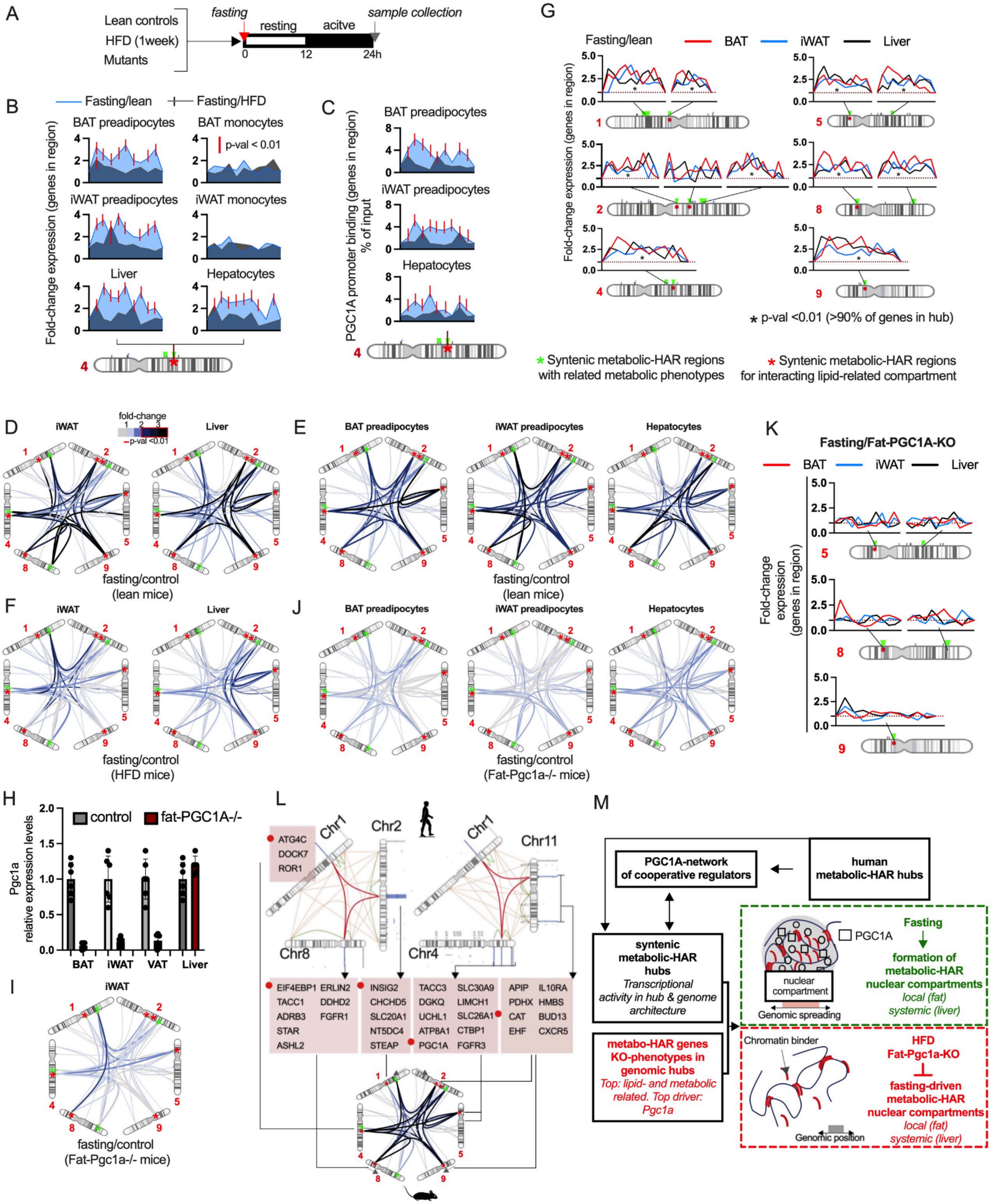
A
Lean controls
HFD (1week)
Mutants
fasting
resting
acitve
sample collection
B
Fasting/lean
Fasting/HFD
BAT preadipocytes
BAT monocytes
p-val < 0.01
iWAT preadipocytes
iWAT monocytes
Liver
Hepatocytes
Fold-change expression (genes in region)
C
PGC1A promoter binding (genes in region) % of input
G
Fasting/lean
BAT
iWAT
Liver
Fold-change expression (genes in region)
* p-val <0.01 (>90% of genes in hub)
* Syntenic metabolic-HAR regions with related metabolic phenotypes
* Syntenic metabolic-HAR regions for interacting lipid-related compartment
D
iWAT
Liver
fold-change
p-val <0.01
fasting/control (lean mice)
E
BAT preadipocytes
iWAT preadipocytes
Hepatocytes
fasting/control (lean mice)
K
Fasting/Fat-PGC1A-KO
BAT
iWAT
Liver
Fold-change expression (genes in region)
F
iWAT
Liver
fasting/control (HFD mice)
J
BAT preadipocytes
iWAT preadipocytes
Hepatocytes
fasting/control (Fat-Pgc1a-/- mice)
H
control
fat-PGC1A-/-
Pgc1a relative expression levels
BAT
iWAT
VAT
Liver
I
iWAT
fasting/control (Fat-Pgc1a-/- mice)
L
Chr1
Chr2
Chr8
Chr1
Chr11
Chr4
ATG4C
DOCK7
ROR1
EIF4EBP1
TACC1
ADRB3
STAR
ASHL2
ERLIN2
DDHD2
FGFR1
INSIG2
CHCHD5
SLC20A1
NT5DC4
STEAP
TACC3
DGKQ
UCHL1
ATP8A1
PGC1A
SLC30A9
LIMCH1
SLC26A1
CTBP1
FGFR3
APIP
PDHX
CAT
EHF
IL10RA
HMBS
BUD13
CXCR5
M
PGC1A-network of cooperative regulators
human metabolic-HAR hubs
syntenic metabolic-HAR hubs
Transcriptional activity in hub & genome architecture
metabo-HAR genes KO-phenotypes in genomic hubs
Top: lipid- and metabolic related. Top driver: Pgc1a
PGC1A
nuclear compartment
Genomic spreading
Fasting
formation of metabolic-HAR nuclear compartments
local (fat)
systemic (liver)
Chromatin binder
Genomic position
HFD
Fat-Pgc1a-KO
fasting-driven metabolic-HAR nuclear compartments
local (fat)
systemic (liver)

**1.8. Fig. S8. PGC1A transcriptional regulation of mHAR-associated compartments** *(related to Fig. 2)*.

**(A)** Illustration showing protocol for fasting metabolic challenge in murine models.
**(B)** Fold change gene expression in metabolic-HAR region (chr4:97-100mb) of hub-forming genes. Each peak represents gene expression variation (arranged following genomic positions left to right), after metabolic challenge in different cells and tissues. Red vertical line represents significant changes between groups p-value<0.01.
**(C)** ChIPqPCR fold-change binding of PGC1A to promoter regions of hub-forming genes described and represented in **B**.
**(D)** Targeted in situ chromosome conformational assays (3C) for metabolic-HAR hub associated nuclear compartments in inguinal fat and liver, as well as in sorted cells from lean wildtype mice (**E**) and HFD-mice **(F)** after a fasting challenge. Targeted contact variations are represented as fold-change over control groups (not fasted littermates). Contacts with FC > 2 have p-value <0.01.
**(G)** Fold-change gene expression after fasting of hub-forming genes in all syntenic metabolic-HAR regions within the associated nuclear compartment. Asterisk shows hub regions with >90% of hub-forming genes with significant fold-change variations.
**(H)** Pgc1a gene expression in tissues from control flox mice and fat-specific PGC1A knockout mice.
**(I)** Targeted in situ chromosome conformational assays (3C) for metabolic-HAR hub associated nuclear compartments in inguinal fat and sorted cells **(J)** from fat-Pgc1a-/- mice after a fasting challenge. Contact variations are represented as fold-change over control groups (not fasted littermates). Contacts with FC > 2 have a p-value <0.01.
**(K)** Fold change gene expression of hub-forming genes in syntenic metabolic-HAR regions within associated nuclear compartment from fat-Pgc1a-/- mice after fasting challenge.
**(L)** Illustration for human mHAR associated nuclear compartment and genomic hubs with some examples of mHAR associated genes (red boxes), and murine syntenic regions forming chromatin contacts and transcriptionally active in fasting.
**(M)** Summary illustration for the identification of syntenic mHAR hubs with similar phenotypes, and validation of compartment regulation by nutrient stress in murine models.
Animal experiments were done with n=5-8 mice per group, both female and male mice were included. Graphs show mean values. Unpaired, two-tailed student's t-test was used when two groups were compared, and ANOVA followed by fisher's least significant difference (LSD) test for post hoc comparisons for multiple groups. * p-value <0.05.

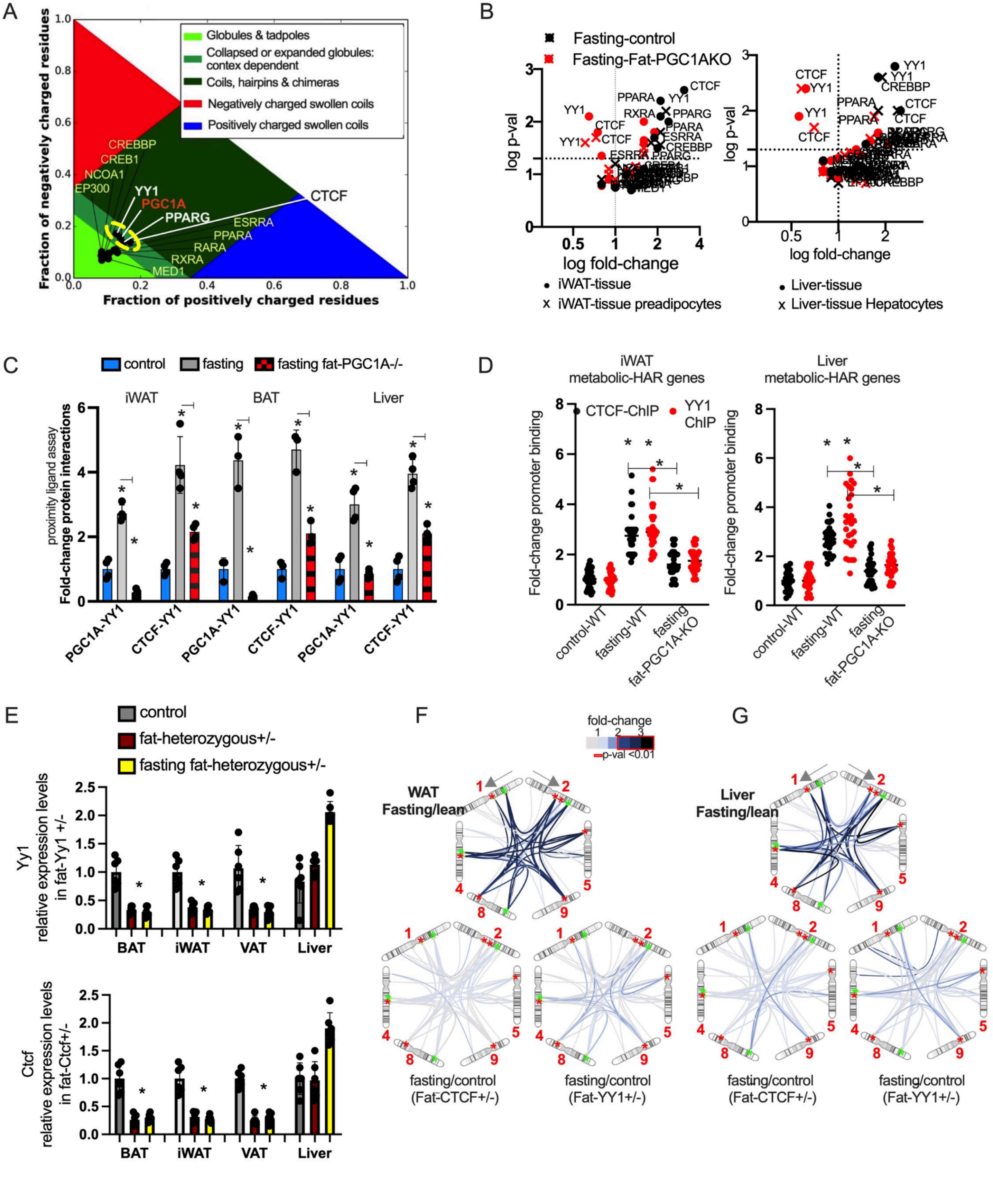
A
Globules & tadpoles
Collapsed or expanded globules: contex dependent
Coils, hairpins & chimeras
Negatively charged swollen coils
Positively charged swollen coils
Fraction of negatively charged residues
Fraction of positively charged residues
CREBBP
CREB1
NCOA1
EP300
YY1
PGC1A
PPARG
CTCF
ESRRA
PPARA
RARA
RXRA
MED1
B
Fasting-control
Fasting-Fat-PGC1AKO
log p-val
log fold-change
iWAT-tissue
iWAT-tissue preadipocytes
Liver-tissue
Liver-tissue Hepatocytes
C
control
fasting
fasting fat-PGC1A-/-
iWAT
BAT
Liver
proximity ligand assay
Fold-change protein interactions
PGC1A-YY1
CTCF-YY1
D
iWAT metabolic-HAR genes
Liver metabolic-HAR genes
CTCF-ChIP
YY1 ChIP
Fold-change promoter binding
control-WT
fasting-WT
fasting fat-PGC1A-KO
E
control
fat-heterozygous+/-
fasting fat-heterozygous+/-
Yy1 relative expression levels in fat-Yy1 +/-
Ctcf relative expression levels in fat-Ctcf+/-
BAT
iWAT
VAT
Liver
F
fold-change
p-val <0.01
WAT Fasting/lean
fasting/control (Fat-CTCF+/-)
fasting/control (Fat-YY1+/-)
G
Liver Fasting/lean
fasting/control (Fat-CTCF+/-)
fasting/control (Fat-YY1+/-)

#### 1.9. Fig. S9. Cooperative transcriptional regulation of mHAR-associated compartments *(related to Fig. 2)*.

**(A)** Diagram for sequence-based classification of disorder scores by CIDER. Different ensemble behavior for PGC1A, cooperative transcription factors, and genome structural factors (CTCF). Light green is associated with droplet formation, while dark green is associated with context-dependant droplet formation (solvation, protein interactions, etc)
**(B)** Gene expression for transcriptional regulators in tissues and sorted cells from control and fat-PGC1A-KO mice after fasting (24h).
**(C)** Proximity ligation assays for transcriptional interactions from tissue extracts from control and fat-PGC1A-KO mice after fasting.
**(D)** ChIP-qPCR binding of YY1 and CTCF to metabolic-HAR genes within the associated nuclear compartment. Each dot represents the mean fold-change variation of their binding to different promoter regions from mice described in **B**.
**(E)** Yy1 and Ctcf gene expression in tissues from control flox mice and fat-specific heterozygous +/- mice and fat-specific heterozygous +/- mice after fasting.
**(F)** Targeted in situ chromosome conformation assays (3C) for metabolic-HAR hubs within the associated nuclear compartment in inguinal fat and liver **(G)** from control and fat-specific YY1+/- and CTCF+/- heterozygous mice after fasting. Contact variations are represented as fold-change over control groups (not fasted littermates). Contacts with FC > 2 have a p-value <0.01.
Animal experiments were done with n=5-8 mice per group, both female and male mice were included. Bars show mean values and SEM. Unpaired, two-tailed student's t-test was used when two groups were compared, and ANOVA followed by fisher's least significant difference (LSD) test for post hoc comparisons for multiple groups. * p-value <0.05.

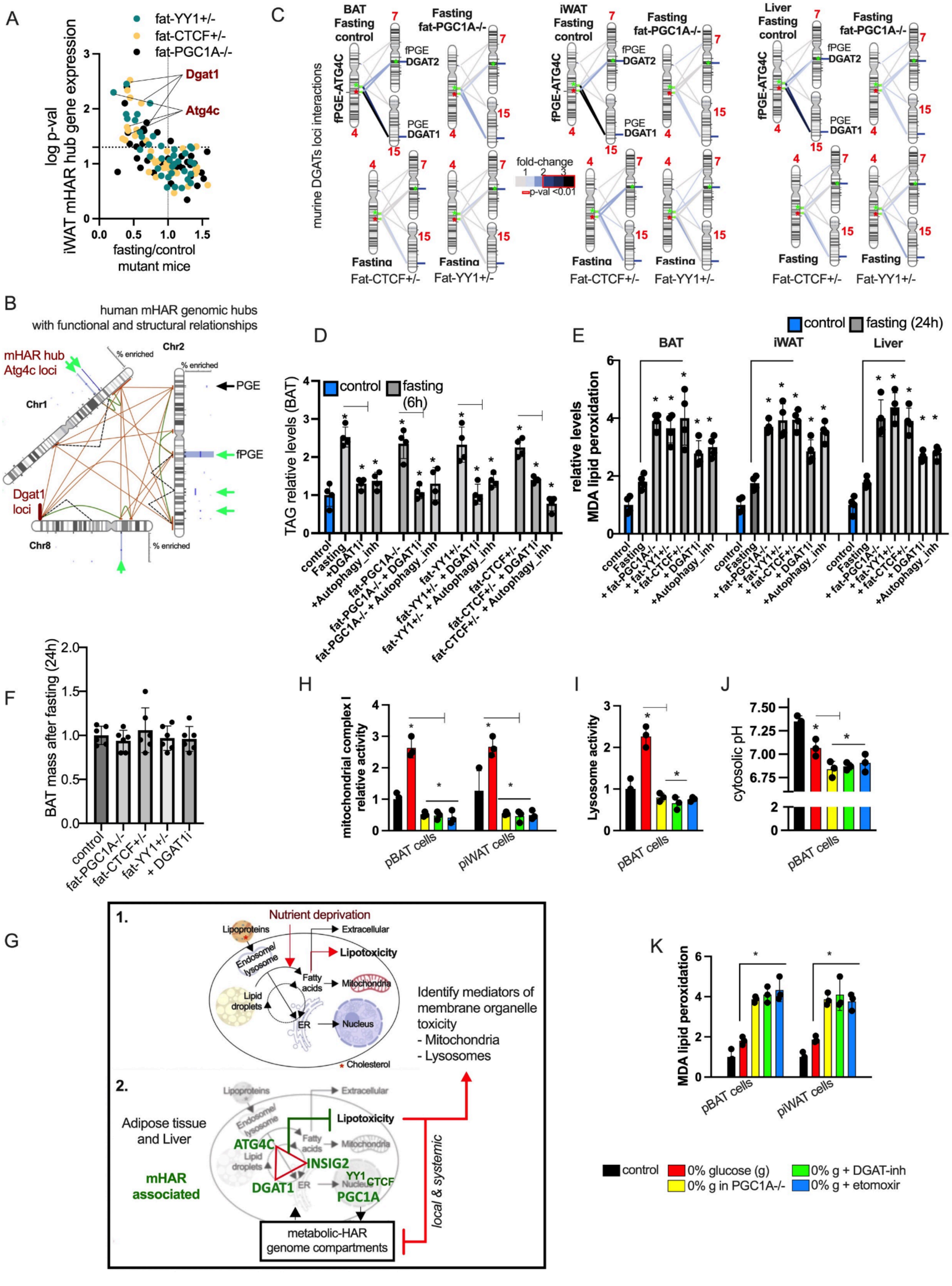
A
fat-YY1+/-
fat-CTCF+/-
fat-PGC1A-/-
Dgat1
Atg4c
log p-val
iWAT mHAR hub gene expression
fasting/control
mutant mice
B
human mHAR genomic hubs
with functional and structural relationships
mHAR hub
Atg4c loci
Chr1
Chr2
Chr8
% enriched
PGE
fPGE
Dgat1
loci
C
murine DGATs loci interactions
BAT
Fasting
control
Fasting
fat-PGC1A-/-
iWAT
Fasting
control
Liver
Fasting
control
fPGE-ATG4C
fPGE
DGAT2
PGE
DGAT1
fold-change
p-val <0.01
Fasting
Fat-CTCF+/-
Fat-YY1+/-
D
TAG relative levels (BAT)
control
fasting
(6h)
control
Fasting
+DGAT1i
+Autophagy_inh
fat-PGC1A-/-
fat-PGC1A-/- + DGAT1i
fat-PGC1A-/- + Autophagy_inh
fat-YY1+/-
fat-YY1+/- + DGAT1i
fat-YY1+/- + Autophagy_inh
fat-CTCF+/-
fat-CTCF+/- + DGAT1i
fat-CTCF+/- + Autophagy_inh
E
control
fasting (24h)
BAT
iWAT
Liver
relative levels
MDA lipid peroxidation
control
Fasting
+ fat-PGC1A-/-
+ fat-YY1+/-
+ fat-CTCF+/-
+ DGAT1i
+Autophagy_inh
F
BAT mass after fasting (24h)
control
fat-PGC1A-/-
fat-CTCF+/-
fat-YY1+/-
+ DGAT1i
H
mitochondrial complex I
relative activity
pBAT cells
piWAT cells
I
Lysosome activity
pBAT cells
J
cytosolic pH
pBAT cells
G
1.
Nutrient deprivation
Lipoproteins
Extracellular
Endosome/
lysosome
Lipotoxicity
Fatty
acids
Mitochondria
Lipid
droplets
ER
Nucleus
Cholesterol
Identify mediators of
membrane organelle
toxicity
- Mitochondria
- Lysosomes
2.
Adipose tissue
and Liver
mHAR
associated
ATG4C
INSIG2
DGAT1
YY1
CTCF
PGC1A
local & systemic
metabolic-HAR
genome compartments
K
MDA lipid peroxidation
pBAT cells
piWAT cells
control
0% glucose (g)
0% g + DGAT-inh
0% g in PGC1A-/-
0% g + etomoxir

##### 1.10. Fig. S10. Phenotypic drivers in mHAR-associated nuclear compartment *(related to Fig. 3)*.

**(A)** Expression of mHAR genes within associated nuclear compartment in inguinal fat from fat-specific mutant mice after fasting followed by functional enrichment of significant genes.
**(B)** Illustration of mHAR associated nuclear compartment and genomic hubs (in green). In red, Atg4c and Dgat1 loci.
**(C)** Targeted in situ chromosome conformation assays (3C) for interactions between mHAR ATG4C-hub and mHAR DGAT1-2 hub in brown fat (BAT), inguinal fat (iWAT), and liver from control and fat-mutant mice after 24 hours fasting. Contact variations are represented as fold-change over control groups (not fasted littermates). Contacts with FC > 2 have a p-value <0.01.
**(D)** Relative triglyceride (TAG or TG) levels in brown fat tissue comparing the response of control and mutant mice to 6h fasting, with or without DGAT1-inhibitor (2 mg/kg, 24 h by i.p.) and autophagy inhibitor (Bafilomycin 1 mg/kg, 24h by i.p.).
**(E)** Malondialdehyde lipid peroxidation assay levels in murine tissues after 24h fasting challenge in control, fat-mutant mice, and mice treated as in D.
**(F)** Relative brown fat mass variation in control and fat-mutant mice after a 24-hour fast.
**(G)** Summary illustration showing fatty acid breakdown and cycling and the effect of dysregulated nutrient-deprivation on lipotoxicity (1). Cooperative transcriptional regulation of mHAR genes associated with lipid cycling and with reduced lipotoxicity. Perturbation of this pathway leads to lipotoxicity locally (seen in fat-specific transcriptional perturbations) and distantly (liver) (2).
Primary cell cultures (brown and white adipocytes) from wildtype and PGC1A-/- mice treated with 0% glucose (12 hours), and, as indicated, glucose deprivation in combination with DGAT1-inhibitor (1μM) and the Cpt1a channel inhibitor etomoxir (50μM). Cells assessed for relative mitochondrial complex I activity **(H)**, lysosome activity **(I),** cytosolic pH levels **(J)**, and MDA-lipid peroxidation assay.
Animal experiments were done with n=5-6 mice per group, both female and male mice were included. Cell experiments were done with 3 independent replicates. Bars show mean values and SEM. Unpaired, two-tailed student's t-test was used when two groups were compared, and ANOVA followed by fisher's least significant difference (LSD) test for post hoc comparisons for multiple groups. * p-value <0.05.

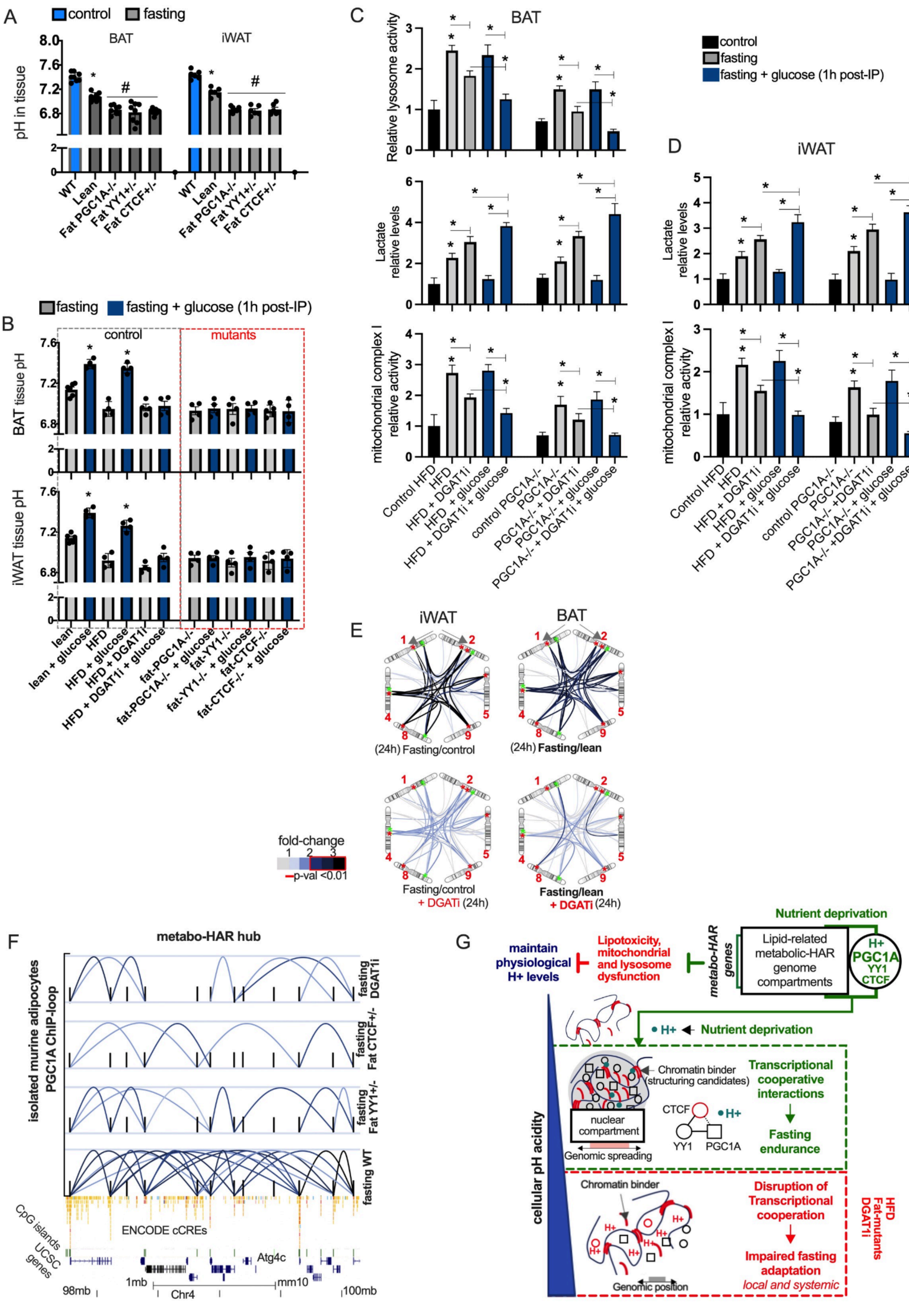

A
control
fasting
BAT
iWAT
pH in tissue
WT
Lean
Fat PGC1A-/-
Fat YY1+/-
Fat CTCF+/-
B
fasting
fasting + glucose (1h post-IP)
control
mutants
BAT tissue pH
iWAT tissue pH
lean
lean + glucose
HFD
HFD + glucose
HFD + DGAT1i
HFD + DGAT1i + glucose
fat-PGC1A-/-
fat-PGC1A-/- + glucose
fat-YY1-/-
fat-YY1-/- + glucose
fat-CTCF-/-
fat-CTCF-/- + glucose
C
BAT
Relative lysosome activity
Lactate relative levels
mitochondrial complex I relative activity
Control HFD
HFD + DGAT1i
HFD + glucose
HFD + DGAT1i + glucose
control PGC1A-/-
PGC1A-/-
PGC1A-/- + DGAT1i
PGC1A-/- + glucose
PGC1A-/- + DGAT1i + glucose
D
iWAT
E
(24h) Fasting/control
(24h) Fasting/lean
Fasting/control + DGATi (24h)
Fasting/lean + DGATi (24h)
fold-change
p-val <0.01
F
metabo-HAR hub
isolated murine adipocytes PGC1A ChIP-loop
fasting DGAT1i
fasting Fat CTCF+/-
fasting Fat YY1+/-
fasting WT
CpG islands
ENCODE cCREs
UCSC genes
Atg4c
98mb
1mb
Chr4
mm10
100mb
G
Nutrient deprivation
metabo-HAR genes
Lipid-related metabolic-HAR genome compartments
H+ PGC1A YY1 CTCF
maintain physiological H+ levels
Lipotoxicity, mitochondrial and lysosome dysfunction
H+ Nutrient deprivation
Chromatin binder (structuring candidates)
nuclear compartment
Genomic spreading
CTCF
YY1
PGC1A
Transcriptional cooperative interactions
Fasting endurance
cellular pH acidity
Chromatin binder
Genomic position
Disruption of Transcriptional cooperation
Impaired fasting adaptation local and systemic
HFD Fat-mutants DGAT1i

##### 1.11. Fig. S11. mHAR hubs modulate lipid, lysosome, and genome plasticity in fasting *(related to Fig. 3)*.

**(A)** Tissue pH levels after 24 hour fasting in control lean and HFD (1 week) mice, fat-PGC1A-/-, fat-YY1+/- and fat-CTCF+/- mice.
**(B)** Brown fat (top panel) and inguinal fat (lower panel) tissue pH levels after 24 hours fasting and corrected by glucose injections (1hour, i.p prior evaluation) in lean control, HFD (1 week), HFD + DGAT1-inhibition mice (2 mg/kg, 24h i.p), fat-mutant mice, and fat-mutant mice treated with DGAT1-inhibition.
Lactate relative levels, lysosome activity, and mitochondrial complex I activity in brown (**C**) and inguinal fat (**D**) from mice as in **B**.
**(E)** Targeted in situ chromosome conformation assays (3C) for metabolic-HAR regions within associated nuclear compartment in brown and white fat from control mice after fasting (24h) and fasting with DGAT1-inhibition (2 mg/kg, 12h i.p). Contact variations are represented as fold-change over control groups (not fasted littermates). Contacts with FC > 2 have a p-value <0.01.
**(F)** PGC1A ChIP-loop of regulatory region interactions in mHAR genomic hub from freshly isolated brown adipocytes from control and mutant mice undergoing fasting (24h) with or without DGAT1-inhibition (2 mg/kg). Contact variations are represented as fold-change over control groups (not fasted littermates). Contacts with FC > 2 have a p-value <0.01.
**(G)** Summary illustration showing cooperative transcriptional regulation of mHAR associated nuclear compartments harboring genomic hubs controlling genome plasticity through lipid and pH homeostasis.
Animal experiments were done with n=5-6 mice per group, both female and male mice were included. Bars show mean values and SEM. Unpaired, two-tailed student's t-test was used when two groups were compared, and ANOVA followed by fisher's least significant difference (LSD) test for post hoc comparisons for multiple groups. * p-value <0.05.

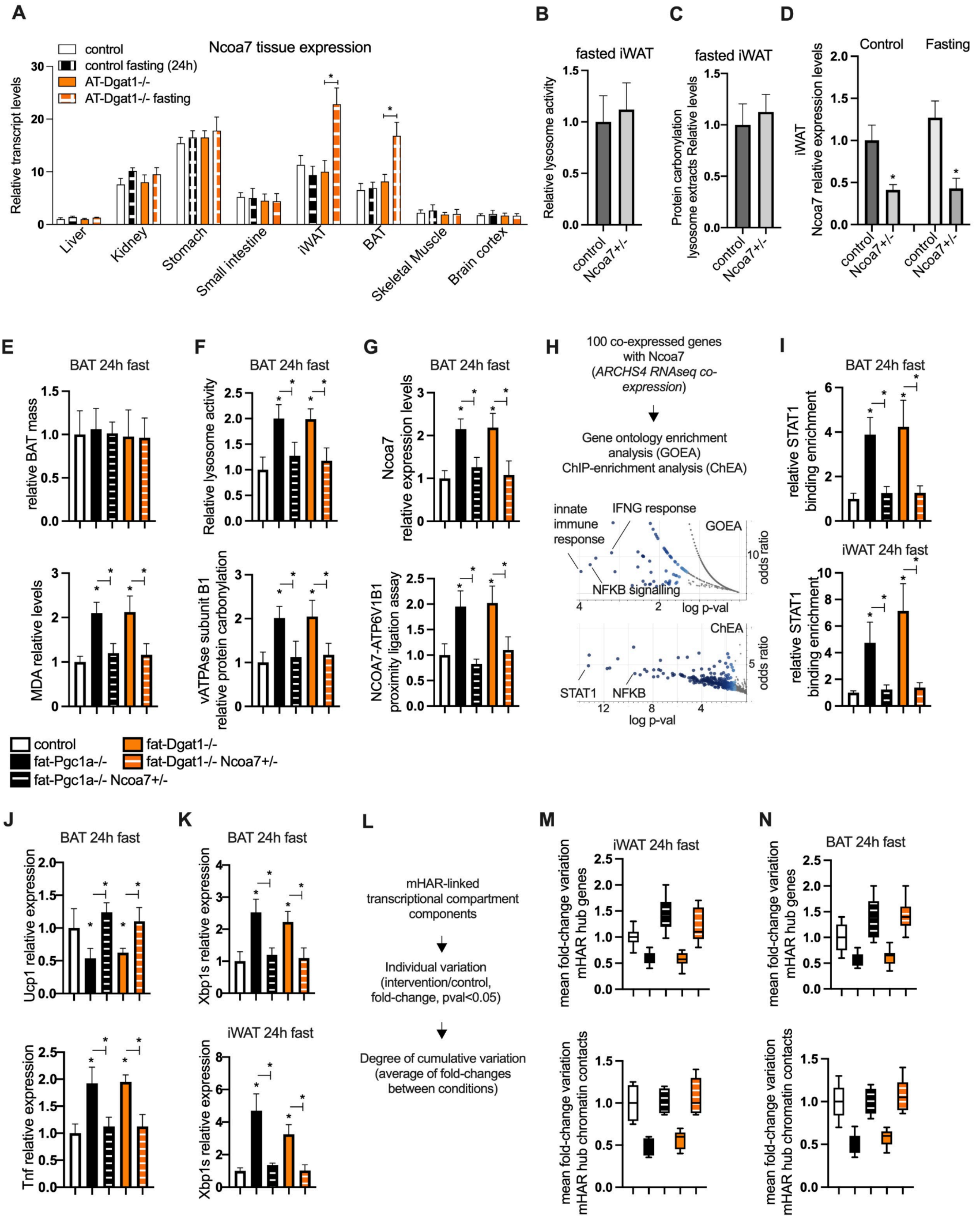
A
Ncoa7 tissue expression
control
control fasting (24h)
AT-Dgat1-/-
AT-Dgat1-/- fasting
Relative transcript levels
Liver
Kidney
Stomach
Small intestine
iWAT
BAT
Skeletal Muscle
Brain cortex
B
fasted iWAT
Relative lysosome activity
control
Ncoa7+/-
C
fasted iWAT
Protein carbonylation lysosome extracts Relative levels
D
Control
Fasting
iWAT Ncoa7 relative expression levels
E
BAT 24h fast
relative BAT mass
MDA relative levels
F
BAT 24h fast
Relative lysosome activity
vATPase subunit B1 relative protein carbonylation
G
BAT 24h fast
Ncoa7 relative expression levels
NCOA7-ATP6V1B1 proximity ligation assay
H
100 co-expressed genes with Ncoa7 (ARCHS4 RNAseq co-expression)
Gene ontology enrichment analysis (GOEA)
ChIP-enrichment analysis (ChEA)
innate immune response
IFNG response
NFKB signalling
GOEA
odds ratio
log p-val
ChEA
STAT1
NFKB
I
BAT 24h fast
relative STAT1 binding enrichment
iWAT 24h fast
control
fat-Pgc1a-/-
fat-Pgc1a-/- Ncoa7+/-
fat-Dgat1-/-
fat-Dgat1-/- Ncoa7+/-
J
BAT 24h fast
Ucp1 relative expression
Tnf relative expression
K
BAT 24h fast
Xbp1s relative expression
iWAT 24h fast
L
mHAR-linked transcriptional compartment components
Individual variation (intervention/control, fold-change, pval<0.05)
Degree of cumulative variation (average of fold-changes between conditions)
M
iWAT 24h fast
mean fold-change variation mHAR hub genes
mean fold-change variation mHAR hub chromatin contacts
N
BAT 24h fast

##### 1.12. Fig. S12. mHAR hubs modulate NCOA7 for fasting endurance *(related to Fig. 3)*.

**(A)** Ncoa7 transcript levels in different murine tissue in control and fat-Dgat1-/- mice in unchallenged or after 24 hours of fasting. Relative expression normalized by expression in the liver
**(B)** In unchallenged control and Ncoa7+/- heterozygous mice, inguinal fat relative lysosomal activity
**(C)** Protein carbonylation assays as in D.
**(D)** Ncoa7 gene expression levels in unchallenged and fasted (24 h) mice.
**(E)** In fasted (24h) fat-specific mice with single or double genetic perturbations, upper panel shows brown fat mass and, in lower panel, MDA lipid peroxidation levels ,
**(F)** Upper panel, brown fat lysosome activity. Lower panel, vATPase-protein carbonylation in mice as in E.
**(G)** Upper panel, brown fat Ncoa7 expression. Lower panel, NCOA7-vATPases interaction by PLA assays in mice as in E.
**(H)** Pipeline description followed by gene ontology and ChIP-binding enrichment analysis (GOEA, ChEA) using Enrichr analysis tool for Ncoa7 co-expressed genes from Enrichr repository.
**(I)** STAT1 binding enrichment to Ncoa7 promoter by ChIP-qPCR in mice described in E.
**(J)** Ucp1 (top panel) and Tnf (lower panel) gene expression.
**(K)** ER-stress gene marker expression in brown and inguinal fat as in E.
**(L)** Pipeline description for evaluating mHAR associated transcriptional compartment components.
Top panel, mean fold-change variation in expression of mHAR genomic hub genes (within associated compartment, n=12-16 genes) in **(M)** white fat and **(N)** brown fat from fasted (24h) fat-specific mice with single or double genetic perturbations, assessed as in L. Lower panel shows chromatin contacts (targeted chromosome conformation assays) between mHAR genomic hubs shown as mean fold change variation in at least 10 contacts per group.
Animal experiments were done with n=4-6 mice per group, both female and male mice were included. Bars show mean values and SEM. Unpaired, two-tailed student's t-test was used when two groups were compared, and ANOVA followed by fisher's least significant difference (LSD) test for post hoc comparisons for multiple groups. * $p$-value $<0.05$.

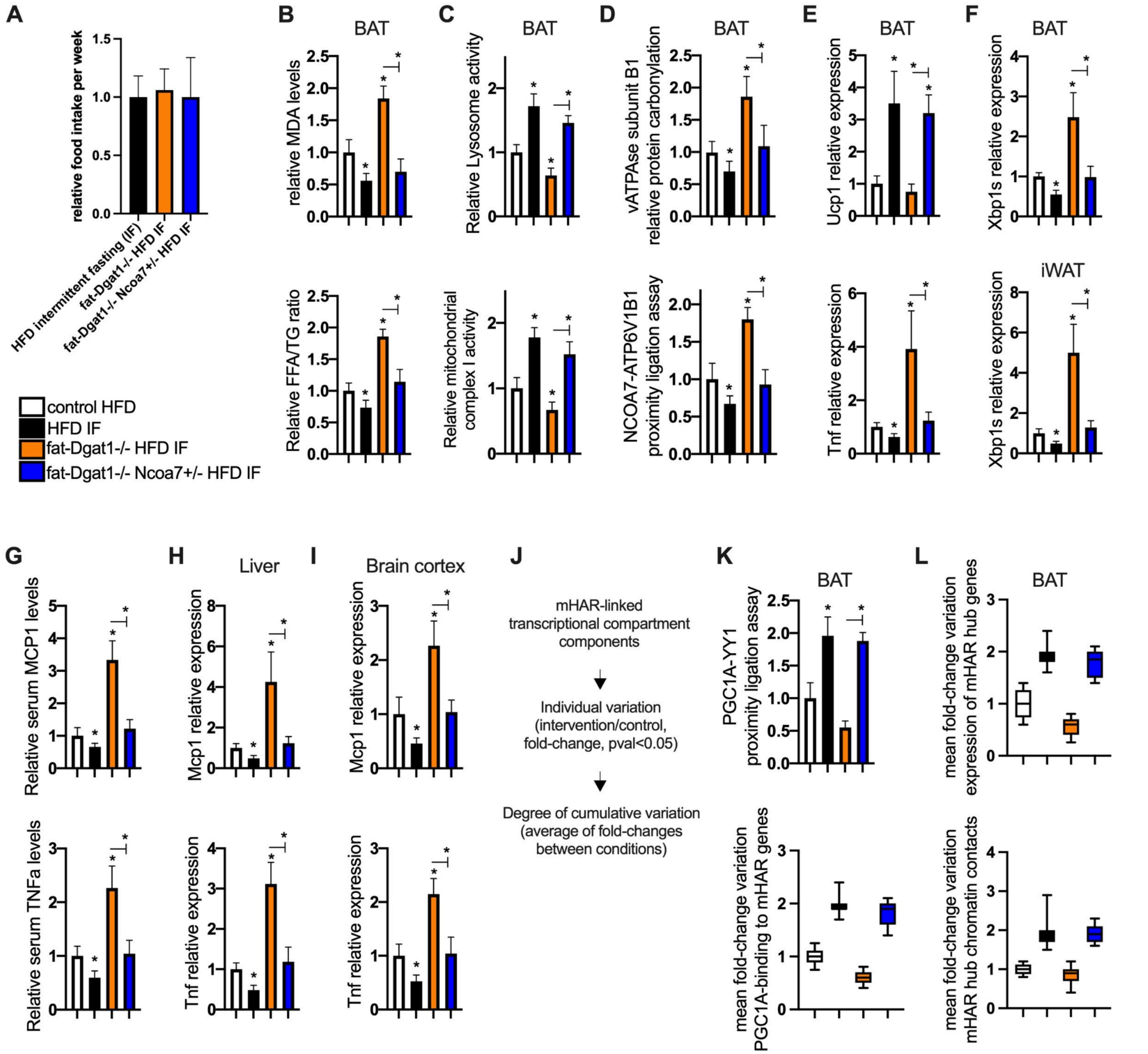

**1.13. Fig. S13. NCOA7 modulation enhances genome plasticity and metabolic homeostasis in intermittent fasting** *(related to Fig. 4)***.**

**(A)** Food intake relative variation per week in control HFD mice and mice (control and mutant mice) on a HFD and treated with 2-day bouts of intermittent fasting (IF) every week.
**(B)** Brown fat MDA lipid peroxidation levels (top panel) and free-fatty-acids/TG ratio (lower panel) from mice as in A.
**(C)** Brown fat lysosome (top panel) and mitochondrial activity levels (lower panel) from mice as in A.
**(D)** Brown fat vATPase subunit protein carbonylation (top panel) and NCOA7-vATPase subunit relative interaction levels (lower panel) from mice as in A.
**(E)** ER-stress gene marker expression in brown (top panel) and inguinal (lower panel) fat as in A.
**(F)** Ucp1 (top panel) and Tnf (lower panel) gene expression.
**(G)** Relative serum MCP1 (top panel) and TNFA (lower panel) levels in mice as in A.
**(H)** Liver Mcp1 (top panel) and Tnf (lower panel) gene expression
**(I)** Brain cortex Mcp1 (top panel) and Tnf (lower panel) gene expression
**(J)** Pipeline description for evaluating mHAR associated nuclear compartment components.

**(K)** Proximity ligation assays for protein interactions (top panel) and PGC1A binding by ChIP-qPCR to mHAR genomic hub genes within the associated nuclear compartment (lower panel). Data in the lower panel is shown as mean fold-change variations in binding to at least 10 mHAR genomic hub promoters.
**(L)** Top panel, mean fold-change variation in expression of mHAR genomic hub genes (within associated compartment, n=12-16 genes) in inguinal fat from mice as in A. Lower panel shows chromatin contacts (targeted chromosome conformation assays) between mHAR genomic hubs shown as mean fold change variation in at least 10 contacts per group. Animal experiments were done with n=5-6 mice per group, both female and male mice were included. Data show mean values and SEM. Unpaired, two-tailed student's t-test was used when two groups were compared, and ANOVA followed by fisher's least significant difference (LSD) test for post hoc comparisons for multiple groups. * p-value <0.05.

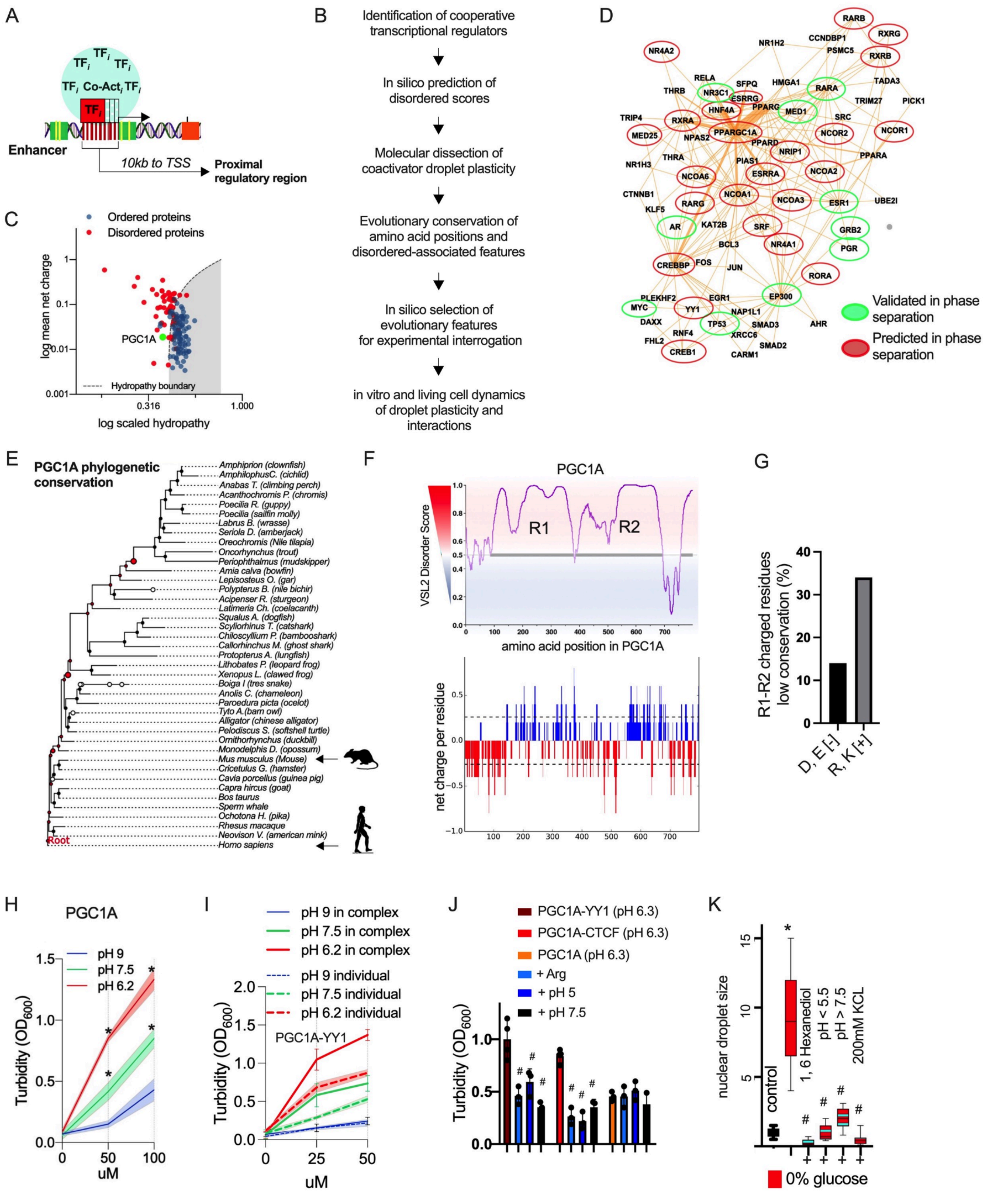
A
TF$_i$ TF$_i$ TF$_i$
TF$_i$ Co-Act$_i$ TF$_i$
TF$_i$
Enhancer
10kb to TSS
Proximal regulatory region
B
Identification of cooperative transcriptional regulators
In silico prediction of disordered scores
Molecular dissection of coactivator droplet plasticity
Evolutionary conservation of amino acid positions and disordered-associated features
In silico selection of evolutionary features for experimental interrogation
in vitro and living cell dynamics of droplet plasticity and interactions
C
Ordered proteins
Disordered proteins
log mean net charge
PGC1A
Hydropathy boundary
log scaled hydropathy
D
Validated in phase separation
Predicted in phase separation
E
PGC1A phylogenetic conservation
Amphiprion (clownfish)
AmphilophusC. (cichlid)
Anabas T. (climbing perch)
Acanthochromis P. (chromis)
Poecilia R. (guppy)
Poecilia (sailfin molly)
Labrus B. (wrasse)
Seriola D. (amberjack)
Oreochromis (Nile tilapia)
Oncorhynchus (trout)
Periophthalmus (mudskipper)
Amia calva (bowfin)
Lepisosteus O. (gar)
Polypterus B. (nile bichir)
Acipenser R. (sturgeon)
Latimeria Ch. (coelacanth)
Squalus A. (dogfish)
Scyliorhinus T. (catshark)
Chiloscyllium P. (bambooshark)
Callorhinchus M. (ghost shark)
Protopterus A. (lungfish)
Lithobates P. (leopard frog)
Xenopus L. (clawed frog)
Boiga I (tres snake)
Anolis C. (chameleon)
Paroedura picta (ocelot)
Tyto A.(barn owl)
Alligator (chinese alligator)
Pelodiscus S. (softshell turtle)
Ornithorhynchus (duckbill)
Monodelphis D. (opossum)
Mus musculus (Mouse)
Cricetulus G. (hamster)
Cavia porcellus (guinea pig)
Capra hircus (goat)
Bos taurus
Sperm whale
Ochotona H. (pika)
Rhesus macaque
Neovison V. (american mink)
Homo sapiens
Root
F
PGC1A
VSL2 Disorder Score
R1
R2
amino acid position in PGC1A
net charge per residue
G
R1-R2 charged residues low conservation (%)
D, E [-]
R, K [+]
H
PGC1A
pH 9
pH 7.5
pH 6.2
Turbidity (OD$_{600}$)
uM
I
pH 9 in complex
pH 7.5 in complex
pH 6.2 in complex
pH 9 individual
pH 7.5 individual
pH 6.2 individual
PGC1A-YY1
Turbidity (OD$_{600}$)
uM
J
PGC1A-YY1 (pH 6.3)
PGC1A-CTCF (pH 6.3)
PGC1A (pH 6.3)
+ Arg
+ pH 5
+ pH 7.5
Turbidity (OD$_{600}$)
K
nuclear droplet size
control
1, 6 Hexanediol
pH < 5.5
pH > 7.5
200mM KCL
0% glucose

##### 1.14. Fig. S14. PGC1A conformational features, evolution, and droplet turbidity *(related to Fig. 5)*.

**(A)** Schematic illustration of cooperative transcriptional regulation through condensates.
**(B)** Pipeline representation for the evaluation of droplet plasticity associated with evolutionary adaptations.
**(C)** Net charge and scaled hydropathy of co-activator PGC1A, along with ordered and disordered regions from PONDR software.
**(D)** Brain and adipose tissue backbone PPI network, displaying interacting PGC1A regulatory network. In green, regulators forming condensates, both predicted in silico and validated experimentally, and, in red, those only predicted in silico.
**(E)** Phylogenetic tree conservation of PGC1A amino acid sequence using CONSURF.
**(F)** VSL2 disorder score for the transcriptional coactivator PGC1A amino acid sequence. Lower panel, net charge per residue for PGC1A sequence.
**(G)** Degree of conservation for positively and negatively charged residues in disordered regions of PGC1A.
**(H)** Turbidity assays of recombinant PGC1A protein with pH variation.
**(I)** Turbidity assays from individual PGC1A and in complex with YY1 recombinant proteins with pH variation.
**(J)** Turbidity assay from individual and in complex recombinant proteins and with arginine and pH variations.
**(K)** Relative nuclear droplet size for PGC1A in HEK293 cells under different conditions (fold-change over control. Data show mean values and SEM. Unpaired, two-tailed student's t-test was used when two groups were compared, and ANOVA followed by fisher's least significant difference (LSD) test for post hoc comparisons for multiple groups. Two-way ANOVA was used to estimate significance between groups constrained by time and concentration variations, followed by Tukey test for post-hoc multiple comparisons. Cell experiments were done with 3 independent replicates. *, # p-value $<0.05$. # p-value $<0.05$ when comparing to 0% glucose treatment (panel M).

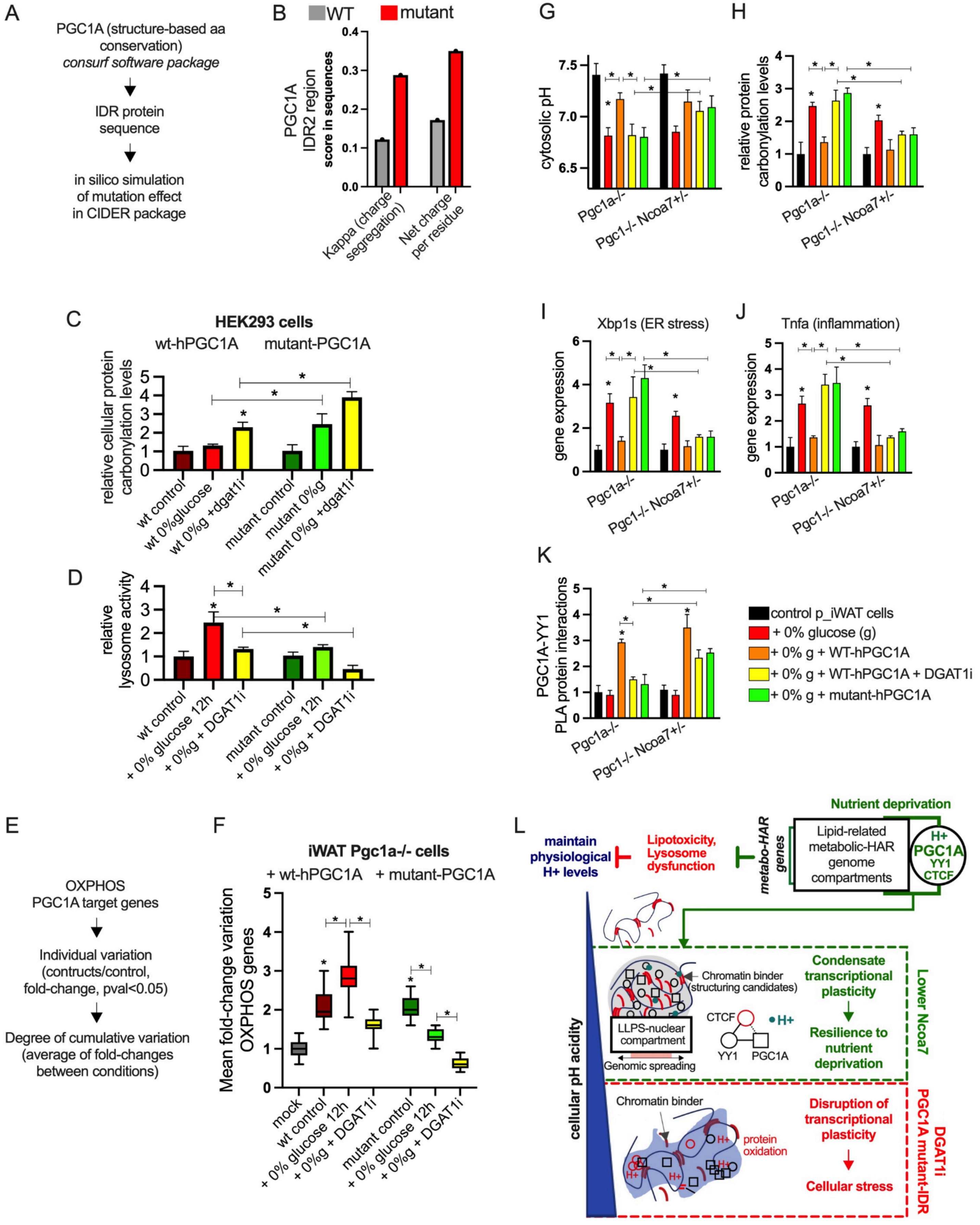
A
PGC1A (structure-based aa conservation)
consurf software package
IDR protein sequence
in silico simulation of mutation effect in CIDER package
B
WT
mutant
PGC1A IDR2 region score in sequences
Kappa (charge segregation)
Net charge per residue
C
HEK293 cells
wt-hPGC1A
mutant-PGC1A
relative cellular protein carbonylation levels
wt control
wt 0%glucose
wt 0%g +dgat1i
mutant control
mutant 0%g
mutant 0%g +dgat1i
D
relative lysosome activity
wt control
+ 0% glucose 12h
+ 0%g + DGAT1i
mutant control
+ 0% glucose 12h
+ 0%g + DGAT1i
E
OXPHOS PGC1A target genes
Individual variation (contructs/control, fold-change, pval<0.05)
Degree of cumulative variation (average of fold-changes between conditions)
F
iWAT Pgc1a-/- cells
+ wt-hPGC1A
+ mutant-PGC1A
Mean fold-change variation OXPHOS genes
mock
wt control
+ 0% glucose 12h
+ 0%g + DGAT1i
mutant control
+ 0% glucose 12h
+ 0%g + DGAT1i
G
cytosolic pH
Pgc1a-/-
Pgc1-/- Ncoa7+/-
H
relative protein carbonylation levels
Pgc1a-/-
Pgc1-/- Ncoa7+/-
I
Xbp1s (ER stress)
gene expression
Pgc1a-/-
Pgc1-/- Ncoa7+/-
J
Tnfa (inflammation)
gene expression
Pgc1a-/-
Pgc1-/- Ncoa7+/-
K
PGC1A-YY1 PLA protein interactions
Pgc1a-/-
Pgc1-/- Ncoa7+/-
control p_iWAT cells
+ 0% glucose (g)
+ 0% g + WT-hPGC1A
+ 0% g + WT-hPGC1A + DGAT1i
+ 0% g + mutant-hPGC1A
L
maintain physiological H+ levels
Lipotoxicity, Lysosome dysfunction
metabo-HAR genes
Lipid-related metabolic-HAR genome compartments
Nutrient deprivation
H+
PGC1A
YY1
CTCF
Chromatin binder (structuring candidates)
LLPS-nuclear compartment
Genomic spreading
CTCF
H+
YY1
PGC1A
Condensate transcriptional plasticity
Resilience to nutrient deprivation
Lower Ncoa7
cellular pH acidity
Chromatin binder
protein oxidation
Disruption of transcriptional plasticity
Cellular stress
DGAT1i
PGC1A mutant-IDR

#### 1.15. Fig. S15. PGC1A droplet plasticity influences mHAR hub activity in nutrient stress *(related to Fig. 5)*.

**(A)** Pipeline summary description to identify IDR residues impacting charge related features and with different conservation scales in IDR of PGC1A using the CONSURF and CIDER packages.
**(B)** In silico simulation of conformational features in CIDER from wild-type PGC1A IDR2 sequence and PGC1A IDR2 sequence with mutant residues (as shown in A). Graph shows disorder parameters for charge segregation and net charge.
**(C)** Protein carbonyl content assay in HEK293 cells transfected with wildtype and mutant PGC1A, and under different conditions. 0% glucose (12h), transcription factor knockdown by RNAi, and DGAT1-inhibition (12h; 1µM).
**(D)** Relative lysosome activity in HEK293 as in C.
**(E)** Pipeline description for evaluating mitochondrial oxidative phosphorylation genes regulated by PGC1A.
**(F)** Mean fold-change variation in gene expression of OXPHOs genes ( n=18-20 genes) in HEK293 treated as in C.
**(G)** Cytosolic pH levels in inguinal primary adipocytes control (from fat-specific Pgc1a-/- mice or mice with double perturbations such as fat-Pgc1a-/ and Ncoa7+/-) and under different conditions including glucose deprivation (12h) either alone or with nucleoporation (using the nucleofector instrument and protocols for primary cells) of PGC1A vectors alone or together with DGAT1-inhibition (12h, 1µM).
**(H)** Relative protein carbonylation levels in cellular protein extracts from primary cells treated as in G.
**(I)** ER-stress gene marker expression from primary cells as in G.
**(J)** Tnf gene marker expression from primary cells as in G.
**(K)** Relative protein interactions by proximity ligation assays in cell extracts from primary cells treated as in G.
**(L)** Summary illustration for transcriptional droplet plasticity during nutrient stress.
Cell experiments were done with 3 independent replicates. Data show mean values and SEM. Unpaired, two-tailed student's t-test was used when two groups were compared, and ANOVA followed by fisher's least significant difference (LSD) test for post hoc comparisons for multiple groups. * p-value <0.05 when indicated and when compared to the control group.

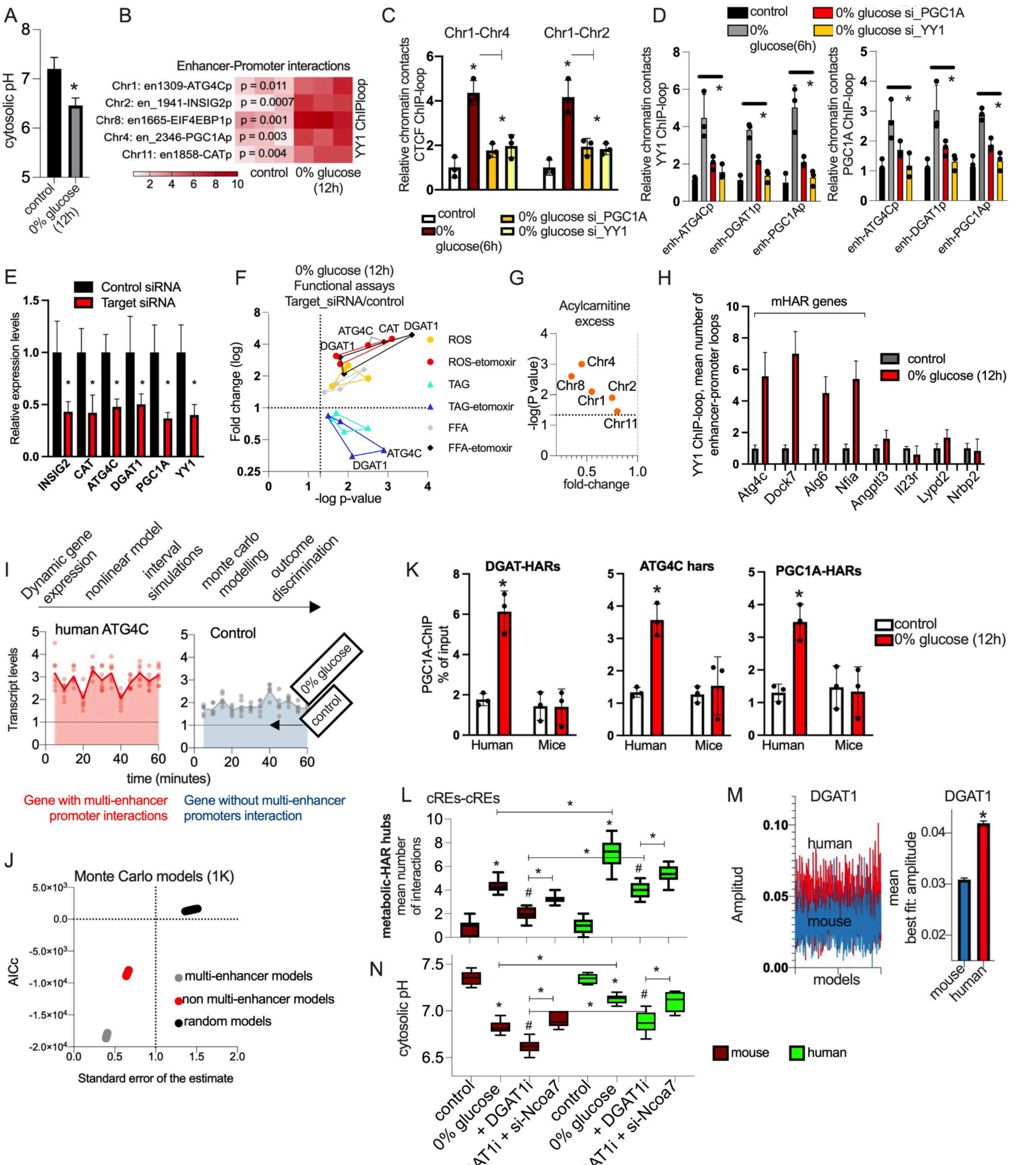
A
cytosolic pH
control
0% glucose (12h)
B
Enhancer-Promoter interactions
Chr1: en1309-ATG4Cp p = 0.011
Chr2: en_1941-INSIG2p p = 0.0007
Chr8: en1665-EIF4EBP1p p = 0.001
Chr4: en_2346-PGC1Ap p = 0.003
Chr11: en1858-CATp p = 0.004
YY1 ChIPloop
control 0% glucose (12h)
C
Chr1-Chr4
Chr1-Chr2
Relative chromatin contacts CTCF ChIP-loop
control
0% glucose(6h)
0% glucose si_PGC1A
0% glucose si_YY1
D
control
0% glucose(6h)
0% glucose si_PGC1A
0% glucose si_YY1
Relative chromatin contacts YY1 ChIP-loop
Relative chromatin contacts PGC1A ChIP-loop
enh-ATG4Cp
enh-DGAT1p
enh-PGC1Ap
E
Control siRNA
Target siRNA
Relative expression levels
INSIG2
CAT
ATG4C
DGAT1
PGC1A
YY1
F
0% glucose (12h)
Functional assays
Target_siRNA/control
Fold change (log)
-log p-value
ROS
ROS-etomoxir
TAG
TAG-etomoxir
FFA
FFA-etomoxir
G
Acylcarnitine excess
-log(P value)
fold-change
Chr4
Chr8
Chr1
Chr2
Chr11
H
mHAR genes
YY1 ChIP-loop, mean number of enhancer-promoter loops
control
0% glucose (12h)
Atg4c
Dock7
Alg6
Nfia
Angptl3
Il23r
Lypd2
Nrbp2
I
Dynamic gene expression
nonlinear model
interval simulations
monte carlo modelling
outcome discrimination
human ATG4C
Control
Transcript levels
time (minutes)
0% glucose
control
Gene with multi-enhancer promoter interactions
Gene without multi-enhancer promoters interaction
J
Monte Carlo models (1K)
AICc
Standard error of the estimate
multi-enhancer models
non multi-enhancer models
random models
K
DGAT-HARs
ATG4C hars
PGC1A-HARs
PGC1A-ChIP % of input
Human
Mice
control
0% glucose (12h)
L
cREs-cREs
metabolic-HAR hubs mean number of interactions
M
DGAT1
Amplitud
human
mouse
models
mean best fit: amplitude
N
cytosolic pH
mouse
human
control
0% glucose
+ DGAT1i
+ DGAT1i + si-Ncoa7

##### 1.16. Fig. S16. Species-specific mHAR activity in nutrient stress. *(related to Fig. 6)*.

**(A)** Cytosolic pH variation after 12 h 0% glucose in human primary adipocytes.
**(B)** YY1-mediated enhancer-promoter loops between HAR-elements and promoters of mHAR genes in human primary adipocytes after 12 h 0% glucose.
**(C)** CTCF-mediated long-range chromatin interactions between mHAR regions by ChIP-loop after 6 hours of glucose starvation and with targeted siRNA.
**(D)** Relative short-range interactions by ChIP-loop as in **C** between top regulated metabolic-HAR gene promoters and HAR-elements.
**(E)** Gene expression after siRNA knockdowns in human primary adipocytes.
**(F)** RNAi perturbation screen for top regulated mHAR genes within lipid-related compartment. Functional assays in human primary adipocytes under 0% glucose for 12h, and with or without CPT1A-channel inhibitor etomoxir (50µM).
**(G)** PGC1A HAR-enhancer binding by ChIP-qPCR in human adipocytes after glucose starvation (12h) with acylcarnitine (AC) excess (20µM).
**(H)** YY1 ChIP-loop, mean number of local enhancer interactions for mHAR promoters and neighboring genes within the mHAR-genomic hub in human primary adipocytes after glucose deprivation (12h). Positive regulatory region interactions have a fold-change over control of (FC) >2 and a p-value <0.01.
**(I)** Top panel, pipeline description for Monte Carlo (MC) modeling of dynamic gene expression. Panels below are dynamic transcript levels (fold-change variation every 10 min) in human adipocytes under 6 hour of glucose starvation: ATG4C (promoter with large number of enhancer-promoter associations) and ANGPTL3 (promoter, without enhancer-promoter association, "Control burst") fold-change variation.
**(J)** Akaike information criterion and standard error of the estimate for MC models from dynamic gene expression comparing genes and random models.
**(K)** PGC1A-binding by ChIP-qPCR (human and mice primary adipocytes, 12h 0% glucose) in selected HAR elements and in mouse orthologue positions for mHAR regions ATG4C-, DGAT1-, and PGC1A-hubs.
**(L)** Primary white adipocytes after glucose starvation (12h) with or without DGAT1-inhibition (1uM), or with DGAT1-inhibition combined with siRNA for Ncoa7 (scramble control in other conditions). Mean number of enhancer interactions by PGC1A ChIP-loop within metabolic-HAR hubs in associated nuclear compartment.
**(M)** Amplitude variation of MC models from dynamic gene expression data for murine and human DGAT1 expression. Right panel, mean amplitude of MC models for DGAT1.
**(N)** Cytosolic pH levels in cells as in L.
Cell experiments were done with 3 independent replicates. Data show mean values and SEM. Unpaired, two-tailed student's t-test was used when two groups were compared, and ANOVA followed by fisher's least significant difference (LSD) test for post hoc comparisons for multiple groups. * p-value <0.05 when indicated and when compared to the control group. # p-value <0.05 when compared to 0% glucose control in panels L and N.

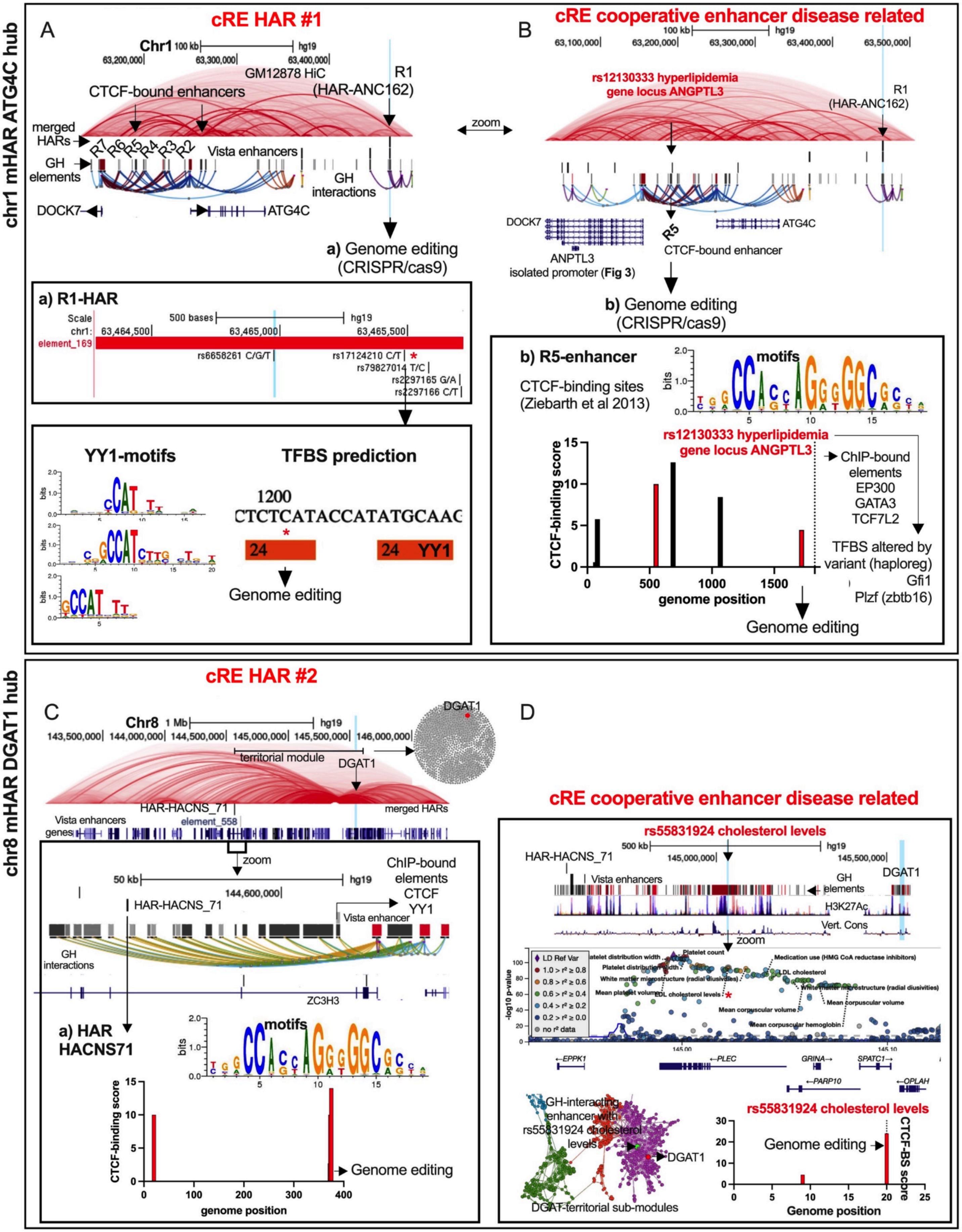

chr1 mHAR ATG4C hub
A
cRE HAR #1
Chr1
100 kb
hg19
63,200,000
63,300,000
63,400,000
GM12878 HiC
R1
(HAR-ANC162)
CTCF-bound enhancers
merged HARs
R7 R6 R5 R4 R3 R2
Vista enhancers
GH elements
GH interactions
DOCK7
ATG4C
a) Genome editing (CRISPR/cas9)
zoom
B
cRE cooperative enhancer disease related
63,100,000
63,500,000
rs12130333 hyperlipidemia gene locus ANGPTL3
R1 (HAR-ANC162)
R5
CTCF-bound enhancer
ANPTL3 isolated promoter (Fig 3)
b) Genome editing (CRISPR/cas9)
a) R1-HAR
Scale
500 bases
chr1:
63,464,500
63,465,000
63,465,500
element_169
rs6658261 C/G/T
rs17124210 C/T
rs79827014 T/C
rs2297165 G/A
rs2297166 C/T
YY1-motifs
TFBS prediction
1200
CTCTCATACCATATGCAAC
24
24 YY1
Genome editing
b) R5-enhancer
motifs
CTCF-binding sites (Ziebarth et al 2013)
rs12130333 hyperlipidemia gene locus ANGPTL3
CTCF-binding score
genome position
ChIP-bound elements
EP300
GATA3
TCF7L2
TFBS altered by variant (haploreg)
Gfi1
Plzf (zbtb16)
Genome editing
chr8 mHAR DGAT1 hub
C
cRE HAR #2
Chr8
1 Mb
143,500,000
144,000,000
144,500,000
145,000,000
145,500,000
146,000,000
DGAT1
territorial module
HAR-HACNS_71
merged HARs
Vista enhancers
element_558
genes
zoom
50 kb
144,600,000
ChIP-bound elements
CTCF
YY1
Vista enhancer
GH interactions
ZC3H3
a) HAR HACNS71
motifs
CTCF-binding score
genome position
Genome editing
D
cRE cooperative enhancer disease related
rs55831924 cholesterol levels
500 kb
145,000,000
145,500,000
HAR-HACNS_71
Vista enhancers
GH elements
H3K27Ac
Vert. Cons
zoom
LD Ref Var
no r² data
-log10 p-value
Platelet distribution width
Platelet count
Medication use (HMG CoA reductase inhibitors)
LDL cholesterol
White matter microstructure (radial diusivities)
Mean platelet volume
LDL cholesterol levels
Mean corpuscular volume
Mean corpuscular hemoglobin
EPPK1
PLEC
GRINA
SPATC1
PARP10
OPLAH
GH-interacting enhancer with rs55831924 cholesterol levels
DGAT1
DGAT-territorial sub-modules
rs55831924 cholesterol levels
Genome editing
CTCF-BS score
Genome position

#### 1.17. Fig. S17. Dissection of mHAR disease circuit with genome editing *(related to Fig. 7)*.

**(A)** Human genome track showing top targets within metabolic-HAR domain in chr1:62-63mb: ncHARs (ANC162), regulatory elements and interactions, GM12878-Hi-C data, CTCF-bound elements, and promoter locations for ATG4C and DOCK7. Lower panel shows variants within the HAR-ANC162 enhancer (element_169). With an arrow, location of transcription factor binding site (TFBS) for YY1 transcription factor, selected for genome editing.
**(B)** Human genome track as in A, showing with arrows selected enhancers, in which the R5 element (CTCF-bound) harbors a Single Nucleotide Polymorphism (SNP) related to hyperlipidemia (rs12130333). Lower panel shows R5 element scanned for CTCF-binding sites as described by Ziebarth et al(*9*). A CTCF-binding site in proximity to the variant rs12130333 was selected for genome editing.
**(C)** Human genome track showing top targets within mHAR domain in chr8:144-146mb: ncHARs (HACNS_71), vista regulatory element 558, GM12878-Hi-C data, DGAT1-promoter location, and DGAT1-module reconstruction from regulatory elements interaction network. Lower panel shows HAR-HACNS_71, vista regulatory element 558, regulatory elements and interactions by GeneHancer. With an arrow to the lowest panel CTCF-binding sites in the HAR element, selected for genome editing.
**(D)** Human genome track showing with an arrow selected variant related to cholesterol levels in regulatory element, followed by the middle panel showing all SNPs within the region (by HugAmp metabolic database) and with an asterisk the lipid-related variant rs55831924. Lowest panel shows force atlas layout of DGAT1 module and submodules from genome-wide regulatory region associations (GeneHancer data). Highlighted in green the lipid-variant and in red DGAT1 promoter element. Right panel, CTCF-binding sites for the element harboring the lipid-variant selected for genome editing. Cell experiments were done with 3 independent replicates. Graphs show mean values and SEM. Unpaired, two-tailed student's t-test was used when two groups were compared, and ANOVA followed by fisher's least significant difference (LSD) test for post hoc comparisons for multiple groups. * p-value <0.05.

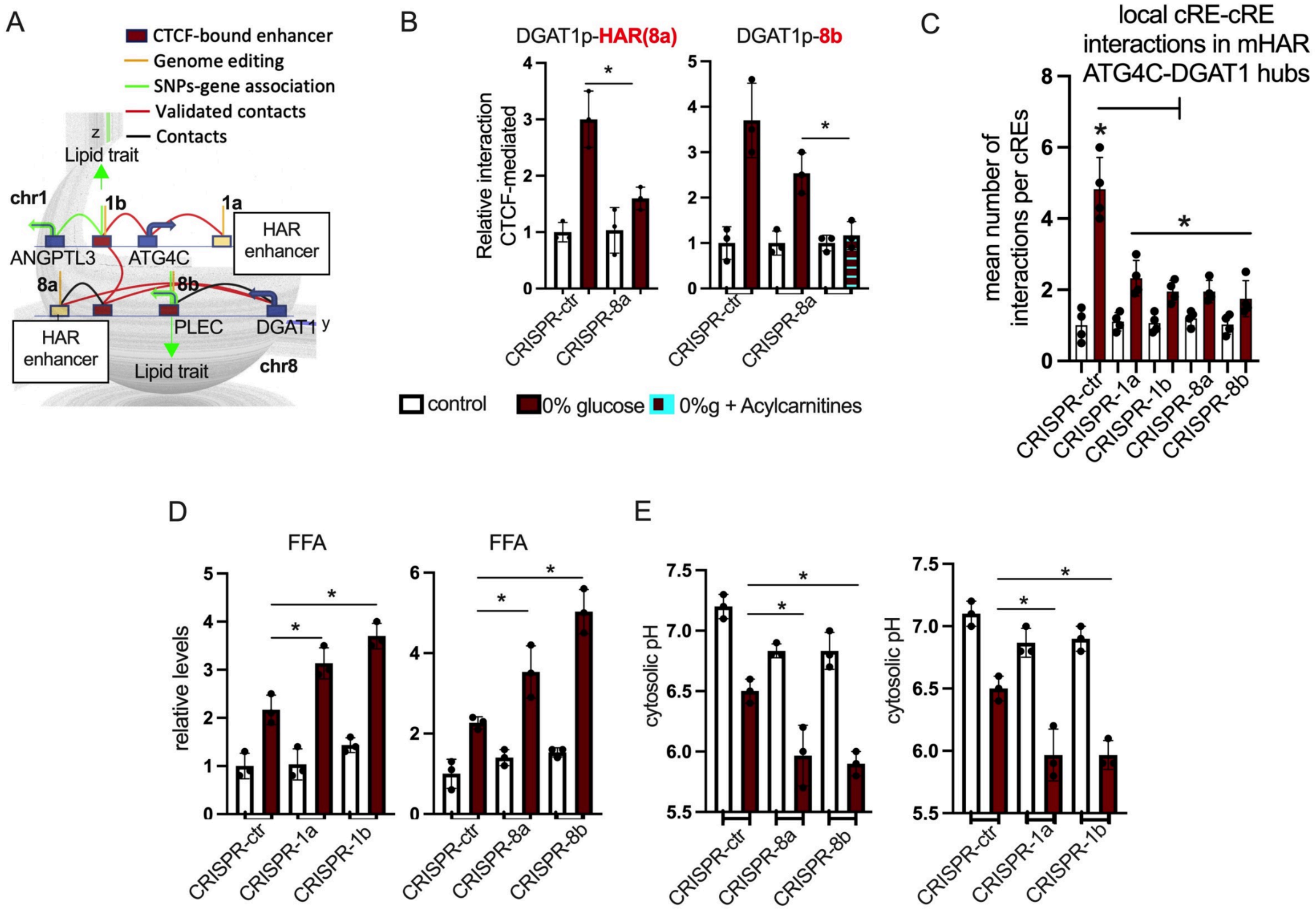

**1.18. Fig. S18. Functional impact of edited mHAR disease circuit** *(related to Fig. 7)***.**

**(A)** Network reconstruction of Chr1-Chr8 corregulated mHAR domains. In green arrow, lipid-related variants within structural enhancers (defined as CTCF-bound), HAR-locations, target genes, chromatin contacts as well as notation for genome editing interventions.
**(B)** Left panel shows relative CTCF-mediated loops (ChIP-loop) between HAR-element and DGAT1_promoter in control mutation and edited HAR CTCF-binding site in human adipocytes after glucose starvation. Right panel shows enhancer 8b-DGAT1_promoter interaction in the same conditions and with acylcarnitine (10uM).
**(C)** Mean number of interactions between regulatory elements by PGC1A ChIP-loop within mHAR hubs comparing genome editing perturbations as described in A**.**
**(D)** Free-fatty acids relative levels in the same conditions, comparing control mutations, HAR mutations (a) and mutation in key-structural enhancers (b).
**(E)** Cytosolic pH in the same conditions.
Cell experiments were done with 3 independent replicates. Graphs show mean values and SEM. Unpaired, two-tailed student's t-test was used when two groups were compared, and ANOVA followed by fisher's least significant difference (LSD) test for post hoc comparisons for multiple groups. * p-value <0.05.

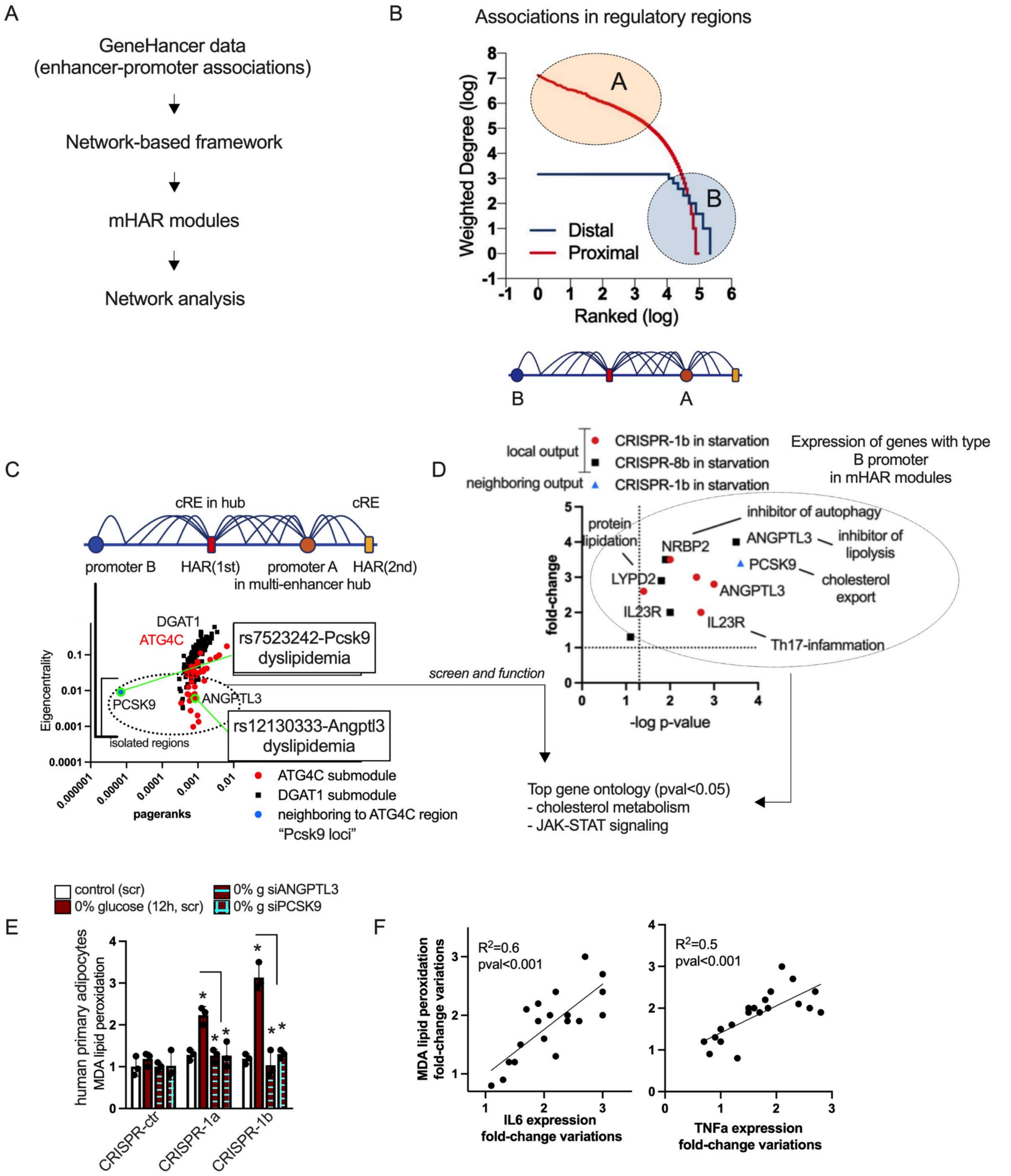
A
GeneHancer data
(enhancer-promoter associations)
Network-based framework
mHAR modules
Network analysis
B
Associations in regulatory regions
Weighted Degree (log)
Ranked (log)
Distal
Proximal
C
cRE in hub
cRE
promoter B
HAR(1st)
promoter A
in multi-enhancer hub
HAR(2nd)
DGAT1
ATG4C
rs7523242-Pcsk9
dyslipidemia
PCSK9
ANGPTL3
rs12130333-Angptl3
dyslipidemia
isolated regions
Eigencentrality
pageranks
ATG4C submodule
DGAT1 submodule
neighboring to ATG4C region
"Pcsk9 loci"
screen and function
D
local output
neighboring output
CRISPR-1b in starvation
CRISPR-8b in starvation
CRISPR-1b in starvation
Expression of genes with type
B promoter
in mHAR modules
protein
lipidation
inhibitor of autophagy
NRBP2
ANGPTL3
inhibitor of
lipolysis
PCSK9
cholesterol
export
LYPD2
ANGPTL3
IL23R
IL23R
Th17-infammation
fold-change
-log p-value
Top gene ontology (pval<0.05)
- cholesterol metabolism
- JAK-STAT signaling
E
control (scr)
0% glucose (12h, scr)
0% g siANGPTL3
0% g siPCSK9
human primary adipocytes
MDA lipid peroxidation
CRISPR-ctr
CRISPR-1a
CRISPR-1b
F
R2=0.6
pval<0.001
MDA lipid peroxidation
fold-change variations
IL6 expression
fold-change variations
R2=0.5
pval<0.001
TNFa expression
fold-change variations

**1.19. Fig. S19. Functional and structural features within mHAR disease circuitry** *(related to Fig. 7)***.**

**(A)** Schematic illustration for the evaluation of enhancer-promoter associations.
**(B)** Network analysis of GeneHancer modules associated with mHAR genomic hubs. Density of interactions by degree for distal (enhancers) and proximal (promoters) regulatory regions. Lower panel shows multi-hub cooperative (type A) and isolated non-cooperative elements (type B).
**(C)** Network-based ranking of connectivity (by both pagerank and eigencentrality) for ATG4C, DGAT1 and PCSK9 regulatory modules. Highlighted with a dotted circle, promoter regions with few connections (type B), and in green, regions with strong genetic association to lipid traits.
**(D)** Gene expression of type B genes (top lowest score for connectivity from C) in human adipocytes after glucose deprivation (12h) comparing control and key structural mutations. Top common functional signatures of type B promoters in both modules and active promoter genes.
**(E)** MDA lipid peroxidation relative levels in human primary adipocytes after glucose starvation (12h), and comparing control mutation, HAR genome editing (a) and editing of key structural enhancer (b) within ATG4C-hub. ANGPTL3 and PCSK9 were knocked down using siRNA.
**(F)** Fold-to-fold change correlation between MDA lipid peroxidation levels and gene (Il6 and Tnf) expression in the same conditions as in E.
Cell experiments were done with 3 independent replicates. Graphs show mean values and SEM. Unpaired, two-tailed student's t-test was used when two groups were compared, and ANOVA followed by Fisher's least significant difference (LSD) test for *post hoc* comparisons for multiple groups. * p-value <0.05.

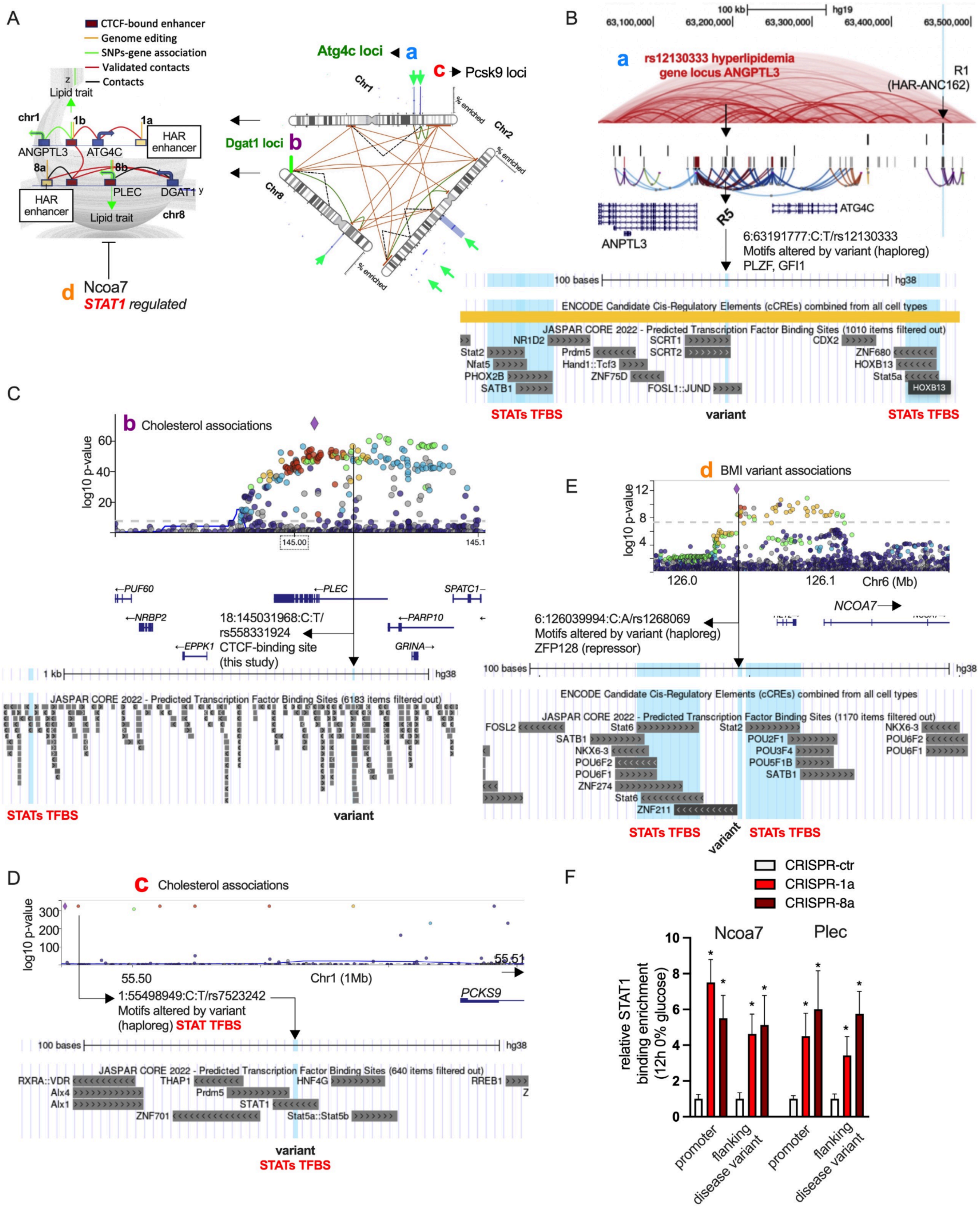

A
CTCF-bound enhancer
Genome editing
SNPs-gene association
Validated contacts
Contacts
Lipid trait
chr1
1b
1a
ANGPTL3
ATG4C
HAR enhancer
8a
8b
PLEC
DGAT1
HAR enhancer
Lipid trait
chr8
d Ncoa7
STAT1 regulated
Atg4c loci a
c Pcsk9 loci
Dgat1 loci b
Chr1
Chr2
Chr8
% enriched
B
rs12130333 hyperlipidemia gene locus ANGPTL3
R1 (HAR-ANC162)
R5
ANPTL3
ATG4C
6:63191777:C:T/rs12130333
Motifs altered by variant (haploreg)
PLZF, GFI1
STATs TFBS
variant
STATs TFBS
C
b Cholesterol associations
log10 p-value
18:145031968:C:T/ rs558331924 CTCF-binding site (this study)
STATs TFBS
variant
D
c Cholesterol associations
1:55498949:C:T/rs7523242
Motifs altered by variant (haploreg) STAT TFBS
PCKS9
variant
STATs TFBS
E
d BMI variant associations
Chr6 (Mb)
NCOA7
6:126039994:C:A/rs1268069
Motifs altered by variant (haploreg)
ZFP128 (repressor)
STATs TFBS
variant
STATs TFBS
F
CRISPR-ctr
CRISPR-1a
CRISPR-8a
Ncoa7
Plec
relative STAT1 binding enrichment (12h 0% glucose)
promoter
flanking disease variant
promoter
flanking disease variant

**1.20. Fig. S20. Dissection of disease variant circuitry within the mHAR axis** *(related to Fig. 7)***.**

**(A)** Schematic illustration for the evaluation of disease-variant circuitry. mHAR associated nuclear compartment shows functional regions (a to d) linked to molecular adaptations to nutrient stress and associated with disease-variants. Dissection of local features and screening of perturbations.
**(B)** Human genome track showing the Atg4c-loci mHAR hub, the HAR-element, regulatory elements (R1 to R7) and interactions (from GeneHancer), GM12878-Hi-C data, promoter locations, and CTCF-bound enhancers (found at the human active enhancer to interpret regulatory variants database; HACER). Motifs altered by disease variant by HaploReg. Lower panel zooms in the variant and shows predicted motifs.
**(C)** LocusZoom picture for significant disease-variants in proximity to the Dgat1-loci (chr8) from the common metabolic disease knowledge portal. With an arrow selected variant with a CTCF-motif and associated with dyslipidemia as described in A. Lower panel shows a genome track picture of regions neighboring to the variant as well as predicted motifs.
**(D)** LocusZoom picture for significant disease-variants in proximity to the Pcsk9-loci (chr1) from the common metabolic disease knowledge portal. With an arrow selected variant associated with dyslipidemia and motif altered in HaploReg. Lower panel shows a genome track picture of regions neighboring to the variant as well as predicted motifs.
**(E)** LocusZoom picture for significant disease-variants in proximity to the Ncoa7-loci (chr6) from the common metabolic disease knowledge portal. With an arrow selected variant associated with metabolic disturbances such as body mass index and motif altered in HaploReg. Lower panel shows a genome track picture of regions neighboring to the variant as well as predicted motifs.
**(F)** STAT1 binding enrichment by ChIP-qPCRs on selected regions, flanking disease-associated variants in genome edited cells (as in A) and after glucose deprivation (12h)
Cell experiments were done with 3 independent replicates. Graphs show mean values and SEM. Unpaired, two-tailed student's t-test was used when two groups were compared, and ANOVA followed by Fisher's least significant difference (LSD) test for *post hoc* comparisons for multiple groups. * p-value $<0.05$.

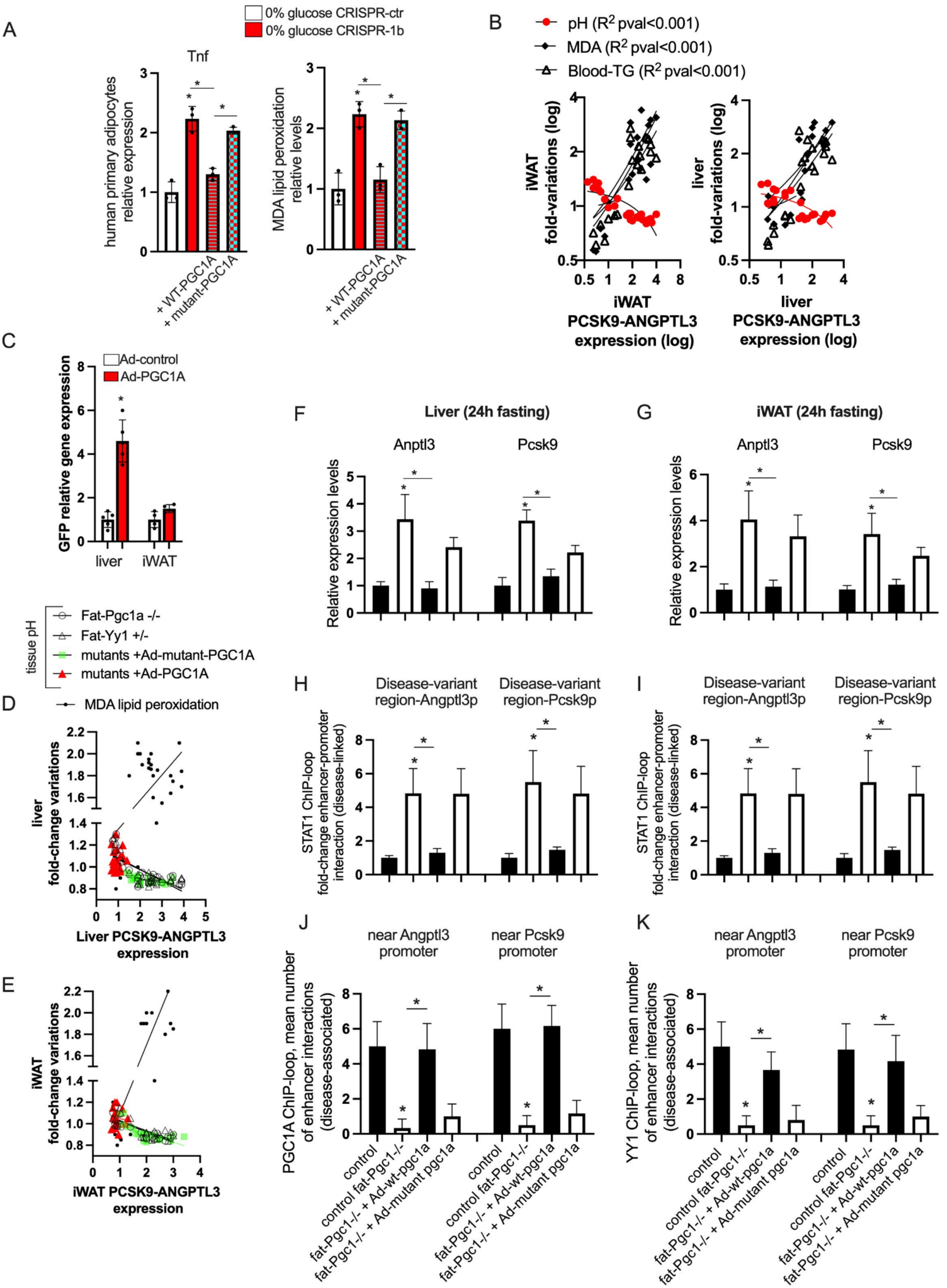
A
0% glucose CRISPR-ctr
0% glucose CRISPR-1b
Tnf
human primary adipocytes relative expression
MDA lipid peroxidation relative levels
+ WT-PGC1A
+ mutant-PGC1A
B
pH ($R^2$ pval<0.001)
MDA ($R^2$ pval<0.001)
Blood-TG ($R^2$ pval<0.001)
iWAT fold-variations (log)
iWAT PCSK9-ANGPTL3 expression (log)
liver fold-variations (log)
liver PCSK9-ANGPTL3 expression (log)
C
Ad-control
Ad-PGC1A
GFP relative gene expression
liver
iWAT
tissue pH
Fat-Pgc1a -/-
Fat-Yy1 +/-
mutants +Ad-mutant-PGC1A
mutants +Ad-PGC1A
MDA lipid peroxidation
D
liver fold-change variations
Liver PCSK9-ANGPTL3 expression
E
iWAT fold-change variations
iWAT PCSK9-ANGPTL3 expression
F
Liver (24h fasting)
Anptl3
Pcsk9
Relative expression levels
G
iWAT (24h fasting)
H
Disease-variant region-Angptl3p
Disease-variant region-Pcsk9p
STAT1 ChIP-loop fold-change enhancer-promoter interaction (disease-linked)
I
J
near Angptl3 promoter
near Pcsk9 promoter
PGC1A ChIP-loop, mean number of enhancer interactions (disease-associated)
control
control fat-Pgc1-/-
fat-Pgc1-/- + Ad-wt-pgc1a
fat-Pgc1-/- + Ad-mutant pgc1a
K
YY1 ChIP-loop, mean number of enhancer interactions (disease-associated)

**1.21. Fig. S21. In vivo dissection of disease variant circuitry** *(related to Fig. 7)***.**

**(A)** Left panel, Tnf relative gene expression in human primary adipocytes under glucose starvation and comparing the effect of genome editing of the key enhancer (disease- and mHAR-associated, as described in Fig S20A). Cells transfected with either wildtype PGC1A or IDR-mutant PGC1A plasmids. Other cells were transfected with empty plasmids as control. Right panel, MDA lipid peroxidation relative levels in the same conditions.
**(B)** Fold-to-fold change correlations in inguinal fat (iWAT, left panel) and liver (right panel) between PCSK9, ANGPTL3 expression, and lipid-related parameters and tissue pH levels in mice models used across this paper after a fasting challenge of 24 hours.
**(C)** Gfp expression levels in iWAT and liver 5 days after tail vein injection of control and Adenovirus-PGC1A constructs. Adenoviral constructs contain a Gfp gene under the regulation of another promoter.
**(D)** Fold-to-fold change correlations in liver from fat-specific mutant mice models after fasting (24h). In red, mice with adenoviral delivery (tail vein injections) of wildtype PGC1A vector. In green, mice with adenoviral delivery of mutant PGC1A vector.
**(E)** As in D, fold-to-fold change correlations in inguinal fat from fat-specific mutant mice models after fasting (24h) and after adenoviral delivery of wildtype and IDR-mutant PGC1A vectors.
**(F)** STAT1 ChIP-loop, fold-change enhancer-promoter interaction (orthologue enhancer with disease-associated variant) after fasting (24h) in liver from control, fat-Pgc1-/- mice, and mutant mice with delivery of adenovirus-PGC1A constructs as described in D.
**(G)** STAT1 ChIP-loop, fold-change enhancer-promoter interaction (as in G) and in inguinal fat from mice as in G.
**(H)** Angptl3 and Pcsk9 expression levels in liver from mice described in G.
**(I)** Angptl3 and Pcsk9 expression levels in inguinal fat from mice described in G.
**(J)** PGC1A ChIP-loop, mean number of local interactions for the enhancer with disease-associated variants in the liver from fasting mice as described in G. Positive regulatory region interactions have a fold-change over control of (FC) >2 and a p-value <0.01.
**(K)** PGC1A ChIP-loop, mean number of local interactions for the enhancer with disease-associated variants in the inguinal fat from fasting mice as described in G. Positive regulatory region interactions have a fold-change over control of (FC) >2 and a p-value <0.01.
Animal experiments were done with n=5-6 mice per group, both female and male mice were included. Cell experiments were done with 3 independent replicates. Graphs show mean values and SEM. Unpaired, two-tailed student's t-test was used when two groups were compared, and ANOVA followed by fisher's least significant difference (LSD) test for post hoc comparisons for multiple groups. Chi-square test was used to determine the statistical significance between expected and observed frequencies. Correlations were calculated by Spearman rank with a two-tailed test. * p-value <0.05.

#### 2. Supplementary tables

##### 2.1. Table S1. Metabolic phenotype comparison and human specific molecular signatures

2.1.A. Metabolic parameters and fasting endurance score in nonprimate mammals
2.1.B. Metabolic parameters and fasting endurance score in primate mammals
2.1.C. Metabolic parameters and fasting endurance score in hominoids
2.1.D. Tissue mass and metabolic parameters across mammals
2.1.E. GREAT analysis from ATACseq comparison between human-primate in fat: human increase
2.1.F. GREAT analysis from ATACseq comparison between human-primate in fat: human decrease
2.1.G. Fat branch-specific acceleration and positive selection from ATACseq comparison between humans and primates
2.1.H. Accelerated evolution signature genes in brain
2.1.I. List of human accelerated regions coordinates
2.1.J. HAR-associated genes discovery from Capra et al. 2013
2.1.K. Full list of HAR-genes compiled for this study
2.1.L. Functional annotation of HAR genes in brain-specific networks
2.1.M. Functional annotation of HAR genes in fat-specific networks
2.1.N. Data

##### 2.2. Table S2. Transcriptional regulatory networks of HAR-genes

2.2.A. ChIP-seq enrichment ChIP-atlas of HAR-genes
2.2.B. Adipocytes ChIP-seq enrichment ChIP-atlas of HAR-genes
2.2.C. Neural ChIP-seq enrichment ChIP-atlas of HAR-genes
2.2.D. Stem cells ChIP-seq enrichment ChIP-atlas of HAR-genes
2.2.E. Blood ChIP-seq enrichment ChIP-atlas of HAR-genes
2.2.F. Muscle ChIP-seq enrichment ChIP-atlas of HAR-genes
2.2.G. Liver ChIP-seq enrichment ChIP-atlas of HAR-genes
2.2.H. TFBS enrichment from MSDB
2.2.I. TFBS enrichment form genome browser
2.2.J. ENCODE and ChEA consensus enrichment
2.2.K. Data

##### 2.3. Table S3. Transcriptional regulatory networks, and PPI reconstitution

2.3.A. Node + 1 network reconstruction of PPIs for HAR-genes regulators
2.3.B. Network topological prioritization of network in A
2.3.C. Full PGC1A-network reconstruction of protein-protein interactions
2.3.D. Network topological prioritization of full PGC1A-network
2.3.E. Reactome functional annotation of HAR cooperative regulators
2.3.F. Gene-ontology functional annotation of HAR cooperative regulators
2.3.G. Fat and brain tissue-specific PPIs for PGC1A-network
2.3.H. Data

##### 2.4. Table S4. Transcriptional regulation of HAR-genes and metabolic-HAR hubs

2.4.A. Functionally-associated and interacting transcriptional regulators: co-regulated genes by promoter ChIP-seq enrichment
2.4.B. Co-bound transcriptional regulators: co-regulated HAR-genes (metabolic-HAR genes)
2.4.C. Positional gene enrichments of metabolic-HAR genes
2.4.D. Cytogenetic band enrichments of metabolic-HAR genes
2.4.E. Functional annotation of metabolic-HAR PGE-domains
2.4.F. Data

##### 2.5. Table S5. Compendia of HiC data and hESC interacting interchromosomal regions

2.5.A. HiC chromatin interacting regions
2.5.B. Frequently interacting regions (FIREs)
2.5.C. Network from hESC-HiC interacting chromatin segments and metabolic-HAR regions overlap
2.5.D. Network ID of interacting segments in A
2.5.E. Topological prioritization and network analysis of hESC-HiC interacting chromatin segments
2.5.F. Data

**2.6. Table S6. Genome-wide network of enhancer-promoter relationships**

2.6.A. GeneHancer database of enhancer-promoter associations.
2.6.B. Regulatory elements associations
2.6.C. Network analyses of regulatory elements associations
2.6.D. Module-based and degree prioritization
2.6.E. Degree-based prioritization of enhancers
2.6.F. Social promoter genes and genomic cluster positions
2.6.G. Nonsocial promoter genes and genomic cluster positions
2.6.H. Data

**2.7. Table S7. mHAR genomic hubs subnetworks from linked transcriptional compartments**

2.7.A. ATG4C-submodule short-range relationships
2.7.B. DGAT1-submodule short-range relationships
2.7.C. PGC1A-submodule short-range relationships
2.7.D. CAT-submodule short-range relationships
2.7.E. ATG4C-submodule territorial ChIP-seq enrichment for social enhancer and promoters
2.7.F. DGAT-submodule territorial ChIP-seq enrichment for social enhancer and promoters
2.7.G. PGC1A-submodule territorial ChIP-seq enrichment for social enhancer and promoters
2.7.H. CAT-submodule short-range relationships
2.7.I. PPI network reconstruction of territorial ChIP-seq enrichment of regulators within ATG4C submodule
2.7.J. Data

**2.8. Table S8. Genomic range genetics of metabolic-HAR compartments**

2.8.A. ATG4C metabolic-HAR domain genomic range genetics (chr1:62mb)
2.8.B. INSIG2 metabolic-HAR domain genomic range genetics (chr2:110mb)
2.8.C. PGC1A metabolic-HAR domain genomic range genetics (chr4:25mb)
2.8.D. CAT metabolic-HAR domain genomic range genetics (chr8:38mb)
2.8.E. APOA5 metabolic-HAR domain genomic range genetics (chr11:117mb)
2.8.F. Data

**2.9. Table S9. Synteny of metabolic-HAR regions and mammalian perturbation networks**

2.9.A. Murine transcriptional hubs of metabolic-HAR regions
2.9.B. Metabolic-HAR gene perturbation-phenotype association
2.9.C. Gene network topological dependencies
2.9.D. Phenotype network topological dependencies
2.9.E. Data

**2.10. Table S10. ChIP-MS structuring candidates and PGC1A-network**

2.10.A. List of proteins identified by chromatin proteomic profiling in mESC dual enhancer-promoter
2.10.B. Interrogation of PPIs for PGC1A-network and ChIP-MS structuring candidates
2.10.C. IDs node summary annotation for B
2.10.D. Functional network annotation
2.10.E. Network topological dependencies
2.10.F. Data

**2.11. Table S11. Intrinsically disordered regions and conservation of HAR-gene regulator**

2.11.A. Consensus disordered regions in PGC1A protein
2.11.B. Hydropathy boundary ordered and disordered proteins
2.11.C. Multivalent pi-pi disordered score human proteins
2.11.D. DisProt disordered score human proteins
2.11.E. Consurf amino acid variation of PGC1A residues
2.11.F. Amino acid conservation scores for PGC1A co-activator
2.11.G. Data

**2.12. Table S12. Summary table of published datasets used for integration and analysis**

2.12.A. Link

#### 3. Computational materials and methods: functional genomics framework

##### 3.1. Introduction

The genome organization facilitates the coordinated integration of multiple transcriptional components for the regulation of gene expression, enabling cell adaptation and functions. Within the nucleus, distinct chromosome territories, long- and short-range chromatin interactions, and nearby transcriptional factories, guide the formation of dynamic transcriptional compartments (*10*, *11*). Interestingly, it has been suggested that these transcription factories are often located near interacting chromosome regions, which act as anchors, enabling the co-localization of key regulators for the co-regulation of distinct gene programs (*11–13*). This hypothesis has been supported by observations of non-random spatial proximity between adjacent chromosomes, where DNA translocation is influenced by contact points (*14*).

Moreover, regions in contiguous or interacting chromatin hubs are found to be regulated by local transcriptional factories (*15*). These chromatin interactions can be increased or reduced in a tissue- or cell-specific way, thus helping define cell-specific transcriptional factories (*15*). These features indicate a high-degree of plasticity, likely depending on the presence or absence of cell-specific transcriptional regulators (*12*). This plasticity is substantiated by the identification of spatially defined chromatin modules via high-throughput profiling (*16*), and by the conservation of higher-order structural data, such as topologically associating domain (TAD) boundaries, frequently interacting chromatin regions (FIREs) (*17*), and trans-interacting regions (*15*, *18*). The level of interactions and functionality of chromatin architecture likely rely on the activity of cell-specific transcription factors. These factors drive the functional interactions of chromatin modules and dictate the location of nuclear speckles within transcriptional factories (*19*). Supporting this notion, transcriptional factories have also been discovered near trans-interacting domains (*12*). Furthermore, genome-wide crosslinked methods that map higher-order interacting regions have revealed that the physical constraints within the nucleus limit these domains to relatively large interacting hub regions (*11–13*). These findings underscore the level of conservation and plasticity of interacting domains across diverse tissues and cells and provide clues for the targeted identification of cell-specific, functionally-distinctive, or evolutionary-tuned transcriptional compartments.

##### 3.2. Description

Given the previous definitions regarding gene co-regulation through the formation of transcriptional compartments, our approach is based on 3 main components: 1) identifying genomic hubs (regions or segments) from gene sets that are either naive or functionally enriched (biological function, transcriptional co dependencies, or cell-enriched), 2) evaluating structural and functional relationships in these genomic hubs, and 3) selecting those with multimodal dependencies for experimental interrogation. Our framework is flexible and can be scaled to other gene regulatory networks. We first define gene sets with transcriptional dependencies by identifying transcriptional co-regulators. This is followed by determining genomic hubs by the positional clustering of these gene sets. We next categorized genomic hubs whose local genes share functional enrichments, and those sharing long- and short-range structural relationships. Regarding the latter and in order to map possible structural combinations, we used a naive approach focusing on interacting chromosome regions and territories. This approach is coupled with a network-based integration, which is used to generate maps with structural relationships suited for downstream computational and experimental interrogation. To investigate the structural dependencies within genomic hubs, we defined interacting regions using Hi-C data from multi-tissue and multi-cell type compendia, including a network-based model of interacting regions. This is combined with local structural data using enhancer-promoter associations from curated and integrated datasets such as GeneHancer repository. With these steps, we identify interacting regions (territories) and local chromatin modules (associated enhancer-promoter modules). For genomic hubs sharing structural dependencies, we evaluate genetic trait enrichments locally and within the associated transcriptional compartment. We further prioritize genomic hubs that are conserved or syntenic in mice and those whose genes share similar phenotypes from knockout repositories. We finally complement this by evaluating computationally the conformational features, disordered regions, and interactions between co-regulators. This is followed by in depth experimental validation of transcriptional multimodal components in model cells and organisms.

##### 3.3. Description of implementation

To study gene sets associated with human accelerated regions (HARs), we reviewed and collected information regarding genes in proximity and linked to conserved regulatory elements with a high rate of human-specific mutations (*20*, *21*) (**fig. S1A, B**). Our computational and experimental framework aims to understand the multifaceted components of transcriptional regulation and activity of HAR-associated regions. We first identify HAR-associated gene subsets with functional, transcriptional, and cell-type enrichments, and locate regions where these genes cluster in HAR-associated genomic hubs (**fig. S1C**). We use positional genomic clustering of metabolic HAR genes to identify hubs, apply functional annotation to each genomic hub, and explore their structural relationships using chromatin architecture data (**fig. S1C, D**). We evaluated both long- and short-range relationships from publicly available compendiums, which allowed us to prioritize genomic hubs with both functional and structural relationships. For example, we identified interacting genomic regions harboring HAR genes with functional and transcriptional dependencies, and consolidated them into nuclear compartments based on spatial proximity and chromosomal territories (**fig. S1D**). From these higher-order structures we then evaluated local short-range relationships using enhancer-promoter associations (**fig. S1E**). In each genomic hub with nuclear proximity from 3C-based approaches, we obtained local neighborhoods or chromatin modules using a network-based approach, in which topologically associated regulatory regions (from enhancer-promoter associations, data stored at GeneHancer) were identified and interrogated for ChIP-seq binding enrichments using repositories (**fig. S1E**). Among regulators with binding enrichments in local chromatin modules, we confirmed their functional enrichments and physical interactions (using protein-protein interactions repositories followed by network modeling; **fig. S1F**). We further complemented this by assessing the phenotype association of genetic variants enriched in these hubs (**fig. S1F**) and the phenotype conservation from murine gene knockouts enriched in syntenic HAR-associated genomic hubs (**fig. S1G**). Additionally, we assessed the tissue and cell-type enrichment of genes forming hubs in publicly available atlases. For interacting and co-binding transcriptional regulators of metabolic HAR genomic hubs and genes, we evaluated conformational features and disordered regions from amino acid sequences (**fig. S1F**). Once features of interest were identified, we interrogated the evolutionary conservation of relevant features. After identifying structural and functional relationships of interest, we proceeded to experimentally validate functionally-enriched nuclear compartments, their cooperative regulators, their condensate plasticity, and their phenotype associations in murine and human models (**fig. S1H**). In the following sections, we will describe the composing steps of our pipeline.

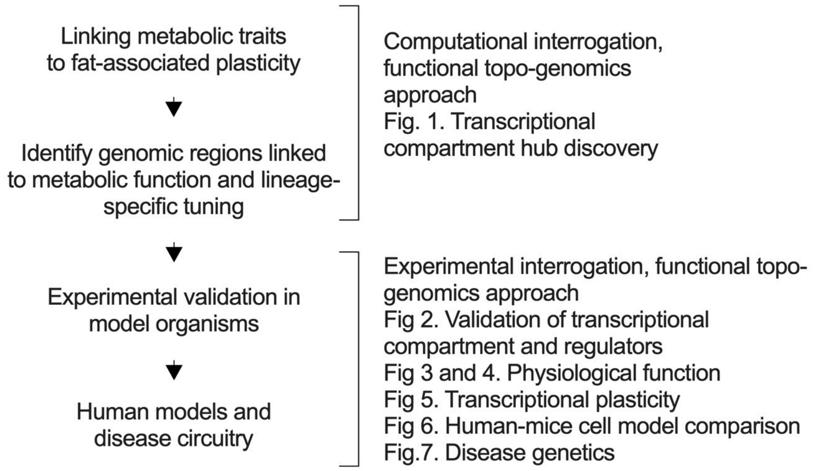

Summary illustration: workflow overview and figure content index.

##### 3.4. Comparison of metabolic phenotypes in mammals

Data for multi-species comparisons of metabolic phenotype were obtained as follows: total energy expenditure (TEE) from, including hominidae fat and lean mass, and (*1*, *2*), experimental fat mass measurement across mammals (*22*) (table S1).

##### 3.5. Fasting endurance score (theoretical survival time)

To calculate energy endurance score, we followed previous work by Lindstedt et al. (*23*), where they generated a fasting endurance score (theoretical survival time in nutritional deficit) as a function of body size in mammals. We used a similar approach including allometric (10% of body weight) and experimentally measured fat mass. Briefly, the theoretical survival time or endurance score is based on the relationship between fat mass energy content (in kilojoules) and the amount of energy consumed per day in the form of total-energy-expenditure (TEE) collected from previous reports and shown in table S2. This measure represents resource availability, assuming energy intake is low. This gives a representation of the number of days a given species can last solely on the energy stored in fat tissue.

To calculate the theoretical survival time for each species, we first calculate the energy content in the fat mass (assuming fat mass is 10% of body mass) and then divide that by the TEE (kcal/d) for each species.

- Energy content in fat = Fat mass (kg) * 7700 kcal/kg (1 g of fat contains approximately 7.7 kcal)
- Theoretical survival time (days) = Energy content in fat (kcal) / TEE (kcal/d)

After obtaining the theoretical survival time or endurance score for each species (wherever data was available), we plot this as a XY graph together with fat mass. Fat mass is derived from both experimentally measured and allometric fat mass. We then performed linear regression to compare both experimental and allometric models, followed by F-test model comparison.

##### 3.6. Comparison of primate and human molecular signatures

We used previously published work to compare primate and human data: from brain cells (*8*) and adipose tissue (*7*). We re-analyzed the data following protocols described there (GEO: GSM3494237–GSM3494249), and used differential changes for downstream analysis.

##### 3.7. Human accelerated regions (HAR) associated genes

Human accelerated regions and HAR associated genes were obtained from previous work (*20*) (table S3). Full HAR-associated genes were completed with subsequent studies (*21*). Enhancer coordinates and HAR-genes were used for downstream analysis. HAR-genes were used as input for network-based module discovery in specific tissues, followed by interrogation of overlapping molecular signatures with results from primate-human high-throughput comparisons.

##### 3.8. Network-based module discovery in pre-built tissue-specific networks

To assess functional gene dependencies in tissue-specific networks, we performed a targeted approach of overlapping network representation using data from tissue-integrated genome-scale analysis (*6*) (archived in http://giant.princeton.edu and https://hb.flatironinstitute.org/). This was followed by functional clustering of dependencies.

##### 3.9. Relationships in transcriptional regulators

###### 3.9.A. Promoter-binding transcriptional regulators

To assess transcriptional binding to promoters of HAR-associated genes, we used a combined enrichment approach including motifs and binding of regulators. For a given geneset we used conserved common regulatory motifs enrichment (archived in http://www.gsea-msigdb.org/gsea/msigdb/index.jsp). Enrichr in R was used to perform functional enrichment analysis based on the database: ChEA 2016 and ENCODE consensus. FDR <0.01 and p-value <0.05 were used as a threshold to select the significant enrichments (*6*, *24*) (archived in https://maayanlab.cloud/Enrichr/). Enrichment results from each class were followed by a venn-diagram of overlapping regulators. These were used for downstream analysis. Prediction tool for TFBS from DNA sequences can be found at http://alggen.lsi.upc.es/. Tool for prediction of CTCF-binding sites can be found at https://insulatordb.uthsc.edu/ (*9*).

###### 3.9.B. Identification of interacting transcriptional regulators

Common regulators from our previous step were used as inputs to identify physical interactions in Protein-Protein interaction (PPI) data stored in several repositories. First, we obtained physical interactions from regulators using human protein interaction atlas (archived in http://www.interactome-atlas.org/ (*25*)). This was followed by filtering of regulators displaying common molecular signatures. These were used for network reconstitution, modeling, and analysis.

###### 3.9.C. Network reconstruction and modeling

Next, we carried out a 1+ node reconstruction (partial) and full network reconstruction of all possible interactors from our filtered targets. To do so, we used both HuRI and Biogrid PPI-data (archived in https://thebiogrid.org/ (*26*)). Partial and fully reconstituted networks were used for downstream analysis. We performed global and local network analysis with clustering and modularity followed by node-prioritization with eigenvector centrality and random-walk algorithms. Nodes with the highest influence by network-ranking were used for functional annotation embedding using DAVID clustering (*27*) and HAR-association with our curated dataset. Top ranking nodes were filtered for downstream analysis.

##### 3.10. Binding combinations of interacting transcriptional regulators

To assess binding combinations of interacting regulators with similar function to promoters of HAR-associated genes, we used the previously described (*28*) ChIP-seq database containing more than 100 thousand experiments (archived in https://chip-atlas.org/). First, we selected top ranked nodes from the interacting transcriptional regulatory network. Next, we parsed and integrated all ChIP-seq experiments for each regulator, binding ±10kb from the transcription start site (TSS). The peak calls in each experiment are based on MACS2 score as previously described (*28*), then we use the average MACS2 score from the available experiments. To filter out all possible combinations of co-regulated target genes, we used a cutoff with an average score higher than 50. Next, we built overlapping maps using venn-diagrams in R from filtered target genes. With this, we quantified the number of co-regulated genes by pairwise combinations of interacting regulators. We used these frequent combinations to select transcriptional regulators with most associations and derive a combination-score for a given gene set. As background sets are all possible regulator combinations with all possible target genes. We quantified the number of genes for each combination (classes are >2, >4, >6, and >8 transcriptional regulator combinations). The combination score for the background set is the log fraction between the number of combinations in each combination class and the number of overlapping target genes (comb score = # of combinations / number of overlapping genes for that combination class). We evaluated the combination score for HAR-associated genes as follows: HAR-genes comb-score is the log fraction between the number of combinations in each binding combination class and the overlapping HAR-associated genes (number of HAR-genes with >2 combinations). We compared our target gene set with random sampling gene sets with the same number of genes to our query set. Finally, we performed hypergeometric enrichment to compare our query to the background and random sampling, and defined statistical significance at p-value <0.01. HAR-associated genes bound by several combinations of interacting regulators were annotated using EnrichR in R followed by selection of common biological terms.

##### 3.11. In silico phase separation scores

We identified intrinsically disordered regions in the amino acid sequences of transcriptional regulators. Disorder scores from protein amino acid sequences were obtained using predictors of naturally disorder regions software (*29*, *30*), inference of intrinsically unstructured proteins (IUPred) with Anchor (binding regions within IDRs) package (*31*), multivalent pi-pi interactions in non-aromatic residues of folded proteins (*32*) and curated experimentally validated phase separation databases (stored at http://www.pondr.com/, https://iupred3.elte.hu/, https://mobidb.bio.unipd.it/ and http://db.phasep.pro/) (*33*). From PONDR, we obtained VSL2- and hydropathy-scores plotted against mean net charge of proteins of interest. The hydropathy-score in PONDR uses a linear discriminate function to define boundaries between disordered and ordered proteins (*34*).

##### 3.12. CIDER package: identification of context-dependent disordered regions

Physical conformations between intrinsically disordered proteins vary depending on different principles and environment (*35*). Interestingly, solvent-mediated electrostatic repulsions and attractions have been shown to depend on linear sequence conformations and charge segregation within disordered regions(*36*). Then, the proportion of linear sequences with charge segregation in IDRs partitioned structural representations of proteins into categories such as globules, context-dependent globules, coils and chimeras. These conformations form an ensemble of parameters for new phase separation classifiers that complement well established low-complexity regions predictors. We used classification of intrinsically disordered ensemble regions (CIDER) to obtain parameters related to phase conformational transitions (archived in http://pappulab.wustl.edu/CIDER/about/). We implemented local-CIDER in python 3 (stored at http://pappulab.github.io/localCIDER/), to compute bulk- conformational variations in large data samples. We first focused on globules and context-dependent globule conformation classification (*36*). These were shown to be influenced by environmental variations such as salt, charge, ligand-binding, cis-interactions and solvation (*35*). Among the parameters obtained that are at the basis of their conformation classifier, charge-mediated transitions can be recapitulated by net charge and mixing of opposing residues within IDRs. A list of parameters evaluated by CIDER is as follows: K or Kappa represents segregation of opposing charge residues with 0 being well mixed or intertwined while 1 being separated stretches of positive or negative residues; Fraction of charge residues (FCR); net charge per residues (NCPR); f- negative fraction; f+ positive fraction; hydropathy (*37*) and disorder promoting scores. Fasta files of protein sequences of interest were downloaded from the NCBI repository.

##### 3.13. Consurf package: phylogenetic amino acid conservation and structural features

Phylogenetic analysis was performed on amino acid sequences of proteins of interest such as PGC1A. We used the ConSurf software package to obtain conservation scores and multi-species comparison (*38*). Briefly, it estimates the degree of conservation of amino acid sequences based on iterative comparison with homologous sequences. The conservation rate of a sequence depends on its importance for the structural function. The evolutionary relationship is dictated by the similarity between species comparison in the substitution matrix using previously established Bayesian or maximum likelihood (ML) approaches (*39*). Here below is a brief description of ConSurf stepwise analysis. First, consurf uses a heuristic algorithm to search default homologous sequences using BLAST across different databases and with known 3D structural features (*40*). A multiple sequence alignment is followed by a phylogenetic tree analysis using neighbor-joining algorithms (*39*). Conservation score or evolutionary rate in each residue is imputed using rate4site, and divided in different scales for the degree of preservation. Residues evolving slowly are referred to as conserved and those evolving rapidly are referred to as variable. The degree of preservation depends on different levels of purifying selection, which can be related to folding constraints, enzymatic activity, ligand-binding or protein interactions. For the rate of evolution, the Bayesian approach takes into account the stochasticity of evolutionary change and the phylogenetic comparison (*41*). This probabilistic model of amino acid replacement estimates how likely a residue is to influence structure and function of protein domains. 3D structures of proteins are obtained from updated databases such as the protein data bank (*42*). It uses simulation based estimation of amino acid substitutions within the tree branch and compares them to known protein structures (*41*). For each residue within the sequence, consurf gives a normalized score of evolutionary conservation from 1 with lowest conservation to 9 with the highest conservation. In addition, consurf maps the estimated conservation score within proteins, to known 3D models of protein structure (*42*). It uses HHPred, which exploits a hidden markov model to look for 3D templates from homologous sequences or similar proteins using PDB (*43*) and extrapolate them to the query sequence with structural features. The features for each residue are the degree of 3D exposure, hiddenness and functionality. The evolutionary phylogenetic tree is calculated using the WASABI platform (*44*) which is integrated with multi-species display of amino acid sequences and the degree of evolutionary conservation based on the aforementioned methods. We evaluated the phylogenetic tree and the degree of conservation for PGC1A sequence. We obtained for each amino acid a conservation scale, with structural features from known 3D representations. We used this to compare the degree of preservation of intrinsically disordered regions within PGC1A. The IDRs were defined as consensus disordered domains within the protein. We extracted structural features and the degree of evolutionary rate. To compare IDR domains with other protein domains we defined consurf estimations as follows: conserved residues have a score of 7 or higher, while low conservation residues have scores lower than 3. As previously described, from the consensus disordered predictions, we obtained 2 IDR domains in PGC1A. One close to the N-terminal and the other in proximity to the RNA-binding domains. For both IDRs we quantified the number of residues with low conservation score, high conservation, and average conservation. We then obtained the proportion for each category relative to the total number of residues. We performed the same approach for the rest of the protein and compared the estimation of evolutionary rate between the domains. A similar approach was carried out to assess evolutionary innovation in charged residues within the IDR domains. Final scoring was obtained as the proportion of charged residues (positively charged: arginine[R], lysine[K], and negatively charged: glutamate[E], aspartate[D]) within IDRs with low conservation score, in relation to the total number of residues.

##### 3.14. ChIP-Mass spectrometry structuring candidates

To evaluate regulators that bind active enhancers and promoters and their relationships with PGC1A-network, we used previously published datasets (*45*), and a similar implementation (*46*). Briefly, structuring candidates were identified with chromatin immunoprecipitation followed by mass spectrometry. Antibodies targeting epigenetic tags of active enhancers (H3K27ac) and promoters (H3K4me3) (*47*), were used to precipitate binding regulators. Mass spectrometry was used in those complexes to identify putative candidates. As previously described by Weintraub et al. binding candidates were filtered using the log2 ratio of the IP over the IgG with a cutoff of $\log 2 > 1$.

##### 3.15. PPIs structuring candidates and PGC1A-associated network

For the resulting structuring candidates, we performed network modeling with PGC1A-network candidates (from transcriptional relationship analysis described before). To this end, we used protein-protein-interaction reference networks (at HuRi) filtered for adipose specific enrichments. Network recomposition (node + 1 interactor) was performed for all direct interactors in both datasets, and network analysis parameters and functional enrichment were obtained as previously described (see transcriptional relationships).

##### 3.16. Identification and functional annotation of genomic hubs by positional gene enrichments (PGE)

To identify genomic hubs from the positional location of genes of interest, we used previously established positional gene enrichment methods (*43*, *48*). Briefly, PGEs exploit topological features of gene locations to determine chromatin regions enrichments. This approach is based on calculating the hypergeometric distribution along genomic distances for gene sets. For a given region, it determines the probability of having observed genes in that region. To test statistical significance, cumulative p-value distributions are used on random simulations or false discovery rate on large gene-sets. The probability of achieving p-value enrichments that are better than chance estimates are used to define hub enrichments. The additional constraints to categorize a genomic region displaying positional enrichments for a given set of genes are as follows: having at least 2 genes; that no smaller region was found by random permutations and cumulative p value distributions; that no bigger regions with more target genes were found. This algorithm defines genomic regions by distance of query genes, followed by estimation of p-value distribution to compare chance expectation. Target genomic ranges for positional enrichment were set to 15-million base pairs (following structural relationships on interacting homotypic TADs), where PGEs with larger domains were filtered out. To obtain qualitative PGEs we used regions that were enriched within cytogenetic bands (*49*). To obtain a global algorithmic performance determined by the number of genes being queried, we additionally performed permutations with random gene sets (same number of inputs), followed by statistical comparisons using Chi-square test and p-value discrimination. For every topological gene cluster in the form of PGEs, we derived functional enrichments using Enrichr in R. FDR <0.01 and p-value < 0.05 was used as a threshold to select the significant enrichments for biological terms and phenotypes associated with a given region. PGEs with common biological enrichment across domain-catalogs were categorized as class fPGEs (functional PGEs hubs) for downstream analysis. These hierarchical chromatin domains were used as seeds for prioritization of structural chromatin data when indicated, and as inputs for long- and short-range structural relationships.

##### 3.17. Combined PGE enrichments

To identify colocalized enrichments or convergence of enrichment between individual gene sets and combined gene sets, we used some modifications. We compared the proportion of PGEs with >5 genes / total regions, in combined sets by group, individual sets by group and random permutations of PGEs with similar parameters. To discriminate between the average of expected proportions for individual and random combinations, with the proportions from biological combinations we used chi-square test and p-value estimation.

##### 3.18. Hi-C Data and genomic hub colocalization

Representative Hi-C data from diverse tissues and cells was procured from 4DN including hESC, human lymphoblastoid cells, colon cancer, hs2-hi-c (*3*, *50*) and uploaded using straw (*51*). A/B compartments and TADs in chromosomes were determined using previously established protocols (*3*, *52*). Briefly, boundaries were defined at 40-kb, 1-mb genomic regions, and 200-kb window for delta vector calculation. Identification of significant Hi-C contacts at 40-kb resolution was done with Fit-Hi-C (*53*). Within a 2-Mb genomic distance p- and q-value were calculated for each bin pair with FDR<1e-5 threshold for peak-calling.

###### 3.18.A. Frequently interacting regions (FIREs)

We used a published compendium of Hi-C data from 21 human tissues and primary cells, downloaded from the GEO database (*17*) (GSE87112), along with previously established protocols described therein (*17*). Briefly, data was normalized using HiCNorm (*54*), vanilla coverage (*50*) or ICE (*55*). FIREs regions were obtained as previously described by Schmitt et al (*17*). Briefly, FIRE bins were identified with a poisson regression model (HiCNormCis) as previously described (*3*, *52*). With this approach the total normalized cis-interactions were obtained, and a gaussian distribution approach was used to estimate the local cis contacts, followed by conversion of values to -ln(p-value) denoted as FIRE score and used in our pipelines for downstream analysis. Once conserved segments were obtained, these FIREs were plotted as chromosome positions. We next quantified FIREs within chromatin domains defined by PGEs hub as previously described. These chromatin regions were defined by regions derived from metabolic-HAR genes or metabo-HAR genomic hubs. We calculated the mean number of FIREs within PGEs hubs and compared it to the mean number of FIREs within PGEs from random gene sets. We define each random permutation with the same number of genes defining the metabo-HAR genomic hubs. This was followed by a chi-square test to evaluate statistical significance and area under the curve of ROC to evaluate the discrimination threshold between comparisons.

###### 3.18.B. Interacting regions in hESC Hi-C data

To identify the relationship between metabolic-HAR genomic hubs and trans-interacting regions in human chromosomes we used data from hESC (GEO: GSE35156 (*56*)). To assess the trans-interacting interchromosomal regions, we used previously established methods with a few modifications (*4*). Briefly, Hi-C data was normalized with hic-pipe, and a probabilistic model was followed to calculate chromatin segment contact maps (*57*). For contact call significance, a binomial distribution p-value estimation was used as previously described (*58*). Bins were defined at 500-kb and contact probabilities were normalized against chromosome length. An interacting network was then built with chromatin segments as nodes and edges as presence of interaction. An undirected graph was built and used to identify chromatin segments with metabo-HAR genomic hubs. Where a metabo-HAR genomic hub was present in a given interacting chromatin segment, we labeled that node as either PGE and fPGEs if they have a functional embedding (see identification and functional annotation of genomic hubs by positional gene enrichments). We quantified the number of interacting bins within chromatin segments defined by PGEs as previously described. These chromatin regions were defined by regions derived from metabolic-HAR genes or metabo-HAR genomic hubs. We next calculated the mean number of interacting bins within metabo-HAR genomic hubs and compared it to the mean number of bins within PGEs hubs identified from random gene sets (of similar number composing genes and length of chromatin segment). We define each random permutation with the same number of genes defining the metabo-HAR hubs. This was followed by a chi-square test to evaluate statistical significance and area under the curve of ROC to evaluate the discrimination threshold between comparisons.

##### 3.18.C. Network modeling and analysis of genomic hubs in chromatin interactions

An interacting network was formed with segments as nodes and edges as presence of interaction as previously described (*4*). We used the NetworkX library in python to perform subsequent topological analysis of the embedded network. We quantified average node degree, mean clustering coefficient, centrality measurements such as betweenness and closeness, information flow parameters such as bridging centrality, and node-prioritization with pagerank and eigenvector centrality. Graph layout such as force atlas, openord, circular, were used for representations when indicated. We performed randomization with similar identified parameters and assessed global structural properties within the hESC interacting network. To compare the topological location properties between PGEs and fPGEs (bio-enriched hubs), we averaged the clustering coefficients and node degrees for the overlapped chromatin segments or nodes. We compared these to full network averages and random sampling with similar number of nodes. With the embedded functional annotation obtained before, we identified top hubs and nodes with a specific functional cluster. The functional annotation of top clusters were obtained with the functional embedding of PGEs (see above). To filter nodes that both display a specific functional annotation and influence information flow through the network, we used bridging coefficient and overlapping functional annotation to hubs and robust segments. Top bridging chromatin segments harboring fPGEs of interest were selected for downstream compartment reconstruction and experimental interrogation. Complementing the network-based prioritization, we functionally annotated genes within top interacting chromatin segments using DAVID functional clustering analysis (https://david.ncifcrf.gov/) (*27*).

#### 3.19. Integration of short-range chromatin relationships

To study genome regulatory elements relationships at high resolution, we used previously reported integration methods and results on enhancer-promoter associations from the GeneHancer repository (*5*). Here, we employed enhancer-promoter relationships followed by network-based analysis. Briefly, GeneHancer map is a genome-wide regulatory regions relationships, comprising around 284000 integrated elements from different databases: ENCODE, the Ensembl regulatory build, the functional annotation of the mammalian genome (FANTOM) project, the VISTA Enhancer Browser, dbsuper super-enhancers, EPDnew species-specific databases of experimentally validated promoters, and UCNEbase ultra-conserved noncoding elements. The approach by the Cohen Lab links regulatory elements to promoter genes, using a combined strategy as follows: correlation between genes and enhancer RNAs (*59*), transcription factors binding enhancers and target genes, expression quantitative trait loci (eQTLs) (*60*), promoter-capture Hi-C (*61*), and distance-based associations on immediate adjacency (*62*). GeneHancer map uses a likelihood-based score for each enhancer–gene association. This generates a hierarchical pairing that is defined by the number of sources supporting it (*5*). Given our interest in studying both global and local topological representations and the fact that human-accelerated regions are elements that might not have known links, we included all linking data for downstream analysis.

##### 3.19.A. Genome-wide relationships in short-range linking network

Cis-interacting short-range chromatin networks were formed as an undirected graph with regulatory regions defined as follows: enhancer elements as distal regulatory regions and promoter genes as proximal regulatory regions. Regulatory regions were defined as the nodes while the edges were the presence or linking association between them. We then interpolated metabo-HAR domain information as follows: HAR-associated genes described before were labeled in the proximal regulatory regions, while non-coding HAR element coordinates were overlapped in the distal regulatory elements. We next used the NetworkX library in python or GEPHI to layout the graph and perform subsequent analysis. We quantified average node degree, mean clustering coefficient, centrality measurements such as betweenness and closeness, information flow parameters such as bridging centrality, and node-prioritization with pagerank and eigenvector centrality. We also performed a modularity based clustering approach to find communities of local regulatory-regions, and represent them as the graph layout hierarchical neighboring modularity. We performed randomization with similar identified parameters and assessed global structural properties. Using hub-based and influence-based node prioritization, we ranked regulatory regions by their density of interactions. We derived genome-wide linking interactions for proximal regulatory regions as the global mean degree. Based on this approach, here we categorized and defined this threshold as a social cutoff, and identified regulatory elements or nodes as social if they had 2 times the genome-wide mean degree (>16). Likewise, isolated regulatory elements or nodes were defined as nodes displaying < 8 degree interactions. To compare the topological location properties for HAR-associated genes, we calculated a hypergeometric enrichment from the metabo-HAR-genes on the list of social promoters derived from our genome-wide social network ranking approach. We compared this with permutations of PGEs genomic hubs built from random gene sets of similar parameters. This was followed by chi-square tests to evaluate statistical significance. To evaluate distal regulatory dependencies of non-coding active elements, we classified all regulatory elements by their density of interactions. We then compared the genome-wide behavior of enhancer linking.

##### 3.19.B. Genomic hubs local relationships in short-range linking subnetworks

Once we evaluated global relationships for the regulatory elements of interest within the short-range network, we took advantage of a sub-module clustering approach to study local structural representations, yet keeping its global dependencies. This method can be used to study any PGE-hub constructed from a given gene set. First, we selected functional-PGEs hubs (fPGE) as previously described and label the modules they are located. Briefly, top bridging trans-interacting segments harboring fPGEs that shared similar functional annotation to top hub trans-interacting segments were chosen for nuclear compartment reconstruction and further experimental validation. For the selected trans-compartments, we labeled the modules where local fPGEs hubs reside, using the short-range linking network, and used these modules for downstream analysis. To embed both network and biological qualitative information in the modules of interests, we used a social network based approach supported by epigenetics and transcriptional binding from experimental data. First, every module was further classified by sub-modules using the module cluster algorithm. This generated territorial clusters of associations for HAR-domains (or any PGE) with intra-submodular and inter-submodular interactions. We then interrogated iteratively binding enrichments of epigenetic chromatin tags given territorial associations (for each module and submodule). To do this, we used the integrative ChIP-atlas data, to calculate binding enrichments for enhancer coordinates that form topological clusters of associations. When necessary we used liftover function to match coordinates between different assemblies (USCS genome browser at https://genome.ucsc.edu/cgi-bin/hgLiftOver). To calculate background ChIP-enrichments per module, we used histone modifications (n=16641 experiments) across all cell types (n=61679 experiments), with a threshold of significance of 50 and random permutations for comparison. Data was subsequently presented as fold-enrichment to p-value followed by word-cloud generator showing most frequent histone modifications. We next embedded topological qualitative information from our network-based analysis to classify social and isolated enhancers (see relationships in short-range linking networks). We differentiated chromatin states between social and isolated enhancers and quantified from the peaks with significant binding enrichment the number of peak differences for all histone marks between both enhancer classes. Given that histone repressive tags govern cell-specific transcriptional identity by blocking unspecific gene programs (*63*), we subsequently compared its binding enrichment for both enhancers classes. We quantified the total number of significantly enriched repressive peaks (H3K9me and H3K27me) for social and isolated enhancers in a given module. This step allowed us to prioritize and identify cooperative social enhancers and their transcriptional regulators for a given module. Social enhancers coordinates were used to study binding enrichment analysis with ChIP-atlas dataset. We used TFs and regulators (n=15217 experiments) with binding enrichment defined as fold-change > 2 and p-value <0.05. Once we established the top regulatory binders to territorial social enhancers, we assessed the binding enrichment for proximal regulatory regions (±10kb from TSS) for genes within the submodule. For a given submodule or module, we then selected top ranked transcriptional regulators binding both social enhancers and promoter regions, and generated a hierarchical list of likely active promoter-enhancers links with their intertwined inter-submodular dependencies. This unbiased approach exploited both integrative and topological associations, which allowed us to generate qualitative information for downstream analysis. In this case, we used for each selected module our previously described pipeline for interrogation of interacting transcriptional regulators (see transcriptional dependencies). Finally, for interacting transcriptional regulators territorially enriched, we highlighted those that recapitulated locally our global transcriptional inferences. This was followed by word-cloud representation (based on number of interactions) and network functional enrichment. We complemented this approach by interrogating local topological behavior of metabo-HAR domains as follows: with a targeted manually curated strategy, we used the ucsc genome browser (archived in https://genome.ucsc.edu/) to find the metabo-HAR domain window containing genes and enhancers of interest. We highlighted active elements and assessed by ChIP-regulatory-binding and TFBS motifs, whether members of the PGC1A-interating network were present among the regulators.

##### 3.20. Selection of active non-coding elements within PGEs

To select PGE or fPGE hubs that contained experimentally tested activity of regulatory elements, we used the VISTA enhancer resource containing in-vivo validated enhancer information (archived in https://enhancer.lbl.gov/frnt_page_n.shtml) (*62*). For any given PGE hub (PGEs derived from HAR-associated genes, table S3), we evaluated the number of active non-coding elements within the chromatin region. For example, around 84% of metabo-HAR PGE regions displayed active elements. We next assessed the number of active elements for the fPGEs that reside within chromosomes composing nuclear compartments of interest (see nuclear compartment reconstruction). We then labeled these elements for experimental validation. Using the short-range linking network, we were able to prioritize genes (promoters or proximal regulatory regions) that were associated with active elements. We next overlapped ncHARs (archived in http://docpollard.org/research/) (*21*), with top ranked active regulatory elements within the network module and submodule of interest. We used these coordinates for downstream experimental validation.

##### 3.21. Transcriptional compartment reconstruction and functional embedding

Once we evaluated global relationships for the trans chromatin segments of interest harboring gene clusters from PGE decomposition, we took advantage of the network-based analysis approach to select with high confidence structural representations valuable for the network global dependencies, yet sharing functional annotation. This protocol can be used to study any PGE constructed from any given gene set and it is based on selecting trans-interacting chromatin segments that both display topological qualities and harbor functional-PGEs (fPGE) of similar function (see PGEs in trans-interacting-network). Briefly, top bridging trans-interacting segments harboring fPGEs that shared similar functional annotation to top hub trans-interacting segments were chosen for nuclear compartment reconstruction and further experimental validation. Given that our combinatorial transcriptional based selection and our topological decomposition both showed lipid metabolism as a top functional candidate, highly interacting segments (with top bridging scores) harboring fPGEs for lipid metabolism were chosen for reconstruction. Following any possible interaction in our interacting chromatin network from hESC Hi-C data, we iteratively sought for other segments with high bridging score and with a functional embedding for lipid metabolism. To evaluate if these interactions were present in other Hi-C datasets, we tested their presence in other tissues and cell types (see Hi-C data analysis). Because we were looking to identify regions with high likelihood of interaction for downstream experimental validations, we only highlighted their behavior as binary, either present or absent without any other integrative method. They were displayed as shared contacts across studies. To fully reconstruct highly ranked and functionally embedded structural relationships, we used the top bridging segment, harboring metabolic-HAR hub, Chr1:48.5mb to derive other local homotypic interacting segments, and other interacting heterotypic segments with high bridging coefficient and related function, such as Chr1:62mb, Chr2:170mb, Chr2:118mb, Chr4:42mb, Chr4:1mb and Chr11:34mb. Interestingly, we found a high density of interacting homotypic regions around chr1:48, including chr1:43, chr1:39, displaying top bridging coefficient scores in the network. This highlights this intertwined region of the genome as a modulator of information flow within chromatin architecture. As shown later, this region contains genes related to lipid metabolism, and displays genetic variations related to lipid metabolism as well. We displayed our selection as Chr1-based sub-compartment triplets with Hi-C derived homotypic and heterotypic interactions, as well as the location of PGEs and fPGEs with similar functional annotation within the composing chromosomes. This subnetwork of compartments was used for downstream genomic-range topological genetics and experimental validation in biological systems of interest.

##### 3.22. Modeling of genomic hubs using chromosome 3D-reconstruction

To assess global and local 3D conformations from structural data of chromosomes enriched in PGEs hubs from our previous analysis, we used established protocols (*64*), (*65*, *66*) (archived as GSDB at http://sysbio.rnet.missouri.edu/3dgenome/GSDB/, and Spacewalk at http://3dg.io/spacewalk/). Briefly, for global chromosome reconstruction GSDB and spacewalk modeling was used to explore the architecture and homotypic interacting clusters where PGEs hubs reside. GDSB is a comprehensive repository of 3D modeling algorithms and

Hi-C data. Spacewalk is an application interface with interactive animation of genomic-ranges. Using the GSDB application, we first compare the performance between 2 distance-based algorithms for 3D Hi-C reconstruction. We obtained structure evaluation for representative Hi-C at 40-kb resolution of 3D reconstruction using 3D maximum likelihood algorithm (*64*) and LorDG, which uses a nonlinear Lorentzian objective function to optimize realistic interactions (*67*). In addition to the same Hi-C data previously used for isolating trans-interacting segments (hESC HiC; GSE35156), we used other HiC dataset including colorectal adenocarcinoma cells (GEO: GSE105318), human brain pericytes (GSE105513), astrocytes from the spinal cord (GSE105957), human fibroblast IMR90 (GSE35156), brain microvascular endothelial cells (GSE105544), and human hepatic endothelial sinusoid cells (GSE105988). To compare the structure reconstruction between these distance-based algorithms, we used the average spearman correlation between Expected Distance vs. Reconstructed Distance (*65*) on the chromosomes that composed our selected genome compartments and the small chromosome 19 as a control. Next, we represented the data in a XY plot as performance score (expected vs reconstructed distances) vs chromosome size (log). We evaluated the performance by performing a linear regression followed by F-test statistics. This algorithmic selection showed that 3DMax maximum likelihood algorithms better represent 3D Hi-C data with less bias against long chromosomes. Therefore, we evaluated in spacewalk 3DMax pdb-files to assess the location of metabolic-HAR hubs, homotypic interactions, homo-heterotypic relationships, as well as the genomic range of their cluster association.

#### 3.23. CTCF-bound structural regulatory regions

##### 3.23.A. CTCF-bound regulatory regions

To identify promoters bound by the structuring protein CTCF we used the ChIP-atlas dataset as previously described. For those genes identified as CTCF-bound, we next assessed whether they are classified as social cooperative regulatory regions by obtaining their association degree from the integrative regulatory network (see short-range linking network). This was followed by hypergeometric enrichment and Chi-square test of CTCF-bound promoters within promoters defined as social or cooperative, compared to random gene sets. In parallel, we interrogated ChIPseq-enrichment of CTCF in enhancers defined as isolated or social (see short-range linking network), using again the ChIP-atlas dataset. In addition, we used other databases for interrogating CTCF-bound elements in local chromatin regions in the next section.

##### 3.23.B. Structural enhancers prioritization

We sought to identify structural enhancers within genomic ranges defined by fPGEs hubs (in the present study as metabolic-HAR genomic hubs). We used enhancer coordinates within submodule territories of the integrative short-range linking network. To prune our global network approach with enhancer datasets including structural experiments (from ChIA-PET and Hi-C), and identify TF binding candidates across several cell lines, we used the HACER (human active enhancer to interpret regulatory variants) database (*68*) (archived in http://bioinfo.vanderbilt.edu/AE/HACER/index.html). In brief, HACER is a comprehensive database that incorporates interaction frequency experiments to the linking association derived from GROseq, PROseq and CAGE profiles. Enhancers in HACER are cross-referenced with VISTA enhancers (*62*), ENCODE elements, the ensemble regulatory build (*69*), and chromHMM (*70*). Enhancer interactions in HACER are defined using similar parameters to the GeneHancer regulatory network used here to perform network propagation of short-range associations. These are genome distance, eQTLs, and FANTOM. Besides these, the authors used structural-mediated linking to improve their associations with experimental interactions from 4D genome consortium (*71*) and chromatin contacts derived datasets (*61*, *72*, *73*). HACER has integrated ENCODE ChIP-seq data from several cell lines and tissues. This includes > 700 000 TF-enhancer binding and 156 TFs. In order to interrogate TF-binding of structuring candidates as well as experimental structural validations of enhancer-gene associations, we used as inputs the coordinates of regulatory regions within genomic hubs (mHAR hubs). We then used HACER to classify enhancers bound by CTCF as social or isolated with the network parameters of the short-range linking networks (see short-range linking network). With this approach we estimated the average number of links that structuring enhancers display within selected chromatin regions. In sum, this was a qualitative step to filter short-range network representations within regions of interest, for downstream experimental validation. For the filtered enhancers within genomic hubs, we used HACER GWAS-integration of regulatory regions displaying SNPs for guiding experimental validations (*74*).

#### 3.24. Cell-specific regulatory regions

To identify cell-specific regulatory elements in our global short-range association network, we used 2 different approaches relying on previously established protocols and datasets. We used the enhancer atlas (archived in http://www.enhanceratlas.org/index.php)(*75*). Briefly, this atlas contains regulatory elements for > 500 tissue and cell types across several species. It incorporates high-throughput experiments including chromatin states, DNAse-seq, ATAC-seq, ChIA-PET, GRO-seq, STARR-seq and MPRA. To combine different tracks from each dataset and generate a consensus assembly the authors used an unsupervised approach that weights each track for the global track (*76*). Peak filtering is done at 2500 bp and merging with the average summit in the size of the average peak width (ASW) (*77*). To focus on the degree of overlap between regulatory elements, Tianshun et al. used the Jaccard index with intersection over union (*75*). Using this resource we downloaded cell-specific enhancer coordinates and mapped them with our short-range regulatory network, highlighting their location and network statistics. We then used this qualitative filtering step for experimental validation of cell-specific activity during fasting. Other tissue-specific modules with motif enrichments were obtained from Epimap (*78*).

##### 3.25. Genetic enrichments in genomic hubs using integrative genomic-range genetics

To find metabolic disease associations for a given genomic-range, we used the curated common metabolic disease portal. (archived in https://hugeamp.org/). To evaluate genetic association enrichments for fPGE hubs or genomic regions of interest, we used previously established methods (currently used at hugeamp) (*79*). Briefly, this protocol is based on meta-analysis of associations to provide an estimate of variant effect by integrating information from diverse studies. By taking into account the sample overlap among datasets, this approach weights in the study contribution for the final estimate. The meta-analyses are implemented in METAL which is a tool for genome-wide scans that was described by Willer et al. (*79*). Their approach is based on p-values and subsequent z-statistics for the impact of the association. Given that all studies used are aligned to the same allele, z and p-value are obtained from the weighted sum of individual statistics. The weights are then the square-root of the number of individuals in each sample. Following their streamline implementation of METAL and genome-wide scans we interrogated the cumulative association of genomic-regions to metabolic disease traits. For this, we used fPGE hubs in chromatin segments composing heterotypic compartments including those in Chr1, Chr2, Chr4, Chr8 and Chr11. Next, we assessed genetic associations to interacting chromatin segments and displayed p-value of association for a given region. We obtained trait groups and top traits within groups. We displayed the integrative results for all genomic-range traits from metabo-HAR regions as a wordcloud representing the number of appearances of groups and trait descriptions.

##### 3.26. Dynamic gene expression and Monte Carlo transcriptional models

We used dynamic gene expression variation between challenged (i.e. fasting) and control states to quantify and model intrinsic gene expression fluctuations captured by targeted gene expression (see analysis of gene expression). Intrinsic variations in dynamic gene expression were then used to build nonlinear statistical models followed by a Monte Carlo approach (see Monte Carlo modeling). Modifications in cell culture for dynamic capture of gene expression were as follows: cells were plated in 96 well plates in equal cell numbers (1x10e4 cells/well). Induction and differentiation were carried out as previously described (see mammalian cell culture). Once differentiated, 0% glucose was induced in half of the wells followed by dynamic time tracking, where every plate was used to track 1 different time point. After 3 hours of fasting, cells were collected (at each time point) and RNA extraction protocol was performed every 10 minutes for 6-10 different times (T0 + 10 additional measurements; where T0 = 3 hours after fasting induction). We collected between 4- 6 biological replicates per time point followed by qPCR gene expression analysis as previously described. To compare the expression of genes within the same TADs, CCDs or submodules as previously defined (see structural chromatin relationships and short-range link networks), we selected TADs within PGE hubs of interest, and quantified fold-change distribution across time variations. We used genes with social or isolated promoters defined by network modeling features as described before (defined by network integration and propagation analysis of short-range link networks). In brief, we obtained fold-change variations (and distributions) by comparing control and glucose fasting in every time point evaluated. These dynamic fold-change fluctuations of fasting-induced genes were used to build downstream statistical models. We were interested in studying, for differentially expressed genes within the same TADs or hubs, their gene expression bursting fluctuations over time. We observed that, for genes regulated during fasting, there were time variations in fold-change over control, indicative of specific bursting variation (**fig. S16**). Genes that were defined as isolated by our integrative network approach did not display fluctuations as those defined as having social cooperative promoters (**fig. S16**). Genes with non cooperative promoters displayed fold-change variations over time that were stable and constant during glucose fasting (**fig. S16**). On the other hand, active genes with social promoters exhibited higher bursting fluctuations in several experimental setups indicative of different kinetics of expression. To control for primer effect we tested several sets and obtained consistent results. In addition, we observed that perturbing genomic plasticity during glucose fasting by environmental variations affected the fluctuation in social genes.

###### 3.26.A. Non-linear sine-wave models on dynamic gene expression data

In order to capture these variations and model statistical differences, we investigated non-linear models. Experimental observations showed an oscillating pattern of expression for some genes within hub domains. Their behavior was reminiscent of sinusoidal spectral patterns where fluctuations seem to capture discrepancies at different frequencies. This oscillatory dynamic behavior has been observed in complex biological systems where synchronicity of population behavior has been exploited to extrapolate principles for non-stationary variability (*80*). Moreover, gene expression burst kinetics have been shown to differ between enhancer- and promoter-encoding burstings as well as models for condensate-control of transcription (*81*). With this in mind, we used non-linear sinusoidal regression (sine-wave with non zero baseline model in graph pad prism) to model our experimental observations. We fitted models for each target gene to assess their best-fit parameter. Non-linear least squares model was used to fit the regression without weighting the data points (*82*). To discriminate the best-fit parameters between models we used the Akaike's information criteria (AICc), which assumes non-nested inferences. Goodness-of-fit was evaluated by $R^2$ and standard error of the estimate. Parameters obtained from the sine-wave models are as follows: amplitude defines the height of the waves from the baseline; frequency is the number of cycles per time unit; baseline is the Y value where the curve oscillates. We next compared the best-fit parameters from biological experiments with randomly generated values that preserved amplitude distributions. To this end, we generated random values per time point followed by sinusoidal non linear regression, and used AICc to evaluate goodness-of-fit between them. In addition, we also modeled basal expression detection of house-keeping and unstimulated genes. This showed discrepancies between the models, with amplitude and frequencies displaying the largest variation. Burst profile models of social cooperative genes exhibited higher frequencies, amplitude and baseline parameters. We therefore used these quantified parameters to simulate in large scale sinewave behavior constrained by these model variations. We used an in-built simulation option in graph pad where XY table values were selected. Time-series variations were on the X values with similar intervals as our experimental data set (>1000 interval simulations). Y values were generated using sinewave non zero baseline model equations with amplitude, frequency, phaseshift and baseline constants from experimentally derived models. Gaussian random error was used to generate random scatter values. Once the simulations were obtained we performed nonlinear sinewave regression on simulated values to get a better estimation of confidence intervals and estimate best-of-fit and oscillatory parameters. We used these statistical approximations for Monte-Carlo simulation followed by outcome discrimination of the models recreated.

###### 3.26.B. Monte Carlo modeling of dynamic gene expression data

To estimate the distribution and range of variation among oscillatory parameters in models from different experimental biological observations, we used a Monte-Carlo approach on large-scale simulated data from our previously described steps (see non-linear sinusoidal models). From the simulated sinusoidal models, we obtained best-fit estimates of models constrained by experimental observations. Using these best-fit estimates, we modeled >1000 paralleled models using Monte-Carlo (graph pad prism in-built monte carlo implementation). We defined sinewave equation parameters for each group followed by nonlinear sinewave regression. Outcome discrimination on each model was carried out with AICc and standard error of the estimate from experimental groups and random generated models. We next tested distribution discrepancies for oscillatory parameters (amplitude, frequency, and baseline) from Monte-Carlo models. Mean distributions between biological groups from which the models were obtained, were used to make statistical comparisons between model fluctuations and estimate p-value for model assumptions.

##### 3.27. Mice-human chromosomal synteny

To evaluate genomic conservation regions between humans and mice, positional gene enrichments of mice orthologue metabolic-HAR genes were obtained following the same criteria as indicated before (see functional embedding of positional gene enrichments). We quantified the number of optimal-PGE hubs (> 5 genes) between human and mice positional enrichments and compared them to the number of PGE hubs derived from random permutations. Subsequently, mice PGEs regions of < 15-Mb were selected for downstream analysis. We next evaluated whether metabolic-HAR regions (mPGEs and hPGEs) were located in syntenic domains, using the synteny portal (archived in http://bioinfo.konkuk.ac.kr/synteny_portal/) (*83*). This revealed that 82% of PGE hubs were in syntenic regions and 18% were located in split regions in mice. We also displayed the level of genomic conservation in breakpoint regions and in syntenic blocks (*56*).

##### 3.28. Murine gene knockout phenotype networks

For a given gene set (e.g., metabolic-HAR genes), we identify gene and phenotype relationships from data stored in the Jackson laboratory repository containing gene perturbation phenotyping. This dataset has been parsed and is stored in ENRICHR as the mammalian perturbation catalog (*24*). Network reconstruction was done with all phenotypes associated with gene knockout followed by network modeling and analysis. In brief, a network was built with nodes defined as gene knockout names connected by edges to each phenotype associated. To identify phenotype modules we used network modularity and to identify gene hubs per module we used node degree, pagerank and eigencentrality. To identify phenotype hubs we used node prioritization (using node degree, pagerank and eigencentrality), followed by quantification of phenotype representations by proportion frequencies (top 30 phenotypes). Intermodule communication was performed by quantifying interactions between modules. We performed positional gene enrichment for gene hubs from each module and from interacting modules. Proportion of colocalized positional gene enrichment was defined by comparing positional clusters (with > 5 genes per region) from genes in each module and combined genes from associated modules (see identification and functional annotations of genomic hubs by positional gene enrichments).

##### 3.29. Data and Code Availability

Analyses were performed in R (3.5 and 3.6) and python (2.7, 3.5 and 3.7). Libraries for data analysis include numpy, pandas, scikit-learn, scanpy and scipy. The datasets reported in this paper are available in supplementary materials and can be found at https://github.com/leandroagudelo189/Metabolic-resilience-topo-genomics. Below is detailed data and code source to different datasets used in this paper (Table S**12**). When indicated, network analysis and visualization were done using the python library NetworkX, and the software GEPHI. Functions to evaluate networks have been created and validated by the Ideker Lab (archived in https://github.com/idekerlab/Network_Evaluation_Tools) and are stored on the network data exchange (NDEx)(*84*). Other functions to evaluate network topology can be found as inbuilt analysis tools in GEPHI. Tissue-specific networks for functional annotation of topological dependencies can be found at http://giant.princeton.edu and https://hb.flatironinstitute.org/. Positional gene enrichment analysis tool can be found at http://silico.biotoul.fr/pge/. Functional annotation tool from a large library of biological repositories can be found at https://maayanlab.cloud/Enrichr/. To filter out PGE regions without cytogenetic band enrichment, we used a curated list of cytogenetic band-gene dependencies. Prediction of TFBS from DNA sequences can be found at http://alggen.lsi.upc.es/. Prediction of conserved TFBS in promoter regions of genes of interest can be found at http://www.gsea-msigdb.org/gsea/msigdb/index.jsp. Tissue-naïve and tissue-specific protein-protein interactions used for n+1 network reconstruction can be found at http://www.interactome-atlas.org/. Tissue-naïve protein-protein interactions used for full network reconstruction can be found at https://thebiogrid.org/. In silico prediction of intrinsically disordered regions in amino acid sequences can be found at http://www.pondr.com/, https://iupred3.elte.hu/, https://mobidb.bio.unipd.it/. Curated experimentally validated phase separation database stored at http://db.phasep.pro/. Context-dependent phase separation score from ensemble parameters of conformation changes present in disordered proteins. They offered discrepancies of conformations that are related to the local environment such as salt, solvation, multi-molecular interactions and ligand presence. These can be found at http://pappulab.wustl.edu/CIDER/about/ and can be implemented locally following instructions stored at http://pappulab.github.io/localCIDER/. Phylogenetic analysis tool information for protein residues conservation score and 3D modelling of residue exposure can be found at https://consurf.tau.ac.il/. Evolutionary domain conservation of protein domains with known structure can be found at https://consurfdb.tau.ac.il/. ChIP-atlas methodology, documentation and functions for analysis can be found at https://github.com/inutano/chip-atlas. Compendium of human chromatin contact maps has been created by the Ren Lab and is stored at GEO: GSE87112 (schmitt, bing ren 2016). Cis-regulatory elements relationship were obtained from GeneHancer, and can be found at http://www.genecards.org/. VISTA enhancer elements can be found at https://enhancer.lbl.gov/frnt_page_n.shtm. HACER, human active enhancer to interpret regulatory variants database is archived in http://bioinfo.vanderbilt.edu/AE/HACER/index.html. A curated database of cell-specific active regulatory regions can be found at http://www.enhanceratlas.org/index.php. Chromatin 3D reconstruction tool and code can be found as follows: GSDB at http://sysbio.rnet.missouri.edu/3dgenome/GSDB/; and Spacewalk at http://3dg.io/spacewalk/. CTCF-binding site prediction tool can be found at https://insulatordb.uthsc.edu/. To isolate metabolic disease associations for a given genomic-range, we used the common metabolic disease portal, archived in https://hugeamp.org/. For topological comparative genomics and to evaluate chromosomal synteny, tools can be found at the synteny portal, archived in http://bioinfo.konkuk.ac.kr/synteny_portal/.

#### 4. Experimental materials and methods: functional topo-genomics framework

##### 4.1. Animal studies

All animal experiments were performed in accordance with internationally accepted guidelines and principles for the use of laboratory animals, and were approved by the Novo Nordisk Research Center Seattle Institutional Animal Care and Use Committee, Novo Nordisk research center Lexington Massachusetts, the Novo Nordisk Ethical Review

Committee, Novo Nordisk Research Center in China, and regional ethics committee of Alicante, and Universidad Miguel Hernández-CSIC Spain. Adult Wild-type male C57BL/6J mice aged 6-12 weeks were obtained from Jackson Laboratories (Stock # 000664). Adipose tissue Pgc1a knockout mice were generated as previously described (*85*). Pgc1a loxp mice (jackson Laboratories Strain #:009666) were bred with adiponectin CRE mice (jackson laboratories, strain #:010803). To generate adipose tissue Ctcf+/- heterozygous mice, Ctcf lo.xp mice were generated as previously described (*86*). Ctcf tm1a(EUCOMM)Wtsi mice were bred with ACT-FLPe mice (jackson laboratory, USA), followed by breeding of Ctcf loxp mice with adiponectin CRE mice. To generate adipose tissue Yy1+/- heterozygous mice, floxed Yy1 mice (jackson laboratories, strain #:014649) were bred with adiponectin CRE mice. Whole body Pgc1a knockout mice were used (jackson laboratories, stain #008597). Floxed Dgat1 mice (jackson laboratories) were bred with adiponectin CRE mice. Ncoa7 knockout mice were generated as previously described (*87*) (*87*) and are part of the knockout mouse project (MMRRC_046720-UCD, C57BL/6N-Atp12atm1(KOMP)Vlcg/MbpMmucd). To generate mice with double deletions, whole body (wb) Ncoa7 KO mice were crossbred with fat-specific mutant mice (fat-Pgc1a-/- or fat-Dgat1-/-), founders were backcrossed until fat-specific mutant and wb-Ncoa7+/- were available. For experiments, both male and female mice of similar age were included. Mice were housed 4-8 per cage with ad libitum access to water and food (Type IV Scanbur Cages; 1800 $cm^2$ floor area). Animal holding rooms temperature ranged from 20-26 °C and humidity ranged from 30-70%. Lighting was on an automated light/dark cycle with lights on between 6:00 AM and 6:00 PM. Mice were provided with Alpha Dri bedding material to just cover the bottoms of the cage, and one Nestlet square. Environmental enrichment included a plastic hut, a paper hut was provided. Mice received a hydrogel cup upon arrival until familiarized with the automatic water system. Standard rodent chow diet (PicoLab Rodent Diet 20 Lab Diet #5R53) was provided to the mice upon arrival until the diet was switched from chow to high fat (described below). All food and water were available ad libitum. Mice were acclimated to housing for one week.

##### 4.2. High fat diet

6-12 weeks mutant mice and control mice were placed on a 60% high fat diet (High-Fat Diet #D12492 (20.0% Protein, 60% Fat, 20% Carbohydrate) during 6 weeks and they reached dietary induced obese status (DIO). After this point, mice were moved from group housing to 2 mice per Type IV Scanbur cage, separated by an opaque cage divider. Bedding and environmental enrichment was as described above. Food and water were provided ad lib regardless of housing condition. Similarly, 6-12 weeks control, and fat-specific mutant mice and mice with double deletion (as previously described) received a high-fat diet for 6 weeks. Control and mutant mice undergoing HFD of 6 weeks were concomitantly subjected to an intermittent fasting protocol (6 weeks), consisting of 2 non-consecutive days per week (Tuesdays and Fridays) without food access (24 hours each day).

##### 4.3. Mice treatments

wild-type, mutant mice, and mice undergoing 1 week of HFD were used for fasting challenges of 6, 12 and 24h (when indicated), after which tissue extraction was performed. Bafilomycin A1 (1 mg/kg, Sigma-Aldrich) was injected intraperitoneally to inhibit autophagy during fasting challenges. DGAT1 inhibitor by intraperitoneally injection (2 mg/kg Sigma Aldrich, CatNo:A922500). To correct pH, sodium bicarbonate ($NaHCO_3$, 20mg/kg, Sigma-Aldrich) and glucose solution (1 mg/kg) were injected intraperitoneally 1 hour prior tissue collection. This was followed by tissue extraction and further processing for molecular experiments.

##### 4.4. Mammalian cell culture and treatments

Human embryonic kidney 293 cells (HEK293) were cultured on glass-bottomed MatTek dishes. Cells were grown in Dulbecco's modified Eagle's medium (Gibco, Thermo Fisher Scientific), 10% fetal bovine serum (Thermo Fisher Scientific) and 10U/mL penicillin-streptomycin (Thermo Fisher Scientific) and incubated at 37ºC and 5% $CO_2$ in humidified incubator. Human primary pre-adipocytes (ATCC) were differentiated into mature adipocytes and used for glucose fasting, chemical treatments and genome editing experiments. Murine inguinal adipocytes: SVF from inguinal fat depots of 6-12-week-old C57BL/6J wild-type and Pgc1a-/- mice were isolated and prepared as previously described (*88*). The differentiation cocktail was used during the first 2 days of culture, followed by rosiglitazone and insulin for 6 to 8 days. Murine brown adipocytes: Primary brown adipocytes from male C57BL/6J wild-type and Pgc1-/- mice were isolated and cultured as previously described (*89*, *90*). Murine hepatocytes: Primary hepatocytes from male C57BL/6J wild-type and Pgc1-/- mice were isolated and cultured as previously described (*91*). Cell treatments: Indicated cell cultures were kept in different condition media: Dulbecco's without glucose for fasting condition or Dulbecco´s high glucose medium for control. Adipocytes under differentiation were transfected using lipofectamine 3000 (Invitrogen, Thermo fisher technologies) with siRNA for YY1, PGC1A, CTCF, EP300, ATG4C, DGAT1, INSIG2, CAT,

ANGPTL3, PCSK9, NCOA7, and scramble control (Applied biosystems). Once they were differentiated, they were processed for analysis of gene expression, or follow up experiments (as indicated). HEK293 cells were transfected as previously indicated with siRNAs or scramble control 24h previous to downstream experiments. DGAT1 specific inhibitors were used at 1uM (Sigma Aldrich, CatNo:A922500). C:16-acylcarnitine was used as representative long-chain acylcarnitines for cellular experiments at different concentrations as indicated (Sigma Aldrich 61251 - Palmitoyl-L-carnitine). Etomoxir (Sigma Aldrich, 50uM) was used to block CPT1A-B mitochondrial fatty acid import channel as indicated. 1,6 Hexanediol at 2.5mM (240117, sigma aldrich), oligomycin (1uM; complex V inhibitor), bafilomycin A1 (1uM; lysosome vATPase-inhibitor), Sodium bicarbonate (Sigma aldrich, 20mM) and DMEM media with and without glucose (Sigma aldrich). Plasmids and transfection: GFP, Cas9 with sgRNAs, homology vector, and pMACs 4.1 plasmids, wild-type and mutant Pgc1a plasmid were transfected to adipose progenitor cells or primary adipose progenitors (during differentiation) using Amaxa-nucleofector (Lonza) and nucleofector kits for transfection of primary cell lines following manufacturer's protocols (Lonza). HEK293 Cells were transfected when they reached 70% confluence with plasmid DNA GFP-PGC1A (Addgene, Massachusetts, USA) using Lipofectamine 3000 (Invitrogen, Thermo Fisher Scientific) and removed 24 h post-transfection, according to standard manufacturer protocols. Transfected cells were imaged 24-48 h post-transfection.

##### 4.5.Analysis of Gene Expression

Total RNA was isolated from cells or tissues using Isol-RNA Lysis Reagent (5 PRIME) according to manufacturer's instructions. Amplification Grade DNase I (Life Technologies) was used to treat 1 µg of RNA, from which, 500 ng were used for cDNA preparation following Applied Biosystem Reverse Transcription Kit (Life Technologies). Quantitative Real-Time PCR was performed in a ViiA 7 and QuantStudio Real-Time PCR system thermal cycler with SYBR Green PCR Master Mix (both Applied Biosystems). Analysis of gene expression was performed using the ΔΔCt method and relative gene expression was normalized to hypoxanthine phosphoribosyltransferase (HPRT) mRNA levels. Gene expression analyses were expressed as mRNA levels relative to controls. When indicated gene expression analyses were shown as mRNA levels relative to controls. To evaluate transcriptional compartment activity of genes within genomic hubs, mean fold-change variation over control for each gene was used to compare overall effects (displayed as a module) for different biological conditions. Primer sequences available upon reasonable request.

##### 4.6. Chromatin immunoprecipitation

ChIP experiments were performed as previously described (*92*) with the following modifications: Cells were plated and (2-10 million) pooled before homogenization. Cells were homogenized in PBS and crosslinked with 1% formaldehyde during 10min. For tissue, 50mg were homogenized in PBS and crosslinked with 1% formaldehyde for 10 min. Glycine (125 mM) was added, followed by centrifugation at 4000 rpm for 5 min. After aspirating the supernatant, the pellet was washed with 1 ml cold PBS followed by centrifugation at 4000 rpm for 5 min. This step was repeated with buffer 1 (0.25% Triton X-100, 10mM EDTA, 0.5mM EGTA, 10mM HEPES, pH 6.5) and buffer 2 (200mM NaCl, 1mM EDTA, 0.5mM EGTA, 10mM HEPES, pH 6.5). The pellet was then resuspended in 300 µl lysis buffer (1% SDS, 10mM EDTA, 50mM Tris-HCl, 1X protease cocktail inhibitor, pH 8). Samples were sonicated on ice using a Bioruptor (Diagenode) 30 times for 30 seconds ON and OFF intervals. After the lysates were sonicated to shear the DNA to fragment lengths of 300bp, the complexes were immunoprecipitated with antibodies specific for CTCF (Abcam, Anti-CTCF antibody [EPR7314(B)], cat# ab128873), YY1 (Abcam, anti-YY1, cat# ab245364), PGC1A (sigma-aldrich calbiochem, cat# ST1204), EP300 (Abcam, Anti-KAT3B / p300 antibody, cat# ab10485), and H3K27Ac (Abcam, Anti-Histone H3 (acetyl K27) antibody, Cat# ab4729), H3K4me3 (Anti-Histone H3 (tri methyl K4) antibody Cat#ab12209), STAT1 (Thermo scientific, #AHO0832). 10-20 ul of antibody was used for each immunoprecipitation. No antibody controls were also included for each ChIP assay and no precipitation was observed by quantitative Real-Time PCR (qPCR) analysis. Input samples were processed in parallel. The antibody/protein complexes were collected by either salmon sperm DNA/protein A agarose slurry or Protein A/G PLUS agarose beads (Santa Cruz sc-2003) and washed several times. The immuno complexes were eluted with 1% SDS and 0.1 M $NaHCO_3$ and samples were treated with proteinase K for 1 hour and DNA was purified by phenol/chloroform extraction, ethanol precipitation and resuspended in 20 ul $H_2O$. Input DNA and immunoprecipitated DNA were analyzed by quantitative PCR. Primer sequences used to detect each DNA region are available upon reasonable request. .

##### 4.7. Chromosome conformation assays

###### 4.7.A. ChIP-loop immunoprecipitation of chromatin interactions:

ChIP-loop-qPCR protocol is based on ChIP immunoprecipitation assay previously described with the following modifications: In brief, 1 to 15 million crosslinked cells were resuspended in lysis buffer ( 60mM Tris pH 7.5 0.5% Igepal, 0.25% Sodium-deoxycholate 0.1% SDS, 150mM NaCl, and protease inhibitors) and rotated at 4° C for 30 minutes. Nuclei were pelleted at 4° C for 5 minutes at 2500 rcf. Pelleted nuclei were washed once with 500 µL of ice-cold lysis buffer. Supernatant was removed again and the pellet was resuspended in 100 µL of 0.4% SDS and incubated at 62° C for 30 minutes. To quench the SDS, 10% Triton X-100 were added and samples were rotated at 37° C for 15 minutes. NEB Buffer 2 and 15 µL of 25 U/µL HindIII (NEB; following manufacturer's recommendation were then added and samples were rotated at 37° C for 2 hours (15 µL was used for 10–15 million cells, 8 µL for 5 million cells, and 4 µL for 1 million cells). The enzyme was then heat-inactivated at 62° C for 20 minutes. 52ul incorporation master mix was then added: 37.5 µL of 0.4 mM biotin-dATP (Thermo Fisher 19524016), 4.5 µL of a dCTP, dGTP, and dTTP mix at 10 mM each, and 10 µL of 5 U/µL DNA Polymerase I, Large (Klenow) Fragment (NEB, M0210). The reactions were rotated at 37° C for 1 hour. 948 µL of ligation master mix was then added: 150 µL of 10X NEB T4 DNA ligase buffer with 10 mM ATP (NEB, B0202), 125 µL of 10% Triton X-100, 3 µL of 50 mg/mL BSA (Thermo Fisher AM2616), 10 µL of 400 U/µL T4 DNA Ligase (NEB, M0202), and 660 µL of water. The reactions were then rotated at room temperature for 4 hours. After proximity ligation, the nuclei with in-situ generated contacts were pelleted at 2500 rcf for 5 minutes at room temperature. As described above, the pellet was then resuspended in 300 µl lysis buffer (1% SDS, 10mM EDTA, 50mM Tris-HCl, 1X protease cocktail inhibitor, pH 8). Samples were sonicated on ice using a Bioruptor (Diagenode) 20 times for 15 seconds ON and OFF intervals. Clarify samples for 15 minutes at 16100 rcf at 4 degrees and w2X volume of ChIP Dilution Buffer (0.01% SDS, 1.1% Triton X-100, 1.2 mM EDTA, 16.7 mM Tris-HCl pH 7.5, 167 mM NaCl) was added. The complexes were immunoprecipitated with 10-20 ul of specific antibodies (for each immunoprecipitation). 60 µL of Protein A beads (Thermo Fisher) were precleared. Beads were washed and resuspended in ChIP Dilution Buffer to a volume of 50 µL per tube (100 µL per sample), then added to samples and rotated at 4° C for 1 hour. Samples were placed on a magnet and supernatants were transferred into fresh tubes. Beads were washed three times each with Low Salt Wash Buffer (0.1% SDS, 1% Triton X-100, 2 mM EDTA, 20 mM Tris-HCl pH 7.5, 150 mM NaCl), High Salt Wash Buffer (0.1% SDS, 1% Triton X-100, 2 mM EDTA, 20 mM Tris-HCl pH 7.5, 500 mM NaCl), and fresh LiCl Wash Buffer (10 mM Tris-HCl pH 7.5, 250 mM LiCl, 1% NP-40, 1% sodium deoxycholate, 1 mM EDTA). The antibody/protein complexes were collected by Protein A/G PLUS agarose beads (Santa Cruz) and washed several times. The immuno complexes were eluted with 1% SDS and 0.1 M $NaHCO_3$ and samples were treated with proteinase K for 1 hour and DNA was purified by phenol/chloroform extraction, ethanol precipitation and resuspended in purified H2O. After reverse cross-linked, ligated DNA was purified and sheared to a length of roughly 250-500 base pairs, at which point ligation junctions were pulled down with streptavidin beads and prepped for targeted quantitative measurement of interactions using qPCR assays comparing different samples. Ligation efficiencies were determined using qPCR assays comparing control templates. Specific primers for computationally selected genomic regions were designed to validate chromatin-chromatin interactions. To evaluate transcriptional compartment activity in the form of chromatin contacts between genomic hubs, mean fold-change variation over control for each chromatin contact was used to compare overall effects (displayed as a group) for different biological conditions. Positive interactions were matched by ChIP-binding of the regulator. Primer sequences available upon reasonable request.

##### 4.7.B. In situ 3C chromosome conformation assays

Depending on isolated cell type (see cell isolation and sorting) between 15 thousand and 2 million cells were used for in situ chromosome conformation assays as previously described (*50*, *93*) with some modifications. Briefly, cells were crosslinked with 1% formaldehyde for 10-15 minutes at room temperature. Cells were lysed with 60mM Tris pH 7.5 0.5% Igepal, 0.25% Sodium-deoxycholate 0.1% SDS, 150mM NaCl, and protease inhibitors. Pelleted nuclei were resuspended in 0.4% SDS and incubated 30 min 60°C for permeabilization. After quenching with 1% Triton-X for 12 min at 37°C, nuclei were digested with 5U/µl DpnII in 1x DpnII buffer overnight at 37°C. Cells were then pelleted (2500g 10min), and the restriction enzyme was inactivated at 65°C for 20 min. For the 1.5hr fill-in at 37°C, biotinylated dATP was used for ligation at 25°C for 4 hours with rotation. Nuclei were pelleted and sonicated in 10mM Tris pH 7.5, 1mM EDTA, 0.25% SDS for 16 min, 100 intensity, 200 cycles per burst, and max temperature of 5°C. DNA was reverse cross-linked overnight at 65°C with proteinase K and RNAse A. Each experiment was performed in biological replicates. After reverse cross-linked, ligated DNA was purified and sheared to a length of roughly 250-500 base pairs, at which point ligation junctions were pulled down with streptavidin beads and prepped for targeted quantitative measurement of interactions using qPCR assays comparing different samples. Data was represented as the fold-change of interaction over control (e.g. fasting/control), where statistically significance and fold-variations were shown as indicated in figure legends. To evaluate transcriptional compartment activity in the form of chromatin contacts between genomic hubs, mean fold-change variation over control for each chromatin contact was used to compare overall effects (displayed as a module) for different biological conditions.

#### 4.8. Commercial assays

Triglyceride (Abcam ab65336), Free Fatty Acid (ab65341), ROS Detection (ab139476), malondialdehyde lipid peroxidation assay (sigma MAK085), Methylated DNA MeDiP ChIP kit (ab117135), mitochondrial isolation kit from tissues (ab110168), mitochondrial complex I activity kit (ab287847), lactate assay kit (ab65331), lysosome activity assay from suspended adipocytes (ab234622), ATP assay kit (ab83355), protein carbonyl content assay kit (ab235631), Oligonucleotide conjugation kit for proximity ligation assays (ab218260) and qPCR on conjugated products following manufacturer's instructions. Plasmid GFP-PGC1 (Addgene, Massachusetts, USA) were mutated at multiple amino acid residues in intrinsically disordered regions following analysis related to phylogenetic residue conservation and in silico condensate features related to charge (see sections in silico phase separation scores, CIDER: identification of context-dependent disordered regions, and consurf package: phylogenetic amino acid conservation and structural features). To perform mutation of selected negative residues for positive residues QuikChange Multi Site-Directed Mutagenesis Kit (Agilent cat#200513) was performed following manufacturer's instructions. TNF alpha ELISA kit (Legend Max mouse kit biolegend, Cat# 430907). MCP1/CCL2 ELISA kit (Legend Max mouse kit biolegend, Cat# 432707). All protocols were performed following manufacturer's instructions.

#### 4.9. Proximity ligation assays

Cell or tissues lysates were obtained using RIPA buffer (50 mM tris pH 7.5, 150 mM NaCl, 1 mM EDTA, 1% (w/v) Triton-X-100, 0.5% (w/v) Na-deoxycholate, 0.1% (w/v) sodium dodecyl sulfate, 20 mM glycerol-2-phosphate, 5 mM sodium pyrophosphate) freshly supplemented with 1 mM DTT and 0.5 mM PMSF. Antibodies were conjugated with oligonucleotides using the oligonucleotide conjugation kit for proximity ligation assays (PLAs, ab218260). Protein concentration was determined by Bradford Protein Assay following manufacturer's instructions (BioRad). Protein extracts and conjugated antibodies were used for PLA in which qPCR on conjugated products was performed following manufacturer's instructions. Proteins of interest (e.g., PGC1A, YY1, and CTCF) were conjugated with 3' or 5' oligonucleotide probes each, followed by immuno-qPCR according to manufacturer's instructions.

#### 4.10. Antibodies

Anti-PGC1A (sigma-aldrich calbiochem, catalog No. ST1204), Anti-PGC1 alpha antibody N-terminal (abcam, ab191838), CTCF (Abcam, Anti-CTCF antibody [EPR7314(B)], cat# ab128873), YY1 (Abcam, anti-YY1, cat# ab245364), EP300 (Abcam, Anti-KAT3B / p300 antibody, cat# ab10485), and H3K27Ac (Abcam, Anti-Histone H3 (acetyl K27) antibody, Cat# ab4729), H3K4me3 (Anti-Histone H3 (tri methyl K4) antibody Cat#ab12209), NCOA7 (Thermo scientific, PA5-26543), ATP6V1B1 (Thermo scientific, PA5-56878), STAT1 (Thermo scientific, #AHO0832).

#### 4.11. Cell isolation and sorting

##### 4.11.A. Adipocyte progenitors from adipose tissue

To isolate adipose progenitors from brown and inguinal adipose tissues, adipose tissue dissociation kit (Miltenyi Biotec) and adipose tissue progenitor isolation kit (Magnetic-activated cell sorting, Miltenyi Biotec) were used following manufacturer's instructions.

##### 4.11.B. Hepatocytes

After the liver is perfused with EMEM at 5 ml/minute for 10 minutes, and then digested with DMEM buffer (containing 0.05% (w/v) collagenase) for 10 minutes, the liver is agitated in a rotary shaker for 10-15 minutes and further dissociated in DMEM medium containing 10µg/ml DNaseI. Parenchyma hepatocytes were then separated from non-parenchymal cells by low-speed centrifugation at gradually increasing rates (30×g, 40×g, and 50×g, for 10 min). The cell pellets were resuspended in perfusion solution, whereas The supernatant fraction containing non-parenchymal cells is used for immune fraction isolation as described below. Hepatocyte cell fraction is resuspended in DMEM medium, viability is checked by trypan blue exclusion, and is used for downstream applications.

##### 4.11.C. Cd11b+ monocyte/macrophage cells from adipose and liver tissue

After adipose tissue dissociation, samples are processed as described below following magnetic-activated cell sorting strategies. Monocytes/macrophages are isolated by the digestion of livers with pronase and collagenase digestion followed by discontinuous gradient ultracentrifugation as described above. After initial centrifugation at 600 rpm, the supernatant cell fraction (containing non-parenchymal cells including immune fraction) is further centrifuged at

150rpm for 5 minutes, and washed with DMEM medium. After washing and centrifugation, the pellet cell fraction is suspended in 90 µl MACS buffer (1xPBS, 0.5% BSA) containing 0.6% citrate-dextrose solution (Sigma). Then, 10 µl of CD11b MicroBeads (Miltenyi Biotec, #130-049-601) are added to the cell suspension and incubated for 20 minutes at 4 degree celsius. During incubation, place a MACS LS column (Miltenyi Biotec, #130-042-401) on MACS MultiStand and wash the column with 3 ml PBS twice and then with 3 ml MACS buffer. The cell suspension is passed to the column to remove CD11b-negative cells. 5 ml of MACS buffer is applied onto the column to collect CD11b+ cell fraction.

##### 4.12. Protein sample preparation and turbidity assays

Protein aliquots were thawed above the upper critical solution temperature to ensure the protein was soluble. Phase separation was induced by mixing (stored in 500 mM NaCl and 20 mM Tris, pH 7.5) with no-salt buffer (0 mM NaCl and 20 mM Tris, pH 7.5) to obtain a solution containing 150 mM NaCl and 20 mM Tris, pH 7.5. Protein concentrations were adjusted as necessary by diluting further with the same buffer conditions. Crowding was induced with different concentrations of PEG 8000 (Sigma Aldrich). Proteins were dissolved in potassium phosphate buffer (pH 7) to a concentration of 160 µM/800 µM as stock and diluted with potassium phosphate buffer (pH 7) in the assay. Recombinant PGC1A and RARA proteins (MyBioSource) were mixed with 0, and 20% PEG 8000 solution with a volume ratio of 1:1 at room temperature to obtain various concentrations. Arginine (R) was dissolved in a potassium phosphate buffer (pH 7) to a concentration of 2, 1.5, and 2 M as stock, respectively. PGC1A, CTCF and YY1 combined or individual solutions were mixed with amino acid solution first to obtain various concentrations of R. Then, 20% PEG 8000 was added to the mixture to a volume ratio of 1:1. Recombinant PGC1A, CTCF and YY1 (MyBioSource) combined or individual solutions were desalted into a potassium phosphate buffer of pH 5, pH 6.2, pH 7.5, or pH 9. Protein solutions were then concentrated into 320 µM/800 µM as stock and diluted with corresponding buffers in the assay. Then, protein solutions were mixed with 20% PEG 8000 solution in a volume ratio of 1:1 at room temperature to obtain various concentrations. Turbidity assays were performed on a UV-Vis spectrophotometer (Thermo Scientific, Evolution 350 UV-Vis Spectrophotometer). Samples (in quartz cuvettes with 1 cm path length; Thorlabs) were first equilibrated, and the instrument was blanked. Throughout the experiment, absorbance was measured at $\lambda = 600$ nm. All samples were examined in triplicates ($n = 3$).

##### 4.13. Live cell imaging and Fluorescence Recovery After Photobleaching (FRAP)

FRAP was performed using an inverted confocal super-resolution microscope Zeiss LSM 880-Airyscan Elyra PS.1 with a 63x oil immersion objective. Cells were maintained in the microscope chamber at 37ºC and 5% $CO_2$. A circular region of interest (ROI) 23 mm was selected on GFP-PGC1 nuclear condensates and bleached with Argon laser at 488 nm wavelength by 10 consecutive bleaching iterations (1ms). Fluorescence intensity was recorded at baseline, during and after the bleaching and corrected by ROI corresponding to the background. Image processing and analysis was performed with ImageJ. Hoechst 33342 (Thermo Fisher) was used to stain the nucleus. Pictures of PGC1A droplets were taken and distance to nuclear lamina was assessed using imageJ. Mean distances in control were used for normalization when comparing different cellular interventions. Different media conditions were used as indicated; Dulbecco's without glucose (6-8h) for fasting condition or Dulbecco´s high glucose medium. When indicated, cells were transfected with targeted siRNA for knockdown experiments or treated with DGAT1 inhibitor. Treatment with free fatty acid palmitate excess (10uM, Sigma Aldrich), acylcarnitines as indicated, and 1,6 Hexanediol at 2.5mM was used as negative control (240117, sigma aldrich). 200mM KCL, HCL and NaOH were used to modify cellular pH.

##### 4.14. Genome editing of regulatory elements by CRISPR/Cas9

###### 4.14.A. Circuitry dissection

Our integrative genomic approach revealed chromatin segments with nonrandom higher-order locations. We have characterized their functional role in resource-conservation endurance across many experimental systems. We have focused on nuclear compartments with similar function, with functional interactions both at short- and long-range levels, along with their metabolic disease risk associations. The latter is observed in unbiased genomic range genetics, and in unbiased epigenomic integration of loci-trait-tissue relationships, where both underscored the importance of inter-trait-disease dependencies for metabolic-related traits, especially lipid associated modules. We next tested in polygenic risk scores and downstream causal genes, the topological relations between T2D and BMI variant-hubs, and how these colocalize with metabolic-HAR domains, interacting chromatin regions, cell-specific transcriptional hotspots, driver regions of metabolic disease, especially lipid disease, and within selected functional compartments. This approach showed functional nuclear compartments for lipid cycling genes, fine-tuned by cis-regulatory human accelerated regions, heterotypic condensates and local thermodynamic states, such as resource limitation. The same functional regions display strong topological relationships to T2D-BMI polygenic risk variant hubs, bridging segments of chromatin architecture and regions of inter-trait and inter-disease dependencies by lipid-related modules. Our experimental validation confirmed this, and helped find top functional dependencies during resource conservation, such as chr1-ATG4C region and chr8-DGAT1 region. During fuel restriction, we showed that HAR-enhancers are more sensitive to metabolic overload by acylcarnitines, while key structural enhancers (CTCF-bound) formed promoter loops that are resilient to acylcarnitine excess. Regarding the latter, even though they preserved basal transcriptional activity of target genes in starvation, fatty acid excess decreased their burst profile. For these reasons, we dissected the local circuitry of this heterotypic compartment using genome editing of key regulatory elements (HARs and structural) in human adipocytes, followed by functional, structural and transcriptional assessment of their genome perturbation in starvation states. Our initial goal is to test our previous findings and find causality as to how HARs offer stability to social genome plasticity, and how structural enhancers are key for intra-domain stability during resource conservation. Second, we want to find context-dependent causality for loss of genome plasticity transitions, and how these are relevant for phenotype outcomes of metabolic genetic variants within the circuitry. Thus, we focused on the local interacting circuitry between HAR elements and promoter regions of target genes, followed by isolation of key CTCF-bound enhancers (this is in addition to experimentally validated loops such as YY1-mediated), with genetic variations related to metabolic traits, especially lipid disorders (as described in the genomic-range genetics section).

##### 4.14.B. ATG4C domain circuitry

Following our validations, we focused on the HAR ANC162 that forms interactions with the promoter region. We carried out C-to-T genome editing using CRISPR/Cas9 in the binding motif (see transcriptional pipeline for TFBS-assessment) of a PGC1A-interacting target, YY1. As shown in our results in fig. S17 on circuit dissection, there is a genetic variation rs17124210 in that position, related to waist-hip-ratio (mutation name CRISPR-1a). Next, we focused on our previously validated key enhancer linked to ATG4C promoter, R5 (fig. S17). In glucose starvation, this enhancer forms YY1-loops, and it is bound by CTCF as well. Given our results on heterotypic cooperative behavior of interacting transcriptional regulators, and how CTCF as a chromatin binder participates in this plasticity, we focus on finding CTCF-binding sites with genetic variations or in proximity. Using the DNA sequence of the whole regulatory element span, we scanned it for CTCF-motifs using Ziebarth et al. approach (*9*). Considering the syntax effect and influence of neighboring motifs, we selected for genome editing the closest CTCF-motif to the strongest genetic variation to lipid disorders, rs12130333. This position is associated with the ANGPTL3 gene, which is located within the ATG4C submodule. In this motif, we also carried out C-to-T genome editing using CRISPR/Cas9 in the binding motif (mutation name CRISPR-1b).

##### 4.14.C. DGAT1 domain circuitry

We have shown in our experimental validations that DGAT1 location in the tail of chromosome 8 lies in a social interchromosomal segment, and belongs to the selected mHAR nuclear compartment. We found that, in glucose starvation, CTCF mediates long-range interactions (homotypic to CAT-domain and heterotypic to metabolic-HAR ATG4C domain). We used the network reconstruction of short-range interactions to obtain DGAT1 territorial module and submodules. We then focused on the HAR-HACNS71 element as a candidate for DGAT1-circuitry, which, due to distance constraints (1mbp), was validated as CTCF-mediated interaction. In close proximity to this HAR element, lies an active element_558 (VISTA), forming cis and trans interactions (bound by CTCF and YY1, both members of PGC1A-heterotypic compartments). Therefore, we mapped top CTCF-motifs in this HAR element to perform C-to-T genome editing using CRISPR/Cas9 (fig. S17; mutation name CRISPR-8a). Next, we scanned regulatory elements within the DGAT1 submodule for genetic variants associated with similar phenotypes to the ones in the trans-interacting ATG4C domain. We found in a regulatory cluster, a genetic variation rs55831924 associated with lipid disorders and related to the PLEC gene. Unbiased mapping of CTCF-motifs in the regulatory element revealed a genetic variant colocalized with a top CTCF-motif. This motif was selected for C-to-T genome editing using CRISPR/Cas9 (mutation name CRISPR-8b). Full network reconstruction of circuitry dissection can be found in supplementary figure S17.

##### 4.14.D. Genome editing

We performed genome editing in human adipose cells as previously described (*94*). In brief, all mutations in selected regulatory elements (some with genetic variations; see previous section) were from CC to TT alleles. hCas9 and guide RNA (gRNA) vectors were used (Addgene). C-to-T change was done using site-directed mutagenesis Q5 kit (New England Biolabs). Guide RNAs were designed using the CRISPR design tool available at http://crispr.mit.edu/. GFP,

Cas9 with sgRNAs, homology vector, and pMACs 4.1 plasmids were co-transfected in adipose progenitor cells using Amaxa-nucleofector (Lonza). Cell sorting was done using MACSelectTM (Miltenyi Biotec), followed by cell culture for 6 days, and propagation of selected clones for downstream experiments. These were functional assays (FFA, ROS, TG, pH, OCR), targeted chromatin interactions (cis and trans within circuitry), expression of target genes (ATG4C and DGAT1). For the last one, we were guided by the interaction frequencies among regulatory elements within the domain (defined in Fig 2). Network or topologically-based prioritization of interacting modules was used to identify regulatory regions with specific features such as the number of associations (see section integration of short-range chromatin relationships), followed by functional annotation, and detection of strong genetic variants in contiguous segments such as PCSK9 in Chr1:55mb (rs11591147). We then tested in cells with mutations in key structural enhancers (1b and 8b), if loss of genomic plasticity during nutrient limitation leads to transition of transcriptional activity to regulatory regions with less number of associations within the domains of interest (ATG4C-, DGAT1-, and PCSK9-domains). We focused on promoter regulatory regions with less number of associations and quantified their transcriptional activity in the aforementioned conditions. Primers are available upon reasonable request.

##### 4.15. Extracellular Flux Analysis (Seahorse) Assays

Cells were differentiated and treated as described above. When indicated, mitochondrial oxidative phosphorylation from human adipocytes was analyzed using extracellular flux analysis (XF24; Seahorse Biosciences) in DMEM buffer (pH 7.4; Sigma Aldrich). Baseline oxygen consumption rates (OCR) were measured every 7 min for control and cells undergoing glucose fasting as previously described.

##### 4.16. Intracellular pH measurement

Cellular models were maintained and treated as indicated (see mammalian cell culture). Cytosolic pH in different conditions were obtained with the intracellular pH indicator kit (pHrodo Red AM, fluorescent, Thermo fisher) following manufacturer's instructions and with a multimode plate reader (Agilent).

##### 4.17. Adenovirus-mediated gene delivery

Adenovirus expressing *Gfp* and wild-type or mutant PGC1A were generated by using the pAdTrack/pAdEasy system (Stratagene). Adenovirus-mediated gene delivery to mouse liver age-matched, mutant mice on a C57BL/6J background received 5 × 100 adenoviral particles in PBS into by tail vein injections (n = 5–6 per group). Five days post-transduction, mice were challenged with acute fasting interventions followed by metabolic and molecular phenotyping.

##### 4.18. Intraperitoneal glucose tolerance test (IP-GTT)

Intraperitoneal GTT was performed on overnight fasted mice. Blood glucose levels were measured at basal state (0 min) and then at 30, 60, and 120 min after i.p. injection of glucose (1.5 mg/kg body weight). Blood glucose concentrations were measured using the Accu-Chek Aviva monitoring system (Roche, Basel, Switzerland).

##### 4.19. Insulin tolerance test (ITT)

ITT was performed on fed mice. Blood glucose levels were measured at basal state (0 min) and then at 30, 60, and 120 min after i.p. injection of human insulin (1.0 mU/g body weight; Actrapid Penfill, Novo Nordisk, Denmark). Blood glucose concentrations were measured using the Accu-Chek Aviva monitoring system (Roche, Basel, Switzerland).

##### 4.20. Tissue pH

Measurement of tissue pH was performed as previously described (*95*). Briefly, mice were sacrificed by cervical dislocation, and brown adipose tissue, inguinal adipose tissue and liver were dissected and collected in independent samples. Tissue samples of relatively similar size were used. 50ul of purified water was added onto each sample, and mixed thoroughly. The pH value was assessed using a portable pH meter LAQUA with a 9618S-10D micro electrode (Horiba, Kyoto, Japan). All measurements were performed immediately after dissection.

##### 4.21. Intracellular pH measurement

Cellular models were maintained and treated as indicated (see mammalian cell culture). Cytosolic pH in different conditions were obtained with the intracellular pH indicator kit (pHrodo Red AM, fluorescent, Thermo fisher) following manufacturer's instructions and with a multimode plate reader (Agilent). 2

##### 4.22. Mitochondrial purification protocol

10 g of tissue were used from each animal for mitochondrial purification protocol. For in vitro experiments cells were collected by centrifugation at 400 xg for 10 min, resuspended in NKM buffer (1 mM Tris HCl pH 7.4, 0.13 M NaCl, 5 mM KCl, 7.5 mM MgCl2) and washed twice with this step. Cells were transferred into a glass homogenizer and incubated for 10 min on ice. Homogenize with a tight pestle with 30 strokes. Pour into a centrifuge tube containing 2 M sucrose solution, followed by mixing gently. Pellet unbroken cells at 1200 g for 5 min and transfer the supernatant to another tube. Repeat this twice and transfer the supernatant to a new tube. Centrifugation at 7000 xg for 10' at 4°C was performed to bring down mitochondria. We resuspend the pellet in mitochondrial suspension buffer (10 mM Tris HCl pH 6..7, 0.15 mM MgCl2, 0.25 mM sucrose, 1 mM PMSF, 1 mM DTT), followed by spinning down at 9500 xg for 5 min to re-pellet the mitochondria. Tissues were homogenized with Mitochondria Isolation buffer (225 mM Mannitol, 75 mM Sucrose, 30 mM Tris-HCl pH 7.4, 0,1mM EGTA). Tissues were slowly homogenized with strokes to break them up to 80%–90% in the homogenization buffer (10 mM Tris-HCl pH 6.7, 10 mM KCl, 0.15 mM MgCl2, 1 mM PMSF, 1 mM DTT). They were spun down at 600 xg for 10' to bring down unbroken cells and cell debris. The supernatant was then transferred to a previous weighted Eppendorf. Centrifugation at 7000 xg for 10' at 4°C was performed to bring down mitochondria. We resuspend the pellet in mitochondrial suspension buffer (10 mM Tris HCl pH 6..7, 0.15 mM MgCl2, 0.25 mM sucrose, 1 mM PMSF, 1 mM DTT), followed by spinning down at 9500 xg for 5 min to re-pellet the mitochondria. Samples were stored at -80 for further processing for example protein carbonylation assays, immunoblotting followed by carbonylation assay, or mitochondrial activity assays.

##### 4.23. Lysosome enrichment kit

For lysosome purification we used the lysosome enrichment kit for tissue and culture cells (Thermo Scientific, 89839) following manufacturer instruction protocols. Extraction of lysosome, when indicated, was followed up by protein carbonylation assay on whole extracts, or immunoblotting of specific protein with carbonylation assays, or by lysosome activity kits.

##### 4.24. Protein extracts

Cell cultures were homogenized in lysis buffer (50 mM Tris-HCl pH 7.4, 180 mM NaCl, 1% Triton X-100, 15% glycerol, and 1 mM EDTA) supplemented with 1 mM DTT, 0.5 mM PMSF and phosphatase inhibitor cocktail (Sigma-Aldrich). Protein concentration was determined by Bradford Protein Assay following manufacturer's instructions (BioRad).

##### 4.25. Data, Quantification, and Statistical Analysis

Statistical analyses related to experimental procedures were performed using GraphPad Prism Software (San Diego, CA) and all parameters are indicated in the corresponding figure legend. Quantitative data are presented as the mean ± SEM and n is indicated for each experiment. All experiments were carried out with 3 biological replicates and 3 independent times. Animal experiments were performed in more than 3 animals per group, both female and males were used when indicated. Unpaired Student's t test was used to determine statistical significance when two groups were compared. One-way ANOVA followed by Fisher's least significant difference (LSD) test for post hoc comparisons was used to determine statistical significance when multiple groups were compared. Two-way ANOVA was used to estimate significance between groups constrained by time-based or concentration-based measurements, followed by Tukey test for post-hoc multiple comparisons. Hypergeometric test enrichments and Chi-square tests were used to determine the statistical significance between expected and observed frequencies. Correlations were calculated by Spearman rank and Pearson correlation with a two-tailed test. Statistical significance was defined as $p$ value $< 0.05$ by either test and is denoted with an asterisk as follows: * p-value 0.05-0.01, ** p-value 0.01-0.001 and *** < 0.001 when indicated.

**Disclosure on the use of generative AI:** During the preparation of this work, the authors used the assistance of LLMs such as GPT5, Claude 4.5, Gemini 2.5 pro to edit and improve readability and grammar. All suggestions of text edits were edited, reviewed, validated, and manually corrected by the authors prior to submission of this work. The authors take full responsibility for the originality and content of this work. In addition, during preparation of specific parts, the authors used the same models for code assistance. The authors reviewed and edited the content as needed and take responsibility for the content of this work.

#### 5. References

1. H. Pontzer, M. H. Brown, D. A. Raichlen, H. Dunsworth, B. Hare, K. Walker, A. Luke, L. R. Dugas, R. Durazo-Arvizu, D. Schoeller, J. Plange-Rhule, P. Bovet, T. E. Forrester, E. V. Lambert, M. E. Thompson, R. W. Shumaker, S. R. Ross, Metabolic acceleration and the evolution of human brain size and life history. *Nature* **533**, 390–392 (2016).
2. H. Pontzer, D. A. Raichlen, A. D. Gordon, K. K. Schroepfer-Walker, B. Hare, M. C. O'Neill, K. M. Muldoon, H. M. Dunsworth, B. M. Wood, K. Isler, J. Burkart, M. Irwin, R. W. Shumaker, E. V. Lonsdorf, S. R. Ross, Primate energy expenditure and life history. *Proc. Natl. Acad. Sci. U. S. A.* **111**, 1433–1437 (2014).
3. A. D. Schmitt, M. Hu, I. Jung, Z. Xu, Y. Qiu, C. L. Tan, Y. Li, S. Lin, Y. Lin, C. L. Barr, B. Ren, A Compendium of Chromatin Contact Maps Reveals Spatially Active Regions in the Human Genome. *Cell Rep.* **17**, 2042–2059 (2016).
4. S. Kaufmann, C. Fuchs, M. Gonik, E. E. Khrameeva, A. A. Mironov, D. Frishman, Inter-chromosomal contact networks provide insights into Mammalian chromatin organization. *PLoS One* **10**, e0126125 (2015).
5. S. Fishilevich, R. Nudel, N. Rappaport, R. Hadar, I. Plaschkes, T. Iny Stein, N. Rosen, A. Kohn, M. Twik, M. Safran, D. Lancet, D. Cohen, GeneHancer: genome-wide integration of enhancers and target genes in GeneCards. *Database* **2017** (2017).
6. C. S. Greene, A. Krishnan, A. K. Wong, E. Ricciotti, R. A. Zelaya, D. S. Himmelstein, R. Zhang, B. M. Hartmann, E. Zaslavsky, S. C. Sealfon, D. I. Chasman, G. A. FitzGerald, K. Dolinski, T. Grosser, O. G. Troyanskaya, Understanding multicellular function and disease with human tissue-specific networks. *Nat. Genet.* **47**, 569–576 (2015).
7. D. Swain-Lenz, A. Berrio, A. Safi, G. E. Crawford, G. A. Wray, Comparative Analyses of Chromatin Landscape in White Adipose Tissue Suggest Humans May Have Less Beigeing Potential than Other Primates. *Genome Biol. Evol.* **11**, 1997–2008 (2019).
8. S. Berto, I. Mendizabal, N. Usui, K. Toriumi, P. Chatterjee, C. Douglas, C. A. Tamminga, T. M. Preuss, S. V. Yi, G. Konopka, Accelerated evolution of oligodendrocytes in the human brain. *Proc. Natl. Acad. Sci. U. S. A.* **116**, 24334–24342 (2019).
9. J. D. Ziebarth, A. Bhattacharya, Y. Cui, CTCFBSDB 2.0: a database for CTCF-binding sites and genome organization. *Nucleic Acids Res.* **41**, D188–94 (2013).
10. T. Misteli, The Self-Organizing Genome: Principles of Genome Architecture and Function. *Cell* **183**, 28–45 (2020).
11. C. H. Eskiw, N. F. Cope, I. Clay, S. Schoenfelder, T. Nagano, P. Fraser, Transcription factories and nuclear organization of the genome. *Cold Spring Harb. Symp. Quant. Biol.* **75**, 501–506 (2010).
12. J. Dekker, T. Misteli, Long-Range Chromatin Interactions. *Cold Spring Harb. Perspect. Biol.* **7**, a019356 (2015).
13. S. A. Quinodoz, N. Ollikainen, B. Tabak, A. Palla, J. M. Schmidt, E. Detmar, M. M. Lai, A. A. Shishkin, P. Bhat, Y. Takei, V. Trinh, E. Aznauryan, P. Russell, C. Cheng, M. Jovanovic, A. Chow, L. Cai, P. McDonel, M. Garber, M. Guttman, Higher-Order Inter-chromosomal Hubs Shape 3D Genome Organization in the Nucleus. *Cell* **174**, 744–757.e24 (2018).
14. M. R. Branco, A. Pombo, Intermingling of chromosome territories in interphase suggests role in translocations and transcription-dependent associations. *PLoS Biol.* **4**, e138 (2006).
15. A. Belyaeva, S. Venkatachalapathy, M. Nagarajan, G. V. Shivashankar, C. Uhler, Network analysis identifies chromosome intermingling regions as regulatory hotspots for transcription. *Proc. Natl. Acad. Sci. U. S. A.* **114**, 13714–13719 (2017).
16. G. van Mierlo, O. Pushkarev, J. F. Kribelbauer, B. Deplancke, Chromatin modules and their implication in genomic organization and gene regulation. *Trends Genet.* **39**, 140–153 (2023).
17. R. Fang, M. Yu, G. Li, S. Chee, T. Liu, A. D. Schmitt, B. Ren, Mapping of long-range chromatin interactions by proximity ligation-assisted ChIP-seq. *Cell Res.* **26**, 1345–1348 (2016).
18. E. Lieberman-Aiden, N. L. van Berkum, L. Williams, M. Imakaev, T. Ragoczy, A. Telling, I. Amit, B. R. Lajoie, P. J. Sabo, M. O. Dorschner, R. Sandstrom, B. Bernstein, M. A. Bender, M. Groudine, A. Gnirke, J. Stamatoyannopoulos, L. A. Mirny, E. S. Lander, J. Dekker, Comprehensive mapping of long-range interactions reveals folding principles of the human genome. *Science* **326**, 289–293 (2009).
19. G. Cavalli, T. Misteli, Functional implications of genome topology. *Nat. Struct. Mol. Biol.* **20**, 290–299 (2013).
20. K. S. Pollard, S. R. Salama, N. Lambert, M.-A. Lambot, S. Coppens, J. S. Pedersen, S. Katzman, B. King, C. Onodera, A. Siepel, A. D. Kern, C. Dehay, H. Igel, M. Ares Jr, P. Vanderhaeghen, D. Haussler, An RNA gene expressed during cortical development evolved rapidly in humans. *Nature* **443**, 167–172 (2006).
21. J. A. Capra, G. D. Erwin, G. McKinsey, J. L. R. Rubenstein, K. S. Pollard, Many human accelerated regions are developmental enhancers. *Philos. Trans. R. Soc. Lond. B Biol. Sci.* **368**, 20130025 (2013).
22. A. Navarrete, C. P. van Schaik, K. Isler, Energetics and the evolution of human brain size. *Nature* **480**, 91–93 (2011).
23. S. L. Lindstedt, M. S. Boyce, Seasonality, Fasting Endurance, and Body Size in Mammals. [Preprint] (1985). https://doi.org/10.1086/284385.
24. M. V. Kuleshov, M. R. Jones, A. D. Rouillard, N. F. Fernandez, Q. Duan, Z. Wang, S. Koplev, S. L. Jenkins, K. M. Jagodnik, A. Lachmann, M. G. McDermott, C. D. Monteiro, G. W. Gundersen, A. Ma'ayan, Enrichr: a comprehensive gene set enrichment analysis web server 2016 update. *Nucleic Acids Res.* **44**, W90–7 (2016).
25. K. Luck, D.-K. Kim, L. Lambourne, K. Spirohn, B. E. Begg, W. Bian, R. Brignall, T. Cafarelli, F. J. Campos-Laborie, B. Charloteaux, D. Choi, A. G. Coté, M. Daley, S. Deimling, A. Desbuleux, A. Dricot, M. Gebbia, M. F. Hardy, N. Kishore, J. J. Knapp, I. A. Kovács, I. Lemmens, M. W. Mee, J. C. Mellor, C. Pollis, C. Pons, A. D. Richardson, S. Schlabach, B. Teeking, A. Yadav, M. Babor, D. Balcha, O. Basha, C. Bowman-Colin, S.-F. Chin, S. G. Choi, C. Colabella, G. Coppin, C. D'Amata, D. De Ridder, S. De Rouck, M. Duran-Frigola, H. Ennajdaoui, F. Goebels, L. Goehring, A. Gopal, G. Haddad, E. Hatchi, M. Helmy, Y. Jacob, Y. Kassa, S. Landini, R. Li, N. van Lieshout, A. MacWilliams, D. Markey, J. N. Paulson, S. Rangarajan, J. Rasla, A. Rayhan, T. Rolland, A. San-Miguel, Y. Shen, D. Sheykhkarimli, G. M. Sheynkman, E. Simonovsky, M. Taşan, A. Tejeda, V. Tropepe, J.-C. Twizere, Y. Wang, R. J. Weatheritt, J. Weile, Y. Xia, X. Yang, E. Yeger-Lotem, Q. Zhong, P. Aloy, G. D. Bader, J. De Las Rivas, S. Gaudet, T. Hao, J. Rak, J. Tavernier, D. E. Hill, M. Vidal, F. P. Roth, M. A. Calderwood, A reference map of the human binary protein interactome. *Nature* **580**, 402–408 (2020).
26. R. Oughtred, J. Rust, C. Chang, B.-J. Breitkreutz, C. Stark, A. Willems, L. Boucher, G. Leung, N. Kolas, F. Zhang, S. Dolma, J. Coulombe-Huntington, A. Chatr-Aryamontri, K. Dolinski, M. Tyers, The BioGRID database: A comprehensive biomedical resource of curated protein, genetic, and chemical interactions. *Protein Sci.* **30**, 187–200 (2021).
27. D. W. Huang, B. T. Sherman, R. A. Lempicki, Systematic and integrative analysis of large gene lists using DAVID bioinformatics resources. *Nat. Protoc.* **4**, 44–57 (2009).
28. S. Oki, T. Ohta, G. Shioi, H. Hatanaka, O. Ogasawara, Y. Okuda, H. Kawaji, R. Nakaki, J. Sese, C. Meno, ChIP-Atlas: a data-mining suite powered by full integration of public ChIP-seq data. *EMBO Rep.* **19** (2018).
29. Z. Obradovic, K. Peng, S. Vucetic, P. Radivojac, C. J. Brown, A. K. Dunker, Predicting intrinsic disorder from amino acid sequence. *Proteins* **53 Suppl 6**, 566–572 (2003).
30. R. Pancsa, W. Vranken, B. Mészáros, Computational resources for identifying and describing proteins driving liquid-liquid phase separation. *Brief. Bioinform.*, doi: 10.1093/bib/bbaa408 (2021).
31. Z. Dosztányi, B. Mészáros, I. Simon, ANCHOR: web server for predicting protein binding regions in disordered proteins. *Bioinformatics* **25**, 2745–2746 (2009).
32. R. M. Vernon, P. A. Chong, B. Tsang, T. H. Kim, A. Bah, P. Farber, H. Lin, J. D. Forman-Kay, Pi-Pi contacts are an overlooked protein

feature relevant to phase separation. *Elife* **7** (2018).
33. K. You, Q. Huang, C. Yu, B. Shen, C. Sevilla, M. Shi, H. Hermjakob, Y. Chen, T. Li, PhaSepDB: a database of liquid-liquid phase separation related proteins. *Nucleic Acids Res.* **48**, D354–D359 (2020).
34. V. N. Uversky, J. R. Gillespie, A. L. Fink, Why are “natively unfolded” proteins unstructured under physiologic conditions? *Proteins* **41**, 415–427 (2000).
35. A. S. Holehouse, J. Ahad, R. K. Das, R. V. Pappu, CIDER: Classification of Intrinsically Disordered Ensemble Regions. [Preprint] (2015). https://doi.org/10.1016/j.bpj.2014.11.1260.
36. R. K. Das, R. V. Pappu, Conformations of intrinsically disordered proteins are influenced by linear sequence distributions of oppositely charged residues. *Proc. Natl. Acad. Sci. U. S. A.* **110**, 13392–13397 (2013).
37. J. Kyte, R. F. Doolittle, A simple method for displaying the hydropathic character of a protein. *J. Mol. Biol.* **157**, 105–132 (1982).
38. H. Ashkenazy, S. Abadi, E. Martz, O. Chay, I. Mayrose, T. Pupko, N. Ben-Tal, ConSurf 2016: an improved methodology to estimate and visualize evolutionary conservation in macromolecules. *Nucleic Acids Res.* **44**, W344–50 (2016).
39. T. Pupko, I. Pe’er, M. Hasegawa, D. Graur, N. Friedman, A branch-and-bound algorithm for the inference of ancestral amino-acid sequences when the replacement rate varies among sites: Application to the evolution of five gene families. *Bioinformatics* **18**, 1116–1123 (2002).
40. A. Biegert, J. Söding, Sequence context-specific profiles for homology searching. *Proc. Natl. Acad. Sci. U. S. A.* **106**, 3770–3775 (2009).
41. I. Mayrose, D. Graur, N. Ben-Tal, T. Pupko, Comparison of site-specific rate-inference methods for protein sequences: empirical Bayesian methods are superior. *Mol. Biol. Evol.* **21**, 1781–1791 (2004).
42. P. W. Rose, A. Prlić, C. Bi, W. F. Bluhm, C. H. Christie, S. Dutta, R. K. Green, D. S. Goodsell, J. D. Westbrook, J. Woo, J. Young, C. Zardecki, H. M. Berman, P. E. Bourne, S. K. Burley, The RCSB Protein Data Bank: views of structural biology for basic and applied research and education. *Nucleic Acids Res.* **43**, D345–56 (2015).
43. M. Remmert, A. Biegert, A. Hauser, J. Söding, HHblits: lightning-fast iterative protein sequence searching by HMM-HMM alignment. *Nat. Methods* **9**, 173–175 (2011).
44. A. Veidenberg, A. Medlar, A. Löytynoja, Wasabi: An Integrated Platform for Evolutionary Sequence Analysis and Data Visualization. *Mol. Biol. Evol.* **33**, 1126–1130 (2016).
45. X. Ji, D. B. Dadon, B. J. Abraham, T. I. Lee, R. Jaenisch, J. E. Bradner, R. A. Young, Chromatin proteomic profiling reveals novel proteins associated with histone-marked genomic regions. *Proc. Natl. Acad. Sci. U. S. A.* **112**, 3841–3846 (2015).
46. A. S. Weintraub, C. H. Li, A. V. Zamudio, A. A. Sigova, N. M. Hannett, D. S. Day, B. J. Abraham, M. A. Cohen, B. Nabet, D. L. Buckley, Y. E. Guo, D. Hnisz, R. Jaenisch, J. E. Bradner, N. S. Gray, R. A. Young, YY1 Is a Structural Regulator of Enhancer-Promoter Loops. *Cell* **171**, 1573–1588.e28 (2017).
47. M. P. Creyghton, A. W. Cheng, G. G. Welstead, T. Kooistra, B. W. Carey, E. J. Steine, J. Hanna, M. A. Lodato, G. M. Frampton, P. A. Sharp, L. A. Boyer, R. A. Young, R. Jaenisch, Histone H3K27ac separates active from poised enhancers and predicts developmental state. *Proc. Natl. Acad. Sci. U. S. A.* **107**, 21931–21936 (2010).
48. K. De Preter, R. Barriot, F. Speleman, J. Vandesompele, Y. Moreau, Positional gene enrichment analysis of gene sets for high-resolution identification of overrepresented chromosomal regions. *Nucleic Acids Res.* **36**, e43 (2008).
49. J. R. Dixon, D. U. Gorkin, B. Ren, Chromatin Domains: The Unit of Chromosome Organization. *Mol. Cell* **62**, 668–680 (2016).
50. S. S. P. Rao, M. H. Huntley, N. C. Durand, E. K. Stamenova, I. D. Bochkov, J. T. Robinson, A. L. Sanborn, I. Machol, A. D. Omer, E. S. Lander, E. L. Aiden, A 3D map of the human genome at kilobase resolution reveals principles of chromatin looping. *Cell* **159**, 1665–1680 (2014).
51. N. C. Durand, J. T. Robinson, M. S. Shamim, I. Machol, J. P. Mesirov, E. S. Lander, E. L. Aiden, Juicebox Provides a Visualization System for Hi-C Contact Maps with Unlimited Zoom. *Cell Syst* **3**, 99–101 (2016).
52. E. Crane, Q. Bian, R. P. McCord, B. R. Lajoie, B. S. Wheeler, E. J. Ralston, S. Uzawa, J. Dekker, B. J. Meyer, Condensin-driven remodelling of X chromosome topology during dosage compensation. *Nature* **523**, 240–244 (2015).
53. F. Ay, T. L. Bailey, W. S. Noble, Statistical confidence estimation for Hi-C data reveals regulatory chromatin contacts. *Genome Res.* **24**, 999–1011 (2014).
54. M. Hu, K. Deng, S. Selvaraj, Z. Qin, B. Ren, J. S. Liu, HiCNorm: removing biases in Hi-C data via Poisson regression. *Bioinformatics* **28**, 3131–3133 (2012).
55. M. Imakaev, G. Fudenberg, R. P. McCord, N. Naumova, A. Goloborodko, B. R. Lajoie, J. Dekker, L. A. Mirny, Iterative correction of Hi-C data reveals hallmarks of chromosome organization. *Nat. Methods* **9**, 999–1003 (2012).
56. J. R. Dixon, S. Selvaraj, F. Yue, A. Kim, Y. Li, Y. Shen, M. Hu, J. S. Liu, B. Ren, Topological domains in mammalian genomes identified by analysis of chromatin interactions. *Nature* **485**, 376–380 (2012).
57. E. Yaffe, A. Tanay, Probabilistic modeling of Hi-C contact maps eliminates systematic biases to characterize global chromosomal architecture. *Nat. Genet.* **43**, 1059–1065 (2011).
58. K. Kruse, S. Sewitz, M. M. Babu, A complex network framework for unbiased statistical analyses of DNA-DNA contact maps. *Nucleic Acids Res.* **41**, 701–710 (2013).
59. R. Andersson, C. Gebhard, I. Miguel-Escalada, I. Hoof, J. Bornholdt, M. Boyd, Y. Chen, X. Zhao, C. Schmidl, T. Suzuki, E. Ntini, E. Arner, E. Valen, K. Li, L. Schwarzfischer, D. Glatz, J. Raithel, B. Lilje, N. Rapin, F. O. Bagger, M. Jørgensen, P. R. Andersen, N. Bertin, O. Rackham, A. M. Burroughs, J. K. Baillie, Y. Ishizu, Y. Shimizu, E. Furuhata, S. Maeda, Y. Negishi, C. J. Mungall, T. F. Meehan, T. Lassmann, M. Itoh, H. Kawaji, N. Kondo, J. Kawai, A. Lennartsson, C. O. Daub, P. Heutink, D. A. Hume, T. H. Jensen, H. Suzuki, Y. Hayashizaki, F. Müller, A. R. R. Forrest, P. Carninci, M. Rehli, A. Sandelin, An atlas of active enhancers across human cell types and tissues. *Nature* **507**, 455–461 (2014).
60. H.-J. Westra, M. J. Peters, T. Esko, H. Yaghootkar, C. Schurmann, J. Kettunen, M. W. Christiansen, B. P. Fairfax, K. Schramm, J. E. Powell, A. Zhernakova, D. V. Zhernakova, J. H. Veldink, L. H. Van den Berg, J. Karjalainen, S. Withoff, A. G. Uitterlinden, A. Hofman, F. Rivadeneira, P. A. C. ’t Hoen, E. Reinmaa, K. Fischer, M. Nelis, L. Milani, D. Melzer, L. Ferrucci, A. B. Singleton, D. G. Hernandez, M. A. Nalls, G. Homuth, M. Nauck, D. Radke, U. Völker, M. Perola, V. Salomaa, J. Brody, A. Suchy-Dicey, S. A. Gharib, D. A. Enquobahrie, T. Lumley, G. W. Montgomery, S. Makino, H. Prokisch, C. Herder, M. Roden, H. Grallert, T. Meitinger, K. Strauch, Y. Li, R. C. Jansen, P. M. Visscher, J. C. Knight, B. M. Psaty, S. Ripatti, A. Teumer, T. M. Frayling, A. Metspalu, J. B. J. van Meurs, L. Franke, Systematic identification of trans eQTLs as putative drivers of known disease associations. *Nat. Genet.* **45**, 1238–1243 (2013).
61. B. Mifsud, F. Tavares-Cadete, A. N. Young, R. Sugar, S. Schoenfelder, L. Ferreira, S. W. Wingett, S. Andrews, W. Grey, P. A. Ewels, B. Herman, S. Happe, A. Higgs, E. LeProust, G. A. Follows, P. Fraser, N. M. Luscombe, C. S. Osborne, Mapping long-range promoter contacts in human cells with high-resolution capture Hi-C. *Nat. Genet.* **47**, 598–606 (2015).
62. A. Visel, S. Minovitsky, I. Dubchak, L. A. Pennacchio, VISTA Enhancer Browser--a database of tissue-specific human enhancers. *Nucleic Acids Res.* **35**, D88–92 (2007).
63. W. J. Shim, E. Sinniah, J. Xu, B. Vitrinel, M. Alexanian, G. Andreoletti, S. Shen, Y. Sun, B. Balderson, C. Boix, G. Peng, N. Jing, Y. Wang, M. Kellis, P. P. L. Tam, A. Smith, M. Piper, L. Christiaen, Q. Nguyen, M. Bodén, N. J. Palpant, Conserved Epigenetic Regulatory Logic Infers Genes Governing Cell Identity. *Cell Syst* **11**, 625–639.e13 (2020).
64. O. Oluwadare, Y. Zhang, J. Cheng, A maximum likelihood algorithm for reconstructing 3D structures of human chromosomes from

chromosomal contact data. *BMC Genomics* **19**, 161 (2018).
65. O. Oluwadare, M. Highsmith, D. Turner, E. Lieberman Aiden, J. Cheng, GSDB: a database of 3D chromosome and genome structures reconstructed from Hi-C data. *BMC Mol Cell Biol* **21**, 60 (2020).
66. M. Wlasnowolski, M. Sadowski, T. Czarnota, K. Jodkowska, P. Szalaj, Z. Tang, Y. Ruan, D. Plewczynski, 3D-GNOME 2.0: a three-dimensional genome modeling engine for predicting structural variation-driven alterations of chromatin spatial structure in the human genome. *Nucleic Acids Res.* **48**, W170–W176 (2020).
67. T. Trieu, J. Cheng, 3D genome structure modeling by Lorentzian objective function. *Nucleic Acids Res.* **45**, 1049–1058 (2017).
68. J. Wang, X. Dai, L. D. Berry, J. D. Cogan, Q. Liu, Y. Shyr, HACER: an atlas of human active enhancers to interpret regulatory variants. *Nucleic Acids Res.* **47**, D106–D112 (2019).
69. D. R. Zerbino, S. P. Wilder, N. Johnson, T. Juettemann, P. R. Flicek, The ensembl regulatory build. *Genome Biol.* **16**, 56 (2015).
70. J. Ernst, M. Kellis, Chromatin-state discovery and genome annotation with ChromHMM. *Nat. Protoc.* **12**, 2478–2492 (2017).
71. L. Teng, B. He, J. Wang, K. Tan, 4DGenome: a comprehensive database of chromatin interactions. *Bioinformatics* **32**, 2727 (2016).
72. Z. Tang, O. J. Luo, X. Li, M. Zheng, J. J. Zhu, P. Szalaj, P. Trzaskoma, A. Magalska, J. Wlodarczyk, B. Ruszczycki, P. Michalski, E. Piecuch, P. Wang, D. Wang, S. Z. Tian, M. Penrad-Mobayed, L. M. Sachs, X. Ruan, C.-L. Wei, E. T. Liu, G. M. Wilczynski, D. Plewczynski, G. Li, Y. Ruan, CTCF-Mediated Human 3D Genome Architecture Reveals Chromatin Topology for Transcription. *Cell* **163**, 1611–1627 (2015).
73. G. Li, X. Ruan, R. K. Auerbach, K. S. Sandhu, M. Zheng, P. Wang, H. M. Poh, Y. Goh, J. Lim, J. Zhang, H. S. Sim, S. Q. Peh, F. H. Mulawadi, C. T. Ong, Y. L. Orlov, S. Hong, Z. Zhang, S. Landt, D. Raha, G. Euskirchen, C.-L. Wei, W. Ge, H. Wang, C. Davis, K. I. Fisher-Aylor, A. Mortazavi, M. Gerstein, T. Gingeras, B. Wold, Y. Sun, M. J. Fullwood, E. Cheung, E. Liu, W.-K. Sung, M. Snyder, Y. Ruan, Extensive promoter-centered chromatin interactions provide a topological basis for transcription regulation. *Cell* **148**, 84–98 (2012).
74. J. MacArthur, E. Bowler, M. Cerezo, L. Gil, P. Hall, E. Hastings, H. Junkins, A. McMahon, A. Milano, J. Morales, Z. M. Pendlington, D. Welter, T. Burdett, L. Hindorff, P. Flicek, F. Cunningham, H. Parkinson, The new NHGRI-EBI Catalog of published genome-wide association studies (GWAS Catalog). *Nucleic Acids Res.* **45**, D896–D901 (2017).
75. T. Gao, J. Qian, EnhancerAtlas 2.0: an updated resource with enhancer annotation in 586 tissue/cell types across nine species. *Nucleic Acids Res.* **48**, D58–D64 (2020).
76. T. Gao, B. He, S. Liu, H. Zhu, K. Tan, J. Qian, EnhancerAtlas: a resource for enhancer annotation and analysis in 105 human cell/tissue types. *Bioinformatics* **32**, 3543–3551 (2016).
77. Y. Yang, J. Fear, J. Hu, I. Haecker, L. Zhou, R. Renne, D. Bloom, L. M. McIntyre, Leveraging biological replicates to improve analysis in ChIP-seq experiments. *Comput. Struct. Biotechnol. J.* **9**, e201401002 (2014).
78. C. A. Boix, B. T. James, Y. P. Park, W. Meuleman, M. Kellis, Regulatory genomic circuitry of human disease loci by integrative epigenomics. *Nature* **590**, 300–307 (2021).
79. C. J. Willer, Y. Li, G. R. Abecasis, METAL: fast and efficient meta-analysis of genomewide association scans. *Bioinformatics* **26**, 2190–2191 (2010).
80. M. L. Heltberg, S. Krishna, M. H. Jensen, On chaotic dynamics in transcription factors and the associated effects in differential gene regulation. *Nat. Commun.* **10**, 71 (2019).
81. A. J. M. Larsson, P. Johnsson, M. Hagemann-Jensen, L. Hartmanis, O. R. Faridani, B. Reinius, Å. Segerstolpe, C. M. Rivera, B. Ren, R. Sandberg, Genomic encoding of transcriptional burst kinetics. *Nature* **565**, 251–254 (2019).
82. H. J. Motulsky, R. E. Brown, Detecting outliers when fitting data with nonlinear regression - a new method based on robust nonlinear regression and the false discovery rate. *BMC Bioinformatics* **7**, 123 (2006).
83. J. Lee, W.-Y. Hong, M. Cho, M. Sim, D. Lee, Y. Ko, J. Kim, Synteny Portal: a web-based application portal for synteny block analysis. *Nucleic Acids Res.* **44**, W35–40 (2016).
84. R. T. Pillich, J. Chen, V. Rynkov, D. Welker, D. Pratt, NDEx: A Community Resource for Sharing and Publishing of Biological Networks. *Methods Mol. Biol.* **1558**, 271–301 (2017).
85. S. Kleiner, R. J. Mepani, D. Laznik, L. Ye, M. J. Jurczak, F. R. Jornayvaz, J. L. Estall, D. Chatterjee Bhowmick, G. I. Shulman, B. M. Spiegelman, Development of insulin resistance in mice lacking PGC-1α in adipose tissues. *Proc. Natl. Acad. Sci. U. S. A.* **109**, 9635–9640 (2012).
86. J. Wang, J. Wang, L. Yang, C. Zhao, L. N. Wu, L. Xu, F. Zhang, Q. Weng, M. Wegner, Q. R. Lu, CTCF-mediated chromatin looping in EGR2 regulation and SUZ12 recruitment critical for peripheral myelination and repair. *Nat. Commun.* **11**, 4133 (2020).
87. D. M. Valenzuela, A. J. Murphy, D. Frendewey, N. W. Gale, A. N. Economides, W. Auerbach, W. T. Poueymirou, N. C. Adams, J. Rojas, J. Yasenchak, R. Chernomorsky, M. Boucher, A. L. Elsasser, L. Esau, J. Zheng, J. A. Griffiths, X. Wang, H. Su, Y. Xue, M. G. Dominguez, I. Noguera, R. Torres, L. E. Macdonald, A. F. Stewart, T. M. DeChiara, G. D. Yancopoulos, High-throughput engineering of the mouse genome coupled with high-resolution expression analysis. *Nat. Biotechnol.* **21**, 652–659 (2003).
88. S. Kajimura, P. Seale, K. Kubota, E. Lunsford, J. V. Frangioni, S. P. Gygi, B. M. Spiegelman, Initiation of myoblast to brown fat switch by a PRDM16-C/EBP-beta transcriptional complex. *Nature* **460**, 1154–1158 (2009).
89. A. Galmozzi, B. P. Kok, E. Saez, Isolation and Differentiation of Primary White and Brown Preadipocytes from Newborn Mice. [Preprint] (2021). https://doi.org/10.3791/62005.
90. W. Gao, X. Kong, Q. Yang, Isolation, Primary Culture, and Differentiation of Preadipocytes from Mouse Brown Adipose Tissue. [Preprint] (2017). https://doi.org/10.1007/978-1-4939-6820-6_1.
91. J. L. Estall, J. L. Ruas, C. S. Choi, D. Laznik, M. Badman, E. Maratos-Flier, G. I. Shulman, B. M. Spiegelman, PGC-1alpha negatively regulates hepatic FGF21 expression by modulating the heme/Rev-Erb(alpha) axis. *Proc. Natl. Acad. Sci. U. S. A.* **106**, 22510–22515 (2009).
92. J. L. Ruas, J. P. White, R. R. Rao, S. Kleiner, K. T. Brannan, B. C. Harrison, N. P. Greene, J. Wu, J. L. Estall, B. A. Irving, I. R. Lanza, K. A. Rasbach, M. Okutsu, K. S. Nair, Z. Yan, L. A. Leinwand, B. M. Spiegelman, A PGC-1α isoform induced by resistance training regulates skeletal muscle hypertrophy. *Cell* **151**, 1319–1331 (2012).
93. K. Monahan, A. Horta, S. Lomvardas, LHX2- and LDB1-mediated trans interactions regulate olfactory receptor choice. *Nature* **565**, 448–453 (2019).
94. M. Claussnitzer, S. N. Dankel, K.-H. Kim, G. Quon, W. Meuleman, C. Haugen, V. Glunk, I. S. Sousa, J. L. Beaudry, V. Puviindran, N. A. Abdennur, J. Liu, P.-A. Svensson, Y.-H. Hsu, D. J. Drucker, G. Mellgren, C.-C. Hui, H. Hauner, M. Kellis, FTO Obesity Variant Circuitry and Adipocyte Browning in Humans. *N. Engl. J. Med.* **373**, 895–907 (2015).
95. K. Shimizu, I. Seiki, Y. Goto, T. Murata, Measurement of the Intestinal pH in Mice under Various Conditions Reveals Alkalization Induced by Antibiotics. *Antibiotics (Basel)* **10** (2021).
